# A genome resource for the marine annelid *Platynereis dumerilii*

**DOI:** 10.1101/2024.06.21.600153

**Authors:** Kevin Nzumbi Mutemi, Oleg Simakov, Leslie Pan, Luca Santangeli, Ryan Null, Mette Handberg-Thorsager, Bruno Cossermelli Vellutini, Kevin J. Peterson, Bastian Fromm, Tomas Larsson, Emily Savage, Mireia Osuna Lopez, Rajna Hercog, Jan Provaznik, Diana Ordoñez-Rueda, Nayara Azevedo, Eve Gazave, Michel Vervoort, Pavel Tomancak, Wenhua Tan, Sylke Winkler, Vladimir Benes, Jerome Hui, Conrad Helm, B. Duygu Özpolat, Detlev Arendt

## Abstract

The marine annelid *Platynereis dumerilii* is a model organism used in many research areas including evolution and development, neurobiology, ecology and regeneration. Here we present the genomes of *P. dumerilii* (laboratory culture reference and a single individual assembly) and of the closely related *P. massiliensis* and *P. megalops* (single individual assembly) to facilitate comparative genomic approaches and help explore *Platynereis* biology. We used long-read sequencing technology and chromosomal-conformation capture along with extensive transcriptomic resources to obtain and annotate a draft genome assembly of ∼1.47 Gbp for *P. dumerilii*, of which more than half represent repeat elements. We predict around 29,000 protein-coding genes, with relatively large intron sizes, over 38,000 non-coding genes, and 105 miRNA loci. We further explore the high genetic variation (∼3% heterozygosity) within the *Platynereis* species complex. Gene ontology reveals the most variable loci to be associated with pigmentation, development and immunity. The current work sets the stage for further development of *Platynereis* genomic resources.

## Introduction

*Platynereis dumerilii* (Audouin & Milne Edwards, 1833) is found along European coastlines, including from the Azores, Mediterranean, North, Black and Red Seas, English Channel, as well as Atlantic and Pacific, Sea of Japan and Persian Gulf (Fauvel, 1914; Teixeira et al., 2022). As a laboratory model species, it has been extensively studied in disciplines spanning ecology, behavior, physiology, development, evolution, regeneration and neurobiology (Fischer et al., 2010; Fischer & Dorresteijn, 2004; Özpolat et al., 2021). However, an annotated genome has been challenging to achieve, due to high polymorphism, heterozygosity and repetitive content (Raible et al., 2005; Zantke et al., 2014). Here, we report the generation of a comprehensive genome resource for *P. dumerilii*, which also builds on previously published extensive transcriptomic data (Chou et al., 2018; Conzelmann et al., 2013; Paré et al., 2023; Raible et al., 2005; Zantke et al., 2014). Utilizing long-read sequencing, dense transcriptomic sampling and high-throughput sequencing of chromosome conformation capture (Hi-C) data (Belton et al., 2012), we present a draft assembly and annotation of the *P. dumerilii* genome measuring ∼1.47 giga-base pairs (Gbp) in size, which is larger than other annelid genomes sequenced and annotated to date (Martín-Durán et al., 2021; Martín-Zamora et al., 2023; Shao et al., 2020; Simakov et al., 2013; Zakas et al., 2022) (Fig. 1). We find that at least 51% of the genome consists of repetitive regions, and model around 29,000 protein-coding genes, with median intron sizes of ∼1.3 kilo-base pairs (Kbp). This suggests that an increase in repeat content, intron sizes as well as modest gene duplication contributed to genome expansion. We explore evolutionary trends in genome organization, ancestral linkage groups, and gene content across Metazoa, focusing on comparisons across annelids and within the *Platynereis* genus. We also describe natural polymorphisms associated with habitats that *Platynereis* occupies globally. Finally, we present draft genomes of two sister species, *P. massiliensis* and *P. megalops,* which appear morphologically indistinguishable from *P. dumerilii* as juveniles, but present different reproductive, larval, and behavioral patterns.

**Figure 1.**
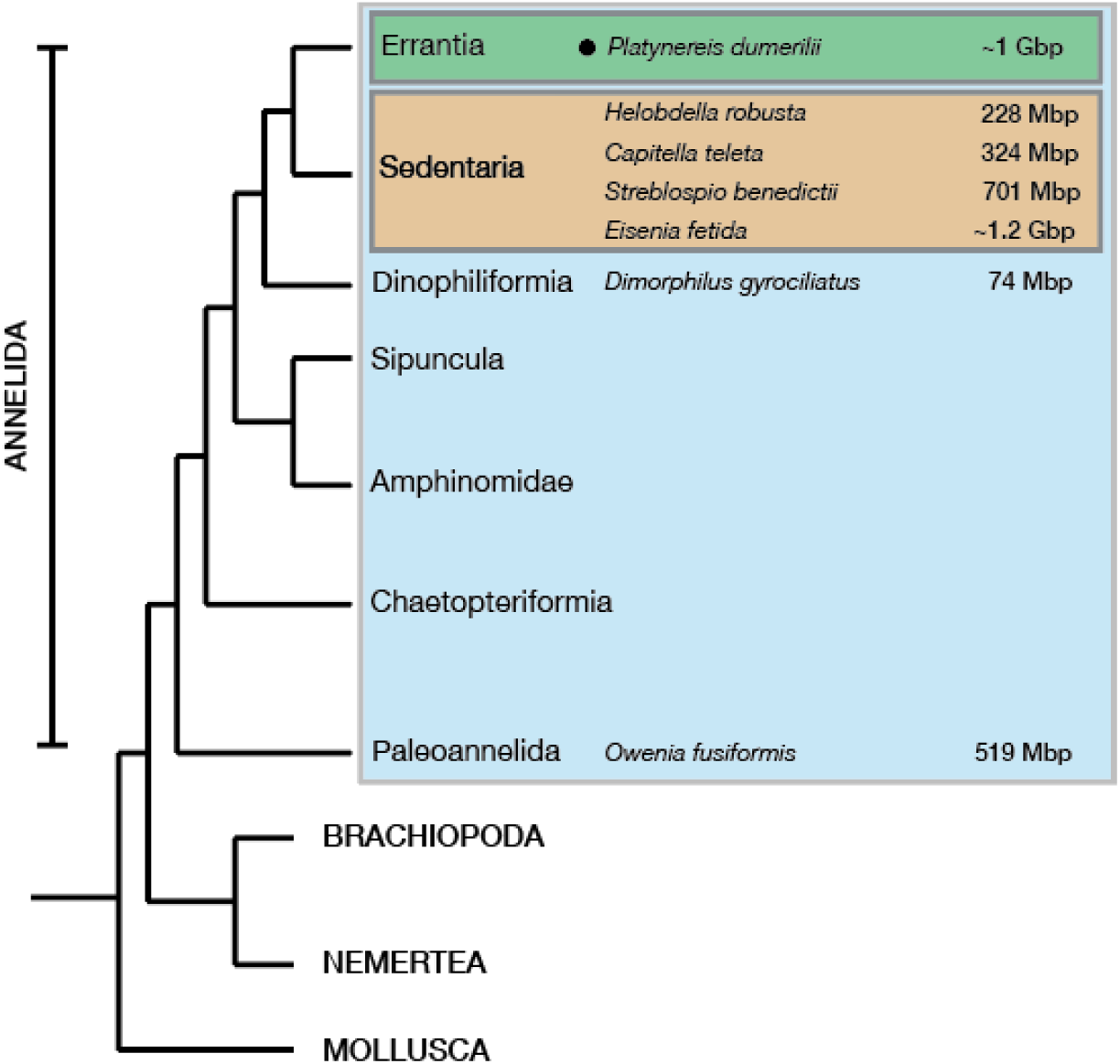
*Platynereis dumerilii* genome within Spiralia. A phylogenetic tree of major Spiralia/Lophotrochozoa groups, with the sequenced and ‘annotated’ annelid genome size estimates highlighted. The genome sizes are based off genome assemblies or DNA nuclei staining methods. The *Platynereis dumerilii* genome size numbers from (Jha et al., 1995), *Helobdella robusta* and *Capitella teleta* values were taken from (Simakov et al., 2013), the *Eisenia fetida* genome size estimates were taken from (Bhambri et al., 2018; Zwarycz et al., 2015), the *Streblospio benedictii* measurements were taken from (Zakas et al., 2022), the *Dimorphilus gyrociliatus* genome size from (Martín-Durán et al., 2021) and the *Owenia fusiformis*’ from (Liang et al., 2022).

### Results and Discussion Towards a chromosome-level assembly of the *P. dumerilii* genome

We sequenced high molecular weight genomic DNA (Supplementary Figs. 1-2), accessed from hundreds of progeny obtained from a single cross (one male with one female), combined with an additional single sexually mature male sample, with all samples coming from cultures bred in laboratory conditions since the 1960s (Fischer & Dorresteijn, 2004). We utilized PacBio Sequel II continuous long read (CLR) technology, sequencing at 200GB (corresponding roughly to 200x coverage, assuming ∼1 Gbp genome size (Jha et al., 1995) (Materials and Methods; Supplementary Fig. 1). Read lengths ranged from 50 – 270,934 bp with a median of 48,761 bp (mean ± standard deviation [SD] = 49,814.5 ± 30,070.61 bp) (Supplementary Fig. 2). From these reads, we assembled the genome using CANU (Koren et al., 2017), yielding a ∼3.41 Gbp assembly size with 9,431 contigs, and an N50 of 640 Kbp (Methods). At this state however, the genome size amounted to more than three times its previously predicted size (Jha et al., 1995). We re-evaluated the genome size, as well as ploidy, using GenomeScope2.0 and Smudgeplot (Ranallo-Benavidez et al., 2020) from high quality 150bp Illumina sequencing reads as well as DNA content quantified via flow cytometry. While these results estimated a genome size of ∼940 Mbp (Supplementary Fig. 3), flow cytometry measurements suggested a genome size ranging between ∼927 Mbp and ∼1.2 Gbp (data not shown), when cross-referenced with the known *Drosophila melanogaster* genome size of ∼220 Mbp (Bosco et al., 2007; Hjelmen et al., 2019), consistent with previous estimates of the *P. dumerilii* genome size (Jha et al., 1995). We thus reasoned that the large genome assembly size of 3.41 Gbp likely reflected a high number of recurrent contigs within the assembly.

To test this, we searched for universal single-copy gene orthologs in the assembly using BUSCO (Simão et al., 2015), and found that a large portion of the assembly contained recurrent and/or duplicate sequences i.e. BUSCO analysis with a metazoa database of 954 genes revealed 96.4% complete genes of which only 20.4% were single-copy and 76.0% duplicate, with 2.3% fragmented. We then identified and separated out redundant contigs and probable haplotypes in the assembly as per Guan and colleagues (Guan et al., 2020) and Roach and colleagues (Roach et al., 2020) (Methods and Materials). Iterative ‘purging’ (i.e. removal of likely redundant sequences/contigs within the assembly) via the purge_dups and purge_haplotigs algorithms, significantly decreased duplicate contigs (and haplotypes) in the assembly, while retaining similar BUSCO gene completion scores with 95.4% complete (89.0% single-copy, 6.4% duplicate) and 3.1% fragmented gene models. To further eliminate likely redundant contigs in this assembly, we filtered out sequences less than 200 Kbp (Mutemi, 2023). This decreased BUSCO genome completion scores by 1.6% (93.8%), and the number of duplicate sequences by 3.5% (90.9% single-copy, 2.9% duplicate) and 3.1% fragmented. Notably, the initial assembly size decreased from ∼3.41 to ∼1.47 Gbp, with the number of contigs also decreasing from the original 9,431 to 964; with an increase in N50 from 640 Kbp to ∼4.3 Mbp.

We then scaffolded the assembly using both long-reads and HiC data. Firstly, we used LINKS (Warren et al., 2015), an algorithm that relies on iterative k-mer pair matching over varying sequence lengths present in the raw long-read sequencing data and the assembly, to then build links between contigs. This approach scaffolded our assembly from 964 contigs to 647 scaffolds, increasing the N50 from 4.3 to 7.95 Mbp. We then polished the genome via POLCA (Zimin & Salzberg, 2020), using dense (∼100X coverage assuming a ∼1 Gbp genome size), high-quality accurate Illumina ∼150bp paired-end sequenced reads (Methods and Materials), returning a consensus assembly quality value of 99.74%. Next, we made use of Hi-C mapping and scaffolding via the SALSA2 pipeline (Ghurye et al., 2019), resulting in 330 scaffolds with an N50 of 54.8 Mbp (Fig. 2). *P. dumerilii* is thought to have 14 pairs of chromosomes (2n=28), as measured by C-banding and silver staining methods (Jha et al., 1995). 50% of the total assembly is represented by 8 scaffolds (Fig. 2; Supplementary Fig. 4). The genome assembly is available on NCBI under accession GCA_026936325.1.

**Figure 2.**
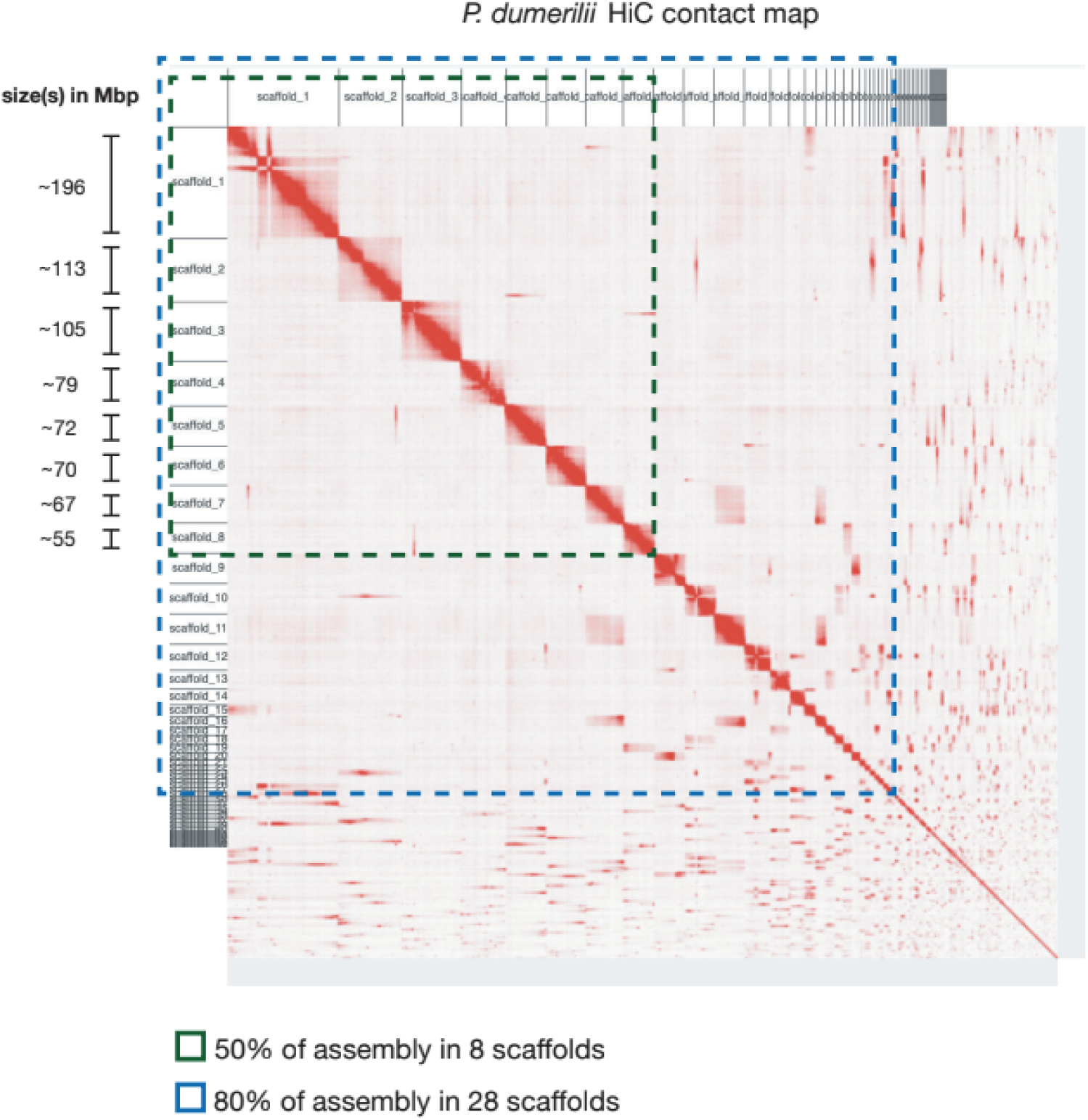
A chromosomal scale *P. dumerilii* genome assembly. A Hi-C contact map of all 330 *P. dumerilii* scaffolds. Highlighted in green are the 8 scaffolds that make up 50% of the assembly and in blue are the 28 scaffolds amounting to 80% of the assembly.

### *P. dumerilii* genome annotation

#### Repeat elements

The *P. dumerilii* genome is among the largest assembled and annotated annelid genomes to date (Bhambri et al., 2018; Martín-Durán et al., 2021; Martín-Zamora et al., 2023; Simakov et al., 2013; Zakas et al., 2022; Zwarycz et al., 2015) (Fig. 1C). Repeat elements (REs) comprise significant portions of animal genomes, and are thought to be major drivers of genome size evolution (López-Flores & Garrido-Ramos, 2012). Recent studies in the marine annelids *Dimorphilus gyrociliatus* (Martín-Durán et al., 2021) and *Streblospio benedictii* (Zakas et al., 2022) provided evidence supporting the hypothesis that the proportion of REs scale with genome size; *D. gyrociliatus* reported to have a genome size of ∼71 Mbp, with REs accounting for ∼8% of its genome content (Martín-Durán et al., 2021), in contrast to the ∼700 Mbp genome of *S. benedictii* and the ∼519 Mbp *Owenia fusiformis* genome revealing ∼43% RE content (Martín-Zamora et al., 2023; Zakas et al., 2022).

We identified and modeled REs using RepeatModeler (Flynn et al., 2020), and masked the genome using RepeatMasker (Tarailo-Graovac & Chen, 2009) (Materials and Methods). We then estimated ∼51% of the *P. dumerilii* genome to be composed of REs (Fig. 3A; Table 1), listing as the annelid with the largest proportion of genomic RE-content sequenced and annotated thus far (Fig. 3B). Of the known REs, retroelements comprise the largest proportion, occupying 14.62% of the *P. dumerilii* genome, with low-complexity REs only making up 0.11% of the total genome (Fig. 3A-B). This was consistent also across the five annelid genomes studied. In *H. robusta* and *D. gyrociliatus* however, simple repeats make up an equally high fraction of REs (*H. robusta*; ∼10.74% retroelements and ∼9.67% simple repeats, and *D. gyrociliatus*; ∼2.65% retroelements and 2.5% simple repeats; Fig. 3B). Across all annelids explored however, the ‘unclassified’ REs made up the bulk of repetitive sequences found in the genome. Despite comprising such a large fraction of the genome (particularly in the larger *O. fusiformis*, *S. benedictii* and *P. dumerilii* assemblies) very little is known about these Unclassified REs. We attempted to group ‘unclassified/unknown’ REs found in annelids and found very little similarity across all five species, with these elements clustering in a species-specific manner (Supplementary Fig. 5). This finding suggests that the bulk of REs found within annelid genomes are likely lineage- and/or species-specific.

**Figure 3.**
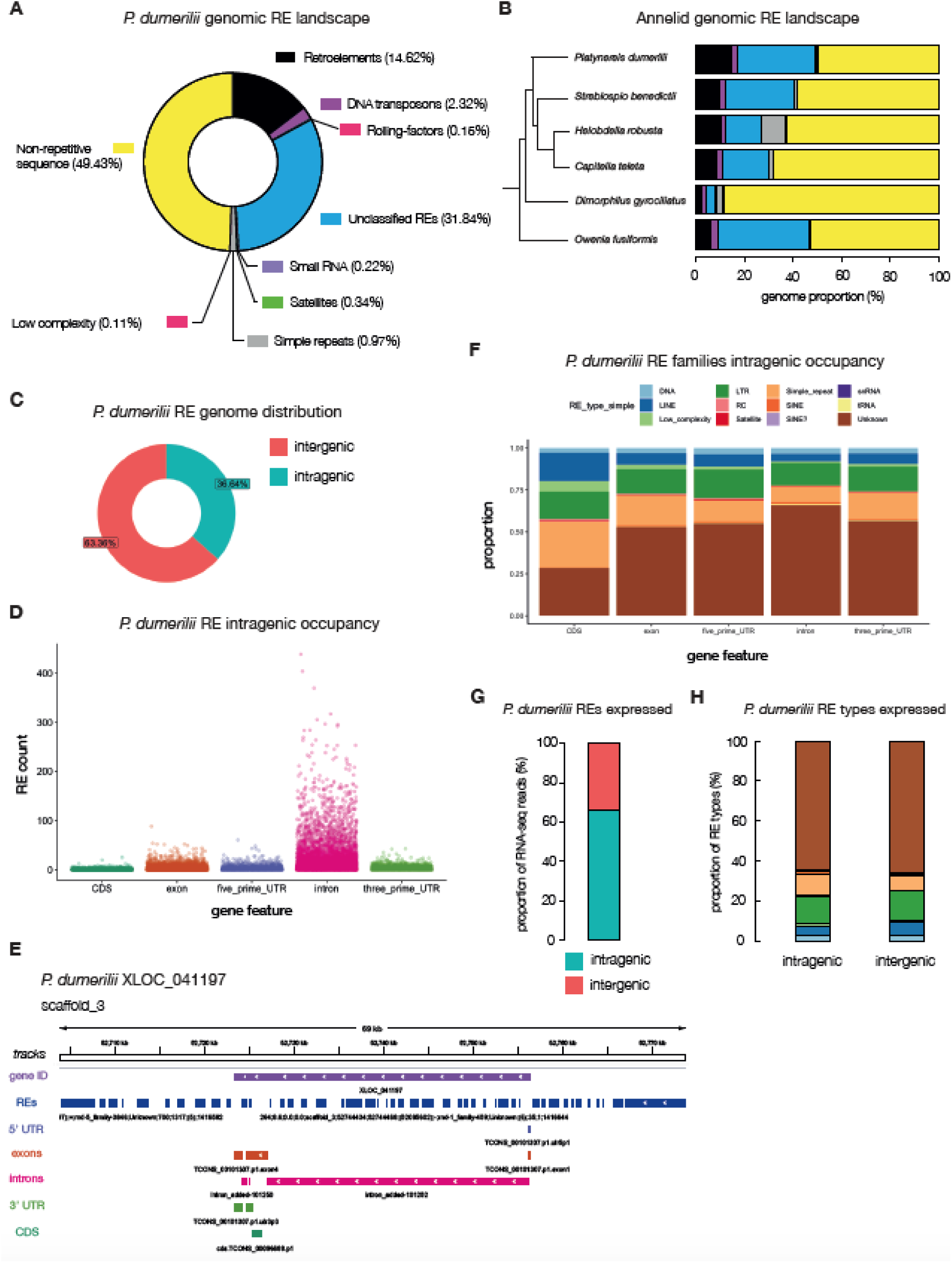
The repeat-element landscape in *P. dumerilii*. **A**, a doughnut plot illustrating the percentages of repeat and non-repeat elements found in the *P. dumerilii* genome. Percentages are of the total assembly (i.e. 49.43% of the entire genome is annotated as non-repetitive; yellow). **B**, annelids – whose relationships are shown in a phylogenetic tree – genomic repeat-element landscape. **C**, the distribution of intra – vs – inter-genic *P. dumerilii* repeat elements. **D**, counts of repeat elements represented as scatterplots within annotated intragenic regions of the *P. dumerilii* genome. **E**, an example gene locus (XLOC_041197) and its flanking regions on scaffold_3 highlighting repeat-element tracks (dark-blue) with the 5’ and 3’ UTRs (light-purple and green tracks respectively), exons (dark-orange track), introns (pink tracks) and the CDS (green tracks). **F**, proportion of repeat-element families and their occupancy at different intragenic regions. **G**, proportion of repeat-element specific RNA-seq reads mapping to intra – vs – inter-genic sites in *P. dumerilii*. **H**, proportion of RNA-seq reads mapping to intra – vs – inter-genic sites within specific RE types, colored according to the same legend in panel **F**.

**Table 1.**
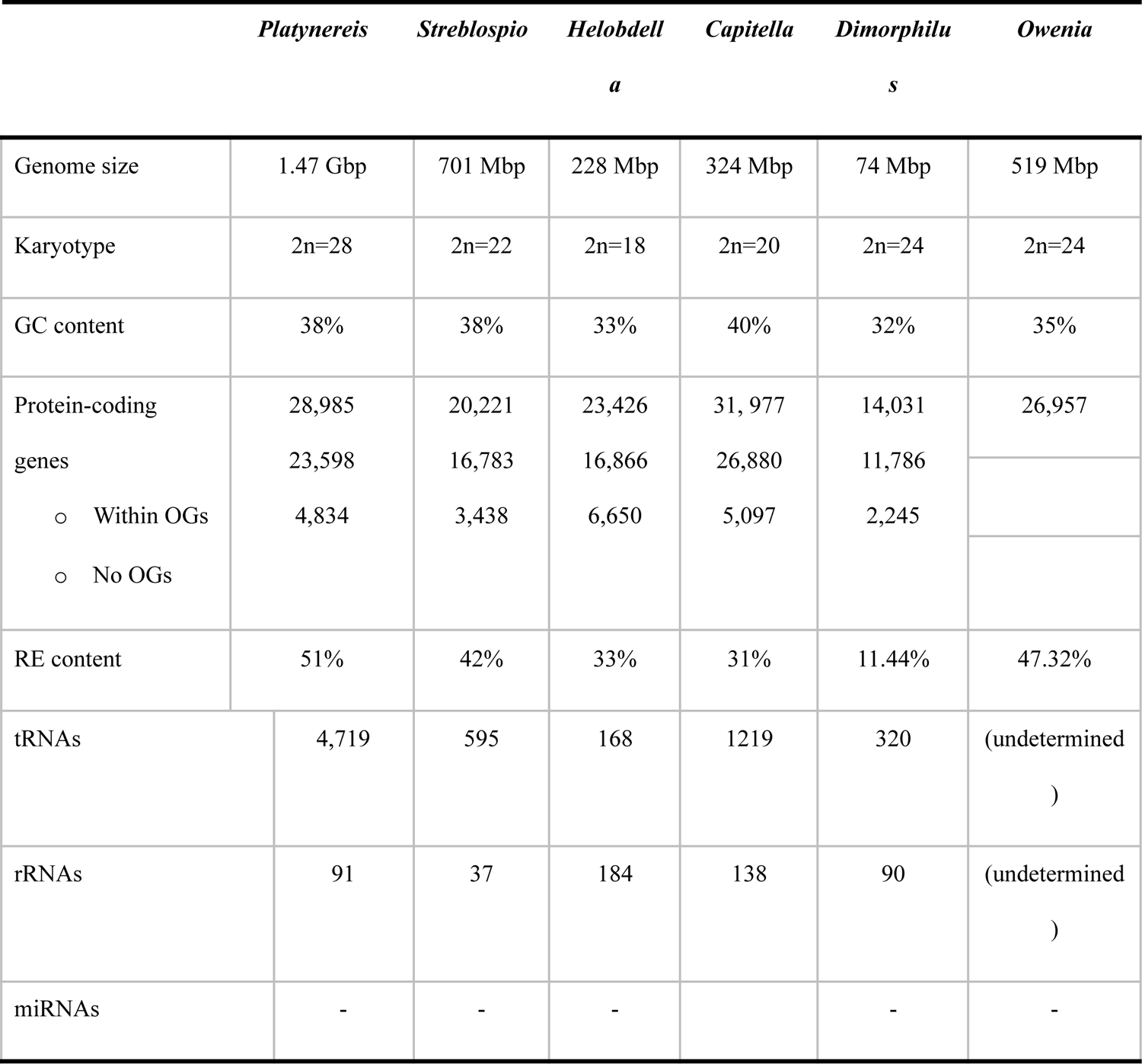
Comparative genome assembly attributes in annelids.

*P. dumerilii* RE sizes varied significantly according to the type of REs (Table 2; Supplementary Fig. 6A) (Kruskal-Wallis test, df = 11, Benjamini and Hochberg adjusted *p-value < 2.2e-16*). To explore whether the number of REs as well as their lengths partially explain genome size expansion or compaction, we also compared RE lengths across the five sequenced and annotated annelid genomes with *P. dumerilii* (Supplementary Fig. 6). We found that RE lengths were not always increased in larger genomes (Supplementary Fig. 6). For instance, *S. benedictii* unclassified REs (n = 1,155,954; median ± MAD = 126 ± 90.44 bp) were shorter than *O. fusiformis* (n = 772,141; median ± MAD = 147 ± 120.09 bp), despite the *S. benedictii* genome being larger (Supplementary Fig. 6B) (pairwise Wilcoxon rank sum test with continuity correction, *p-value < 2e-16*). In *D. gyrociliatus,* DNA Retroelements (n = 4,001; median ± MAD = 201 ± 200) were longer than all other annelids except *C. teleta* (n = 23,545; median ± MAD = 207 ± 195.70; pairwise Wilcoxon rank sum test with continuity correction, *p-value = 0.024*) despite having the smallest genome (Supplementary Fig. 6B). While this may be impacted by the repeat identification and assembly quality, we conclude that it is the number of REs, rather than their lengths that best explain genome expansion in the annelid genomes analyzed thus far.

**Table 2.** Repeat element sizes in P. dumerilii.

| **RE type** | **No. elements** | **mean ± SD (bp)** | **median (MAD) (bp)** | **min - max (bp)** |
| --- | --- | --- | --- | --- |
| DNA | 96,971 | 362.21 ± 517.76 | 200 (173.46) | 9 - 11,327 |
| LINE | 150,127 | 497.39 ± 644.24 | 258 (272.80) | 9 - 13,478 |
| Low complexity | 22,053 | 70.45 ± 118.06 | 43 (16.31) | 9 - 4,301 |
| LTR | 394,173 | 374.79 ± 729.65 | 192 (173.46) | 9 - 21,967 |
| Rolling-circles | 6,196 | 359.00 ± 900.17 | 162 (80.60) | 9 - 9,295 |
| Satellite | 8,663 | 644.00 ± 1,295.79 | 226 (263.90) | 9 - 75,744 |
| Simple repeats | 225,376 | 62.48 ± 76.84 | 42 (23.72) | 4 - 7,786 |
| SINE | 24,734 | 153.45 ± 83.79 | 154 (74.13) | 9 - 1,100 |
| SINE? | 71 | 102.30 ± 7.54 | 104 (0) | 44 - 106 |
| snRNA | 73 | 158.95 ± 76.40 | 212 (23.72) | 37 - 243 |
| *tRNA | 16,985 | 196.19 ± 242.55 | 150 (115.64) | 9 - 8,727 |
| Unclassified | 1,797,443 | 272.49 ± 348.39 | 170 (140.85) | 9 - 106,830 |

We also explored the distribution of REs across the genome and found that approximately 36% of the *P. dumerilii* REs have some overlap with protein-coding genes loci, mainly in the intronic regions, with the majority of other REs residing in intergenic regions (Fig. 3C). A similar analysis performed for the other annelid genomes showed consistently that REs predominantly occupy intergenic portions of the genome, with the major exception being *D. gyrociliatus* (Supplementary Fig. 5), showing ∼91% of REs found within genic regions; likely due to increased compaction of its genome (Martín-Durán et al., 2021).

In *P. dumerilii* protein-coding genes (see below), we found that most genes had some overlap with REs, with only around 9% - 2,608 out 28,985 genes - having no overlap with annotated REs. The majority of the annotated *P. dumerilii* REs within gene loci reside in intronic regions (Fig. 3D), which partially explains the increased size of protein coding genes in *P. dumerilii* (see below). Overall, we observed that ‘Unknown/Unclassified’ families comprised the majority of REs across all intragenic regions, followed by long-terminal repeats (LTRs) (Fig. 3E).

We also assessed the expression of REs, via the mapping RNA-seq reads to the annotated *P. dumerilii* genome (Supplementary Fig. 7). We found that an estimated ∼68% of all annotated REs were expressed (above a threshold cut-off of 10 unique mapping reads) (Fig. 3G). Of these, 66% of reads overlapped within annotated protein-coding gene loci, suggesting that the majority of RE expression likely originates from intragenic sites (Fig. 3H). We have yet to test if this is true RE specific expression, or if REs are “by-riders” of the host gene expression, as many REs sit within intronic regions of the gene loci. All RE types were expressed in both intragenic and intergenic regions, suggesting no overall differences in RE family expression (Supplementary Fig. 7B).

### Gene content and evolution

#### Protein-coding genes

To model protein-coding genes in *P. dumerilii*, we aggregated good-quality publicly available Illumina paired-end short RNA-seq reads (Conzelmann et al., 2013), spanning multiple developmental stages in *P. dumerilii* - allowing us to access all major stages of the *P. dumerilii* life-cycle - and mapped the reads to the genome using STAR (Dobin et al., 2012). We also mapped PacBio (Materials and Methods) and Nanopore long-read RNA sequencing datasets (Paré et al., 2023) (spanning early developmental stages and regenerating samples; see Methods and Materials), using Minimap2 (Li, 2018), to capture isoform diversity and full-length genes. A genome-guided *de novo* transcriptome, combining all three transcriptomic datasets, was then reconstructed using StringTie (Kovaka et al., 2019; Pertea et al., 2015). To survey the protein-coding landscape of the *P. dumerilii* genome, we searched the transcriptome for open reading frames (ORFs) using TransDecoder (Haas, 2018). In total, we modeled 28,985 protein-coding genes in *P. dumerilii*. Many genes contained at least three transcripts (∼3.2 per locus), underscoring that alternative splicing likely also plays a key role in proteome diversity in *P. dumerilii*, as in other species.

#### Gene content

Previous analyses have proposed that although there is a correlation of protein-gene content to genome size, this correlation is relatively weak compared to the non-coding genome and REs (Hou & Lin, 2009). We explored this notion further for annelids, taking into account previous genome analyses (Martín-Durán et al., 2021; Martín-Zamora et al., 2023; Simakov et al., 2013; Zakas et al., 2022) (Fig. 4 and Table 1). The total gene count in *P. dumerilii* was higher than in most annelids analyzed, with *C. teleta* being the only exception with ∼400 more genes than *P. dumerilii*, despite measuring a smaller genome size (Table 1). Despite the small sample size (N = 6 species), a formal test for correlation between protein-coding gene counts and genome size within annelids revealed a weak association (*Spearman’s rank correlation rho*, ⍴: *r_s_*(4) = 0.43, *p-value* = 0.419), consistent with previous reports in many other species (Hou & Lin, 2009). However, this may be impacted by spurious gene predictions in larger genomes.

**Figure 4.**
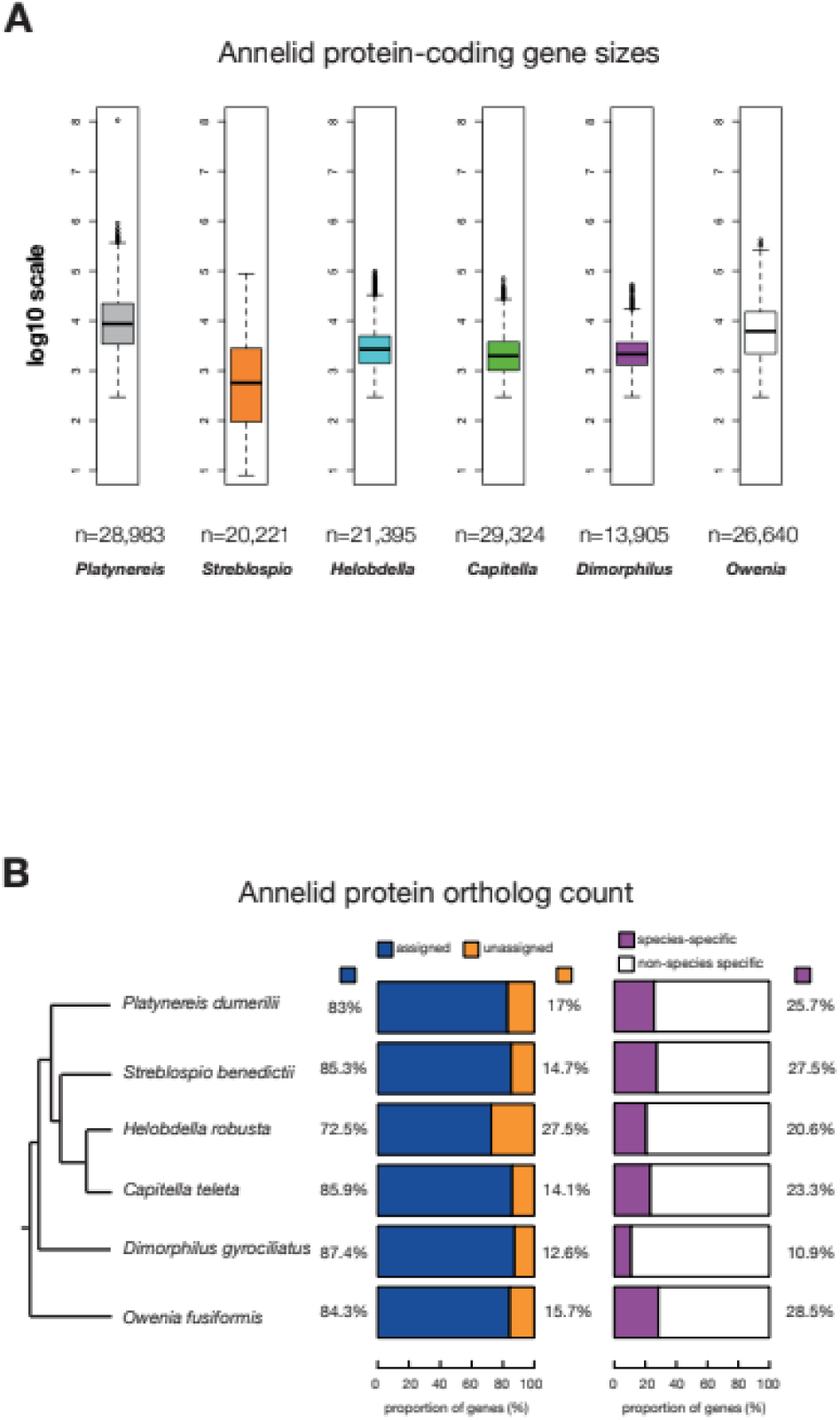
The protein coding repertoire in annelids. A, Annelid protein coding gene sizes plotted in log_10_ scale. The n values represent the total number of protein-coding genes that were measured for gene size, spanning the actual gene locus i.e. exons, introns and UTRs. The longest isoforms per gene were selected for the analysis. **B,** Proportion of annelid protein-coding genes in orthogroups.

We then explored whether gene sizes may have played a significant role in genome expansion in *P. dumerilii*. We extracted overall gene size (including UTRs, exons and introns) and compared *P. dumerilii* to *S. benedictii*, *H. robusta*, *C. teleta*, *D. gyrociliatus* and *O. fusiformis* (Fig. 4, Table 3). *P. dumerilii* genes (n = 28,983) tended to be greater in lengths than the other annelid genes (median ± MAD = 8,693 ± 9,939 bp) (Fig. 4A and Table 3). This measures to a significantly larger gene size than that of the smallest annelid genome *D. gyrociliatus* (n_genes_ = 13,905; median ± MAD = 2,155 ± 1,524 bp), and also much larger than the ∼700 Mbp *S. benedictii* genome (n_genes_ = 20,221; median ± MAD = 1,043 ± 1,290 bp). As this analysis includes all gene features including exons, introns, 5’UTR and 3’UTR and their respective sizes, we wanted to further test if there is a specific gene feature that would explain the gene size expansion in *P. dumerilii* relative to other annelids (Supplementary Fig. 8; Table 3). Across all annelid species investigated except *P. dumerilii* we consistently observed that per gene exon and intron counts were highly correlated with gene size (Pearson’s product-moment correlation *r* > 0.6; Supplementary Fig. 9). *P. dumerilii* exon counts were weakly correlated with transcript size (Pearson’s product-moment correlation *r* = 0.08, *p-value* < 2.2e^-16^; Supplementary Fig. 9A). The average exon size per gene was however poorly correlated with gene size in *D. gyrociliatus* (Pearson’s product-moment correlation *r* = 0.05, *p-value* < 2.2e^-16^) and negatively correlated with gene size in all other annelids (Pearson’s product-moment correlation *r* < −0.01; Supplementary Fig. 9A-B). Unlike exon size however, intronic size was better correlated with gene size in all annelids analyzed (Pearson’s product-moment correlation *r* > 0.27; Supplementary Fig. 9A-B, Table 3), consistent with previous hypotheses supporting the role of intron size in gene size expansion. We also calculated the median values of all exon, intron 5’- and 3’- UTR sizes, and counts across all five annelid species included in this study (Table 2). We found that of all the features that show any correlation with gene size, the 3’UTR and the intron sizes were larger in larger genomes. Taken together, we argue that 3’UTR and intron sizes differences best explain the variation in gene/transcript length sizes in annelids, relative to other gene features such as counts (Table 3, Supplementary Fig. 8 and 9). The increase in intron size in *P. dumerilii* genes is likely a result of REs occupying these regions (Fig. 3D-E).

**Table 3.**
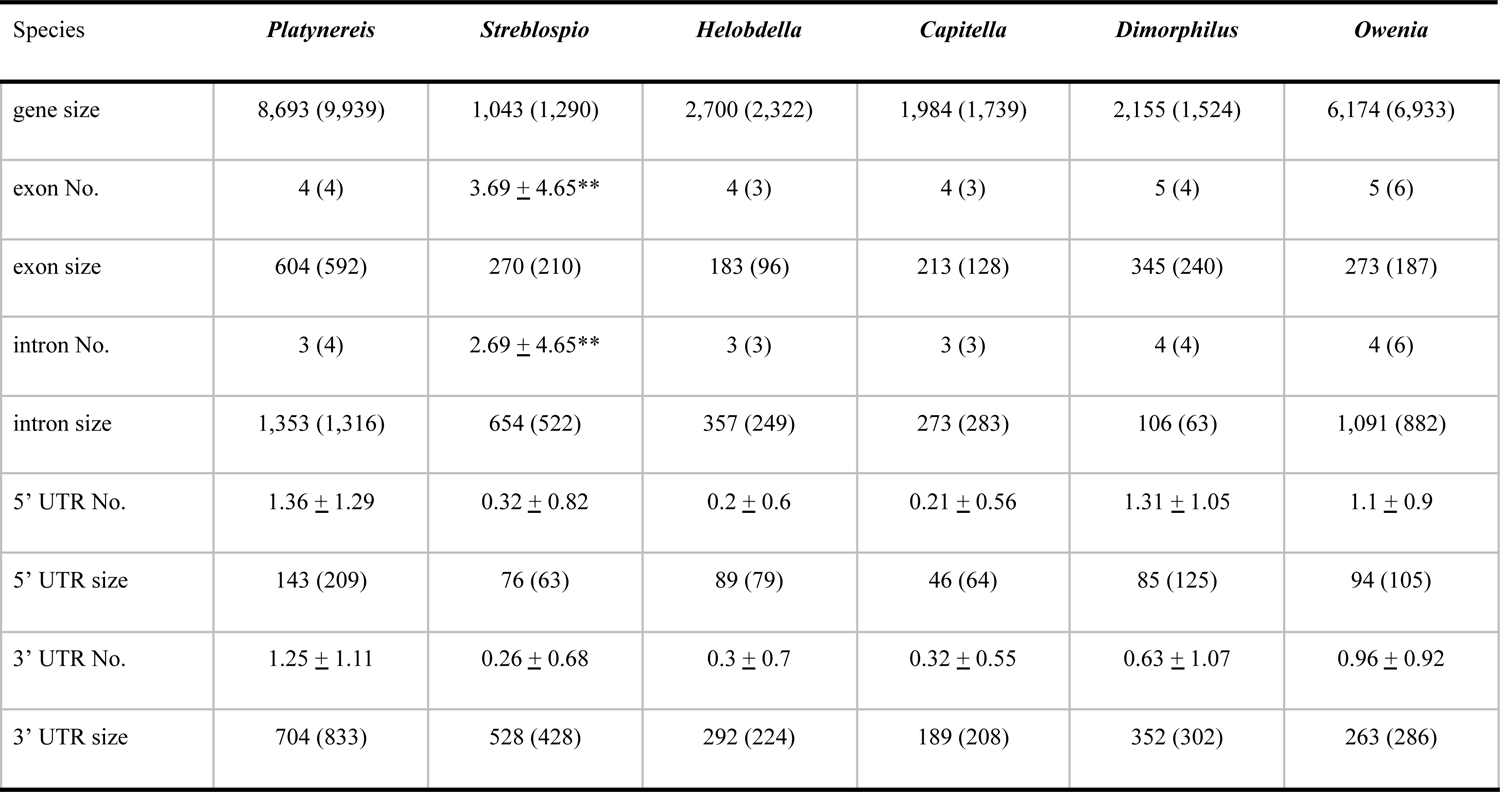
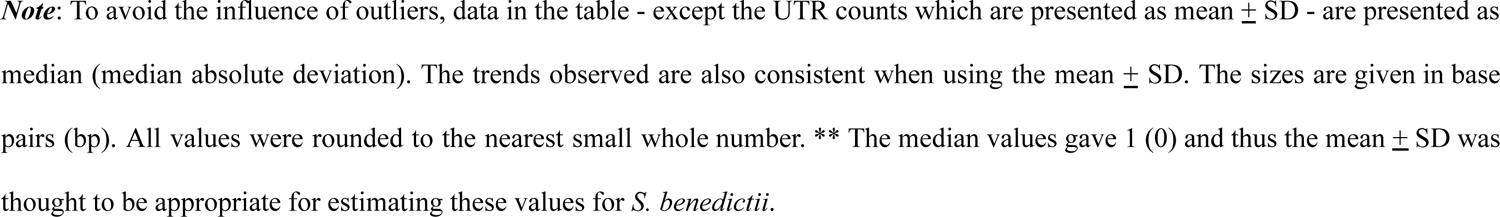
Protein-coding gene feature size and counts in annelids.

| Species | *Platynereis* | *Streblospio* | *Helobdella* | *Capitella* | *Dimorphilus* | *Owenia* |
| --- | --- | --- | --- | --- | --- | --- |
| gene size | 8,693 (9,939) | 1,043 (1,290) | 2,700 (2,322) | 1,984 (1,739) | 2,155 (1,524) | 6,174 (6,933) |
| exon No. | 4 (4) | 3.69 ± 4.65** | 4 (3) | 4 (3) | 5 (4) | 5 (6) |
| exon size | 604 (592) | 270 (210) | 183 (96) | 213 (128) | 345 (240) | 273 (187) |
| intron No. | 3 (4) | 2.69 ± 4.65** | 3 (3) | 3 (3) | 4 (4) | 4 (6) |
| intron size | 1,353 (1,316) | 654 (522) | 357 (249) | 273 (283) | 106 (63) | 1,091 (882) |
| 5' UTR No. | 1.36 ± 1.29 | 0.32 ± 0.82 | 0.2 ± 0.6 | 0.21 ± 0.56 | 1.31 ± 1.05 | 1.1 ± 0.9 |
| 5' UTR size | 143 (209) | 76 (63) | 89 (79) | 46 (64) | 85 (125) | 94 (105) |
| 3' UTR No. | 1.25 ± 1.11 | 0.26 ± 0.68 | 0.3 ± 0.7 | 0.32 ± 0.55 | 0.63 ± 1.07 | 0.96 ± 0.92 |
| 3' UTR size | 704 (833) | 528 (428) | 292 (224) | 189 (208) | 352 (302) | 263 (286) |

**Note:** To avoid the influence of outliers, data in the table - except the UTR counts which are presented as mean ± SD - are presented as median (median absolute deviation). The trends observed are also consistent when using the mean ± SD. The sizes are given in base pairs (bp). All values were rounded to the nearest small whole number. \*\* The median values gave 1 (0) and thus the mean ± SD was thought to be appropriate for estimating these values for *S. benedictii*.

We then analyzed orthology of protein-coding genes in annelids. We first extracted the longest isoform for each protein-coding gene in *P. dumerilii* and downloaded publicly available proteome datasets for the other annelid genomes (Methods and Materials). Using OrthoFinde*r* (Emms & Kelly, 2015), we found 116,625 genes (∼83%) of the total proteome (i.e. all six annelid proteomes used in the dataset) could be assigned to 17,752 orthogroups. The percentage of the protein-coding genes in orthogroups varied from ∼72% (*H. robusta* - 15,505 genes) to ∼87% (*D. gyrociliatus* - 12,155 genes), with around ∼83-84% in *P. dumerilii* - 24,053 and *O. fusiformis* - 22,468 genes, with ∼ 85-86% in *S. benedictii* - 17,241 and *C. teleta* - 25,203 genes (Fig. 4B; Table 1). We interpret that those genes without orthogroups could represent novel single-copy genes (i.e. do not belong to a family) for that particular species, or its lineage. Alternatively, those genes may be modeling artifacts. Increasing the number of sequenced and annotated annelid genomes will help distinguish between these two possibilities, allowing the distinction between species-specific (or novel) single-copy and lineage-specific genes. Gene ontology (GO) term analysis showed minimal differences in GO-term enrichment, when comparing genes assigned to unassigned orthogroups (Supplementary Fig. 10); suggesting that these two different gene sets are likely similar in terms of cellular, molecular function and biological processes.

Further analysis highlighted 25.7% of *P. dumerilii* genes with an orthology assignment as *P. dumerilii*-specific (i.e. these genes form orthogroups with other *P. dumerilii* genes but no other annelid genes), with *D. gyrociliatus* possessing 10.9% species-specific orthogroups (Fig. 4B). We note that the total gene count, proportion of genes in orthogroups or species-specific orthogroups was not correlated with genome size (Fig. 4B; Pearson’s product-moment correlation, *p* > 0.1). This result suggests that gene duplication is not a major factor in genome size expansion in the annelid species analyzed thus far.

#### A diverse and rich repertoire of RNA genes in P. dumerilii

We modeled the non-coding transcriptome of *P. dumerilii* using both RNA-seq reads (putative lncRNAs and miRNAs) and *de novo* genome searches for tRNAs and rRNAs.

#### Transfer RNAs

*P. dumerilii* tRNAs were modeled *de novo* from the genome sequence alone using tRNAscan-SE (Chan et al., 2019; Lowe & Eddy, 1997) (Materials and Methods). We identified approximately 4,719 high-confidence tRNA genes (excluding the mitochondrial genome) in *P. dumerilii*, ranking as one of the few animal genomes annotated thus far to have such a high number of tRNA copy genes (Table 1). Consistent with previous findings, we do not find an obvious association of tRNA gene number and other genome and/or organismal traits (Bermudez-Santana et al., 2010). We did note however that high-confidence tRNA counts were much higher in *P. dumerilii* than in the other annelid genomes with *C. teleta* having 1,219 genes and *H. robusta* possessing the fewest tRNA genes (168) (Table 1). We did not see any association of tRNA gene count and genome size as *D. gyrociliatus* had an estimated 320 tRNA genes (nearly double that of *H. robusta*), despite having a genome size 6 times smaller than *H. robusta* (Table 1).

#### Long non-coding RNAs

We made use of the available RNA-seq datasets described before to model putative lncRNAs. When generating the transcriptome, we limited further analyses to transcripts with a minimum length of 200 nucleotides (Materials and Methods). Of the total 67,020 loci, 28,985 were protein coding with a remaining set of 38,035 non-coding loci. This non-coding gene set comprised ∼287 Mbp (∼19.5% of the total *P. dumerilii* genome). Incorporating these sequences with databases that contain lncRNAs, although promising, remains a challenge as many of these annotated lncRNA sequences are from model organisms that are phylogenetically distant from *Platynereis* and further experimental validation is necessary (Cao et al., 2018; Zhao et al., 2021). Another recent study in *P. dumerilii* has also identified putative lncRNAs with specific expression in germ cells (Ribeiro et al., 2024).

#### microRNAs

miRNAs are small non-coding RNAs that negatively and post-transcriptionally regulate the expression of target mRNAs (Bartel, 2018). Previous studies in *P. dumerilii* identified 34 miRNA genes common to protostomes and deuterostomes (Christodoulou et al., 2010). A more rigorous survey of *P. dumerilii* miRNAs, however, is lacking. To this end, we sequenced small RNAs from multiple early developmental stages for de novo prediction of miRNA genes using miRDeep2 (Friedländer et al., 2012), identifying 587 miRNA unique gene loci (Supplementary File 1[1]). However, because miRDeep2 identifies numerous candidate miRNAs without, for example, a sequenced star read, we next used MirMine (Wheeler et al., 2009) coupled with MirMachine (Umu et al. 2023) to identify a suite of high-confidence miRNA loci using the processing rules established by Ambros et al. (AMBROS et al., 2003) (Fromm, 2024)). We identify 113 miRNA genes belonging to 61 phylogenetically conserved miRNA families and an additional eight miRNA loci specific to *P. dumerilii* (Supplementary File 2 and 3) (Supplementary Fig. 11) (all data deposited in MirGeneDB 3.0, (Clarke et al., 2024)). Like the polychaete annelid *Capitella teleta*, *P. dumerilii* retains a complete repertoire of ancestral miRNA families and has only lost four miRNA paralogues: Mir-10-P6, a Mir-87 paralogue, Mir-193-P1, and Mir-216-P2c.

Because of the unique mode of miRNA evolution (Fromm, 2024; Tarver et al., 2013, 2018), loci are now known that are clade-specific at several hierarchical levels within the annelid lineage, ranging from Lophoptrochozoa (e.g., MIR-1986), the phylum itself (Mir-1995, MIR-2692), and to specific subgroups within Annelida including Pleistoannelida (e.g., MIR-1987). Further, these data support the earlier observation based on miRNAs (Sperling et al., 2009) that sipunculids are nested within the annelids, consistent with numerous phylogenomic analyses (Weigert & Bleidorn, 2016) (Fig. 6A). Several of these novel miRNA genes are genomically linked such that any two genes lie within 50 kb of one another, resulting in the production of polycistronic transcripts, i.e., a single long non-coding mRNA that contains two or more miRNA sequences (Baskerville and Bartel 2005) and thus several of these novel miRNAs are co-expressed with other, also novel miRNA transcripts (Fig. 6A).

Clustering is typical for miRNA loci in most animal species, including *P. dumerilii* (Fig. 6B). In fact, similar to humans (MirGeneDB), just over 50% of the annotated miRNAs fall within clusters consisting of two or more loci within the 50 kb window. Most of these clusters – like the ancestral Mir-1/Mir-133 cluster – consist of two or more different family members and hence house two or more different seed sequences, which means that each unique seed sequence recognizes a unique response element in the 3’-UTR of a target mRNA sequence. Other clusters though, consist of paralogues of a single ancestral gene generated through tandem gene duplication and sometimes, but not always, possess the same seed sequence and hence target the same repertoire of mRNAs.

However, as has long been known, gene duplication can allow for the neo-functionalization of one of the two resulting genes. In the case of miRNAs, various changes to the primary sequence of the mature miRNA can occur to generate unique seed sequences. For example, a genomic hotspot for miRNA innovation is the Hox cluster (Campo-Paysaa et al., 2011; Heimberg & McGlinn, 2012), and annelids, including *P. dumerilii*, are no exception. However, unlike vertebrates and arthropods, which evolved unique miRNAs in their respective Hox complexes, annelids simply duplicated the ancestral Mir-10-P1 locus such that, primitively, the pleistoannelids house five paralogues of Mir-10-P1, one of which in a similar location to the vertebrate Mir-196 and the arthropod Iab loci (Fig. 6C, arrows, right). Interestingly though, none of these paralogues are redundant as there are cases of arm shifting. For example, in Mir-10-P4, it is the 3p arm that is used as the mature gene product (Fig. 6C, left, purple), which is opposite to the ancestral 5p arm seen in Mir-10-P1 (Fig. 6c, left, red). This of course, generates a unique seed sequence and hence a unique set of target interactions. A second way to diversify function is through seed-shifting (Wheeler et al. 2009), where the start of the mature sequence is adjusted to either the 5’ or the 3’ of the ancestral sequence. For example, the starting nucleotide (nt) of Mir-10-P5, a lophotrochozoan-specific paralogue of Mir-10-P1, is adjusted one nt 5’ of the ancestral cut (Fig. 6C, left, blue). Because the seed sequence is defined mechanistically as nucleotides 2-8 of the mature miRNA, seed shifting generates a unique seed sequence to interact with a unique set of target sites. Even when the seed sequence is the same, as is seen with Mir-10-P7 and Mir-10-P8 in relation to Mir-10-P1, changes are seen at positions 13-16, the so-called 3’ complementary region (Fig. 6C, left, orange). These nts can also base pair with target sequences, and changes to the sequence composition can have subtle yet specific effects on the regulation of target mRNAs (Broughton et al., 2016).

### Exploring natural variation in the *Platynereis* populations

*Platynereis* represents a cosmopolitan species complex (Broughton et al., 2016), with previous descriptions or citings of assumed *P. dumerilii* based on morphology likely partially attributable to *Platynereis* sister species, in particular the sympatric sibling species *P. massiliensis*. The laboratory culture sequenced here corresponds to *Platynereis dumerilii sensu stricto* as defined by Teixera et al. (Teixeira et al., 2021, 2022). We collected RNA-seq data equaling ∼20 million read pairs (for each location) from *Platynereis spp.* populations collected in six locations (Las Cruces, Chile; Algarve, Portugal; Kristineberg, Sweden; Ischia, Italy; Oban, Scotland; and Mayotte, France). These data were mapped against the current assembly and annotated genome (Fig. 5A). We found that samples from Ischia best matched our own lab culture RNA-seq data (∼47% of reads from the Ischia samples mapped to the genome as opposed to ∼56% of reads from lab culture samples), followed by samples from Faro (∼32%), Oban (∼18%), Kristineberg (∼13%) and Mayotte (∼1.2%), with Las Cruces samples showing very poor mapping statistics (∼0.6%) (Fig. 5B). This likely suggests these samples are not from the same species; highlighting the utility of our genome resource in testing how *P. dumerilii* species (including the *Platynereis spp.* complex [see below]) are related to each other (Fig. 5A).

**Figure 5.**
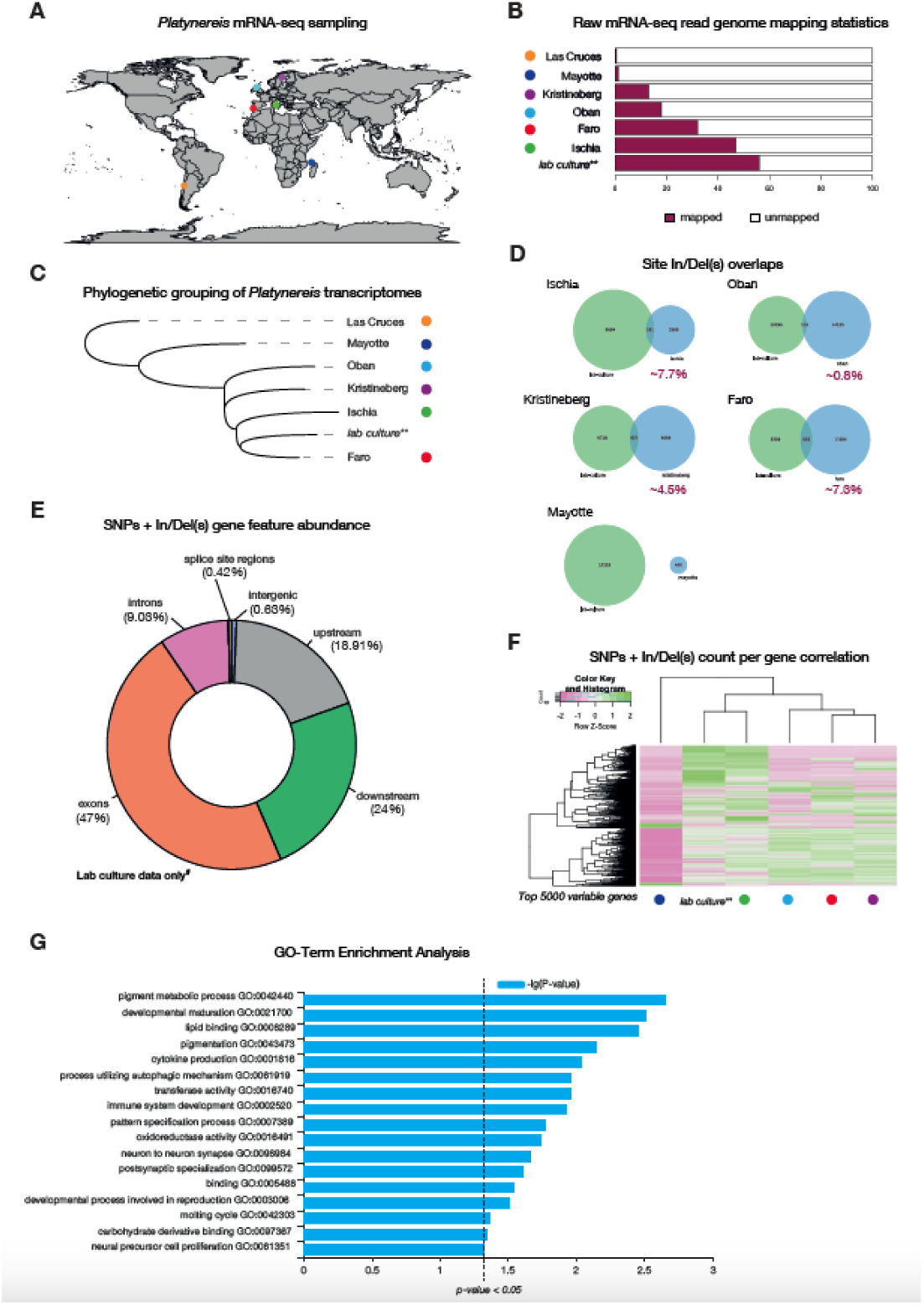
Genomic and transcriptomic variation analyses on wild sampled *P. dumerilii*. **A**, global map of sites of *P. dumerilii* mRNA sampling. **B,** histogram of raw mRNA-seq genome mapping percentages. **C**, phylogenetic grouping/sorting of wild sampled *Platynereis* transcriptomes via OrthoFinder. **D**, proportion of In/Del overlaps identified from the different *Platynereis* samples. **E**, gene feature abundance/occupancy of SNPs and In/Dels from mRNA-seq reads accessed from *P. dumerilii* lab cultures. **F**, SNP and In/Del counts from the same position on the genome correlation for the top 5,000 most variable genes (i.e. genes that showed the most variation in SNP and In/Del counts across the different sites). **G**, GO-term enrichment analysis of the top 5,000 variable genes.

**Figure 6.**
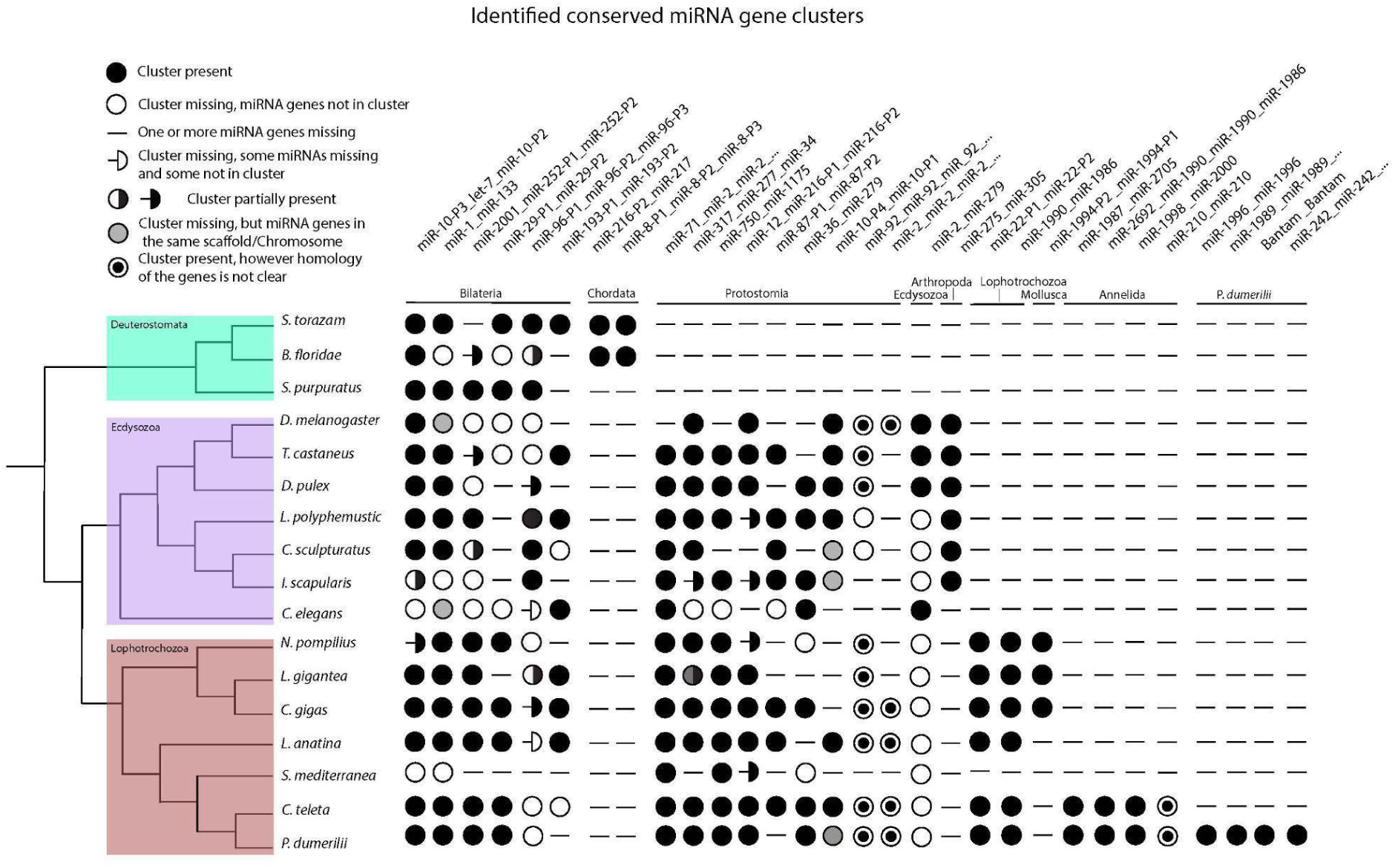
This scheme shows the distribution of phylogenetically conserved miRNA gene clusters in selected bilaterian phyla from the MirGeneDB database. Species are listed on the left, with *P. dumerilii* shown at the bottom. The tree, next to the species names, reflects state of the art of the lophotrochozoan clade phylogeny, with the branching taken from Marlétaz et al. (2019). The names of the miRNA clusters are defined by the comprised miRNA genes, separated by underscores (_), and are listed at the top of the figure. Since the order of miRNA genes in genomic clusters can vary between different species, the nomenclature follows three hierarchical criteria: 1. The gene order in the *P. dumerilii* genome; 2. The most common arrangement in the analysed species 3. Alphabetical order, when the first two criteria cannot be fulfilled. When the same miRNA gene name is repeated in the cluster, it indicates the presence of multiple copies, with uncertain homology, of the correspondent gene family. If the cluster name ends with three dots (…) more copies of the last listed gene are present. The number of copies can vary between species. miRNA clusters are grouped and ordered according to their phylogenetic conservation, with the respective clades indicated just below the clusters. Here follows the description of the symbols. Full circle: the cluster listed above is present in the corresponding specie, with all genes included in the cluster. Empty circle: miRNA genes are found in the genome but not clustered together. Hyphen (-): corresponding miRNA genes are not found in the genome. Including when only one member of a two-gene cluster is missing. For clusters composed of three or more genes, there are three additional scenarios with respective symbols: 1. Hyphen followed by an empty half-circle: some miRNA genes are absent in the genome, while others are present but not clustered; 2. Circle half full: some miRNA genes are clustered, while others are not. 3. Hyphen followed by a full half-circle: some genes are clustered, while others are not present in the genome. Grey-filled circle: genes are not clustered but are found in the same genomic scaffold or chromosome. Circle with a filled center: indicates the presence of a cluster composed of gene copies with uncertain homology.

To study phylogenetic relationships of these populations, we conducted transcriptome assemblies for each of the sites. Consistent with the raw-mapping data, Las Cruces was most distantly related to all sites, with Mayotte samples forming an outgroup to all remaining sites (Fig. 5C). Samples from Oban, Kristineberg, Ischia and Faro form a monophyletic group, including the lab cultures, suggesting that these are either all *P. dumerilii s.s.* or closely related (Teixeira et al., 2022; Valvassori et al., 2015; Wäge et al., 2017). According to the transcriptome data, we also noted that Faro samples were more closely related to lab cultures than samples from Ischia, where the culture originated in the 1960s (Fischer & Dorresteijn, 2004).

From the mapped data, we then called single nucleotide polymorphisms (SNPs) and insertions/deletions (InDels) using SnpEff & SnpSift (Cingolani et al., 2012). We found that Ischia and Faro samples shared a higher proportion of their indels (∼7%) with our lab-cultures, followed by Kristineberg (∼4.5%) and Oban (∼0.8%) samples (Fig. 5D). Mayotte samples showed no overlap of their indels with our lab cultures. As expected of RNA-seq genome mapping data from the site-specific presumed *P. dumerilii* samples, majority of the SNPs and indels were found in exons (∼47%), with ∼19% of variants called 5 Kbp upstream of the genes (not including the 5’ UTR) and 24% of variants called 5 Kbp downstream of genes (not including the 3’ UTR) (Fig. 5E). This was consistent across all sites for which we had RNA-seq data (Supplementary Figs. 12-13; for an example gene locus see Supplementary Fig. 14). We further grouped genes according to their variable SNP/In(Del) counts, identified on the different sites (Fig. 5D). We limited our analyses to a maximum of 5000 genes that showed the highest variation in SNP/In(Del) counts across sites (Fig. 5F). Hierarchical clustering of these SNP/In(Del) counts showed that for any given gene, the Ischia samples showed similar numbers of SNPs/In(Dels) to the lab cultures - acting as our reference samples - (Fig. 5F; Supplementary Fig. 14), with Mayotte samples displaying a more distinct pattern from all other samples. To explore the likely function of these genes, we analyzed their GO term enrichment using WEGO (Ye et al., 2018) (Fig. 5G, Supplementary Fig. 15). The statistically significant enriched terms encoded for terms associated with pigmentation (pigment metabolic processes [GO:0042440], pigmentation [GO:0043473]), development (developmental maturation [GO:0021700], pattern specification process [GO:0007389]) and immune-responses (cytokine production [GO:0001816], process utilizing autophagic mechanism [GO:0061919], immune system development [GO:0002520], oxidoreductase activity [GO:0016491]). Future analyses into if these differences are biologically meaningful is needed prior to any further conclusions and/or interpretations.

### Comparative genomic analyses of three *Platynereis* species: *P. dumerilii*, *P. massiliensis* and P. megalops

Currently, there are around 33 *Platynereis* species, widely distributed across the globe (Read & Fauchald, 2020), of which four species (i.e. *Platynereis bicanaliculata, Platynereis dumerilii, Platynereis massiliensis* and *Platynereis megalops*) have been well described with respect to their development, life-cycle and sexual behaviors (Fischer & Dorresteijn, 2004; Hauenschild, 1951; Just, 1914, 1922; Roe, 1975). *P. dumerilii* is sympatric - occurs in the same area - with *P. massiliensis* (Hauenschild, 1951; Valvassori et al., 2015), a sibling species that has a distinct early developmental program (Helm et al., 2014; Schneider et al., 1992). However, development then converges and individuals become morphologically indistinguishable during juvenile/adult stages. The two species can be again distinguished morphologically only after sexual maturation, based on pigmentation and mating behaviors (Fischer et al., 2010; Hauenschild, 1951; Helm et al., 2014; Lucey et al., 2015; Schneider et al., 1992; Valvassori et al., 2015). Intriguingly, *P. dumerilii* sex maturation is thought to transition directly to either male or female. On the contrary, *P. massiliensis* undergoes sequential hermaphroditism from males to females (Hauenschild, 1951; Helm et al., 2014). The mechanisms underlying these sex-determination modes are not known. *P. megalops* on the other hand, has only been described from Woods Hole (Just, 1914), or the Vineyard Sound region in Massachusetts (Verill, 1873). Curiously, *P. megalops* appears to show a very similar development to *P. dumerilii*, however the mating behavior is once more drastically distinct from that of *P. dumerilii* and *P. massiliensis* (Just, 1914, 1922). *P. dumerilii* and *P. massiliensis* display external fertilization, whereas *P. megalops* fertilization is internal. These observations highlight the ‘intra-genus’ variation in this clade, and the dynamic evolution of these phenotypes. Little is known about the genotypic/genomic variation among these diverse *Platynereis* species. The mitochondrial genomes show little difference in gene number across *Platynereis* species, with only *P. massiliensis* possessing an additional tRNA (total of 23 as opposed to 22) (Alves et al., 2020). To better explore the patterns and dynamics through which genes have been gained/lost in annelids, we carried out genome sequencing, annotation and comparative analyses of *P. massiliensis* and *P. megalops* genomes.

### Assembly statistics across the three studied *Platynereis* genomes

To address this diversity at the genomic level, we sequenced single individuals of each of the three species (Methods). We note that *P. dumerilii* assembly from a single individual results in better contiguity, however as it was generated later and may not be representative of the high polymorphism spectrum of the lab culture, it is so far provided only as-is. While genomes of *P. dumerilii* and *P. megalops* could be assembled to ‘pseudo-chromosome’ resolution, the *P. massiliensis* was only assembled to scaffold-resolution (Table 4). The assemblies were generated using the ‘hifiasm’ algorithm (Cheng et al., 2021), and further scaffolded using the Arima pipeline and SALSA2 (Materials and Methods) and are available on Genbank under accessions GCA_043381215.1, GCA_043380565.1, and GCA_043380595.1. All three assemblies showed greater than 90% BUSCO completeness values (Table 4), implying that the majority of their sequence information was captured by the sequencing data. *P. massiliensis* possessed the most fragmented assembly at 1,478 scaffolds compared to 163 and 370 for *P. dumerilii* and *P. megalops* respectively (Table 4). The *P. megalops* assembly was the largest at ∼1.88 Gbp, approximately 400 Mbp larger than both *P. dumerilii* and *P. massiliensis* at ∼1.42 and ∼1.43 Gbp respectively (Table 7). For both of the *P. dumerilii* assemblies the genome size remained stable (Table 1 and Table 4), suggesting that this assembly size may reflect the actual genome size much more fittingly than the initial ∼980 Mbp estimate (Jha et al., 1995), and the short-read DNA Illumina sequence data (see above).

**Table 4.**
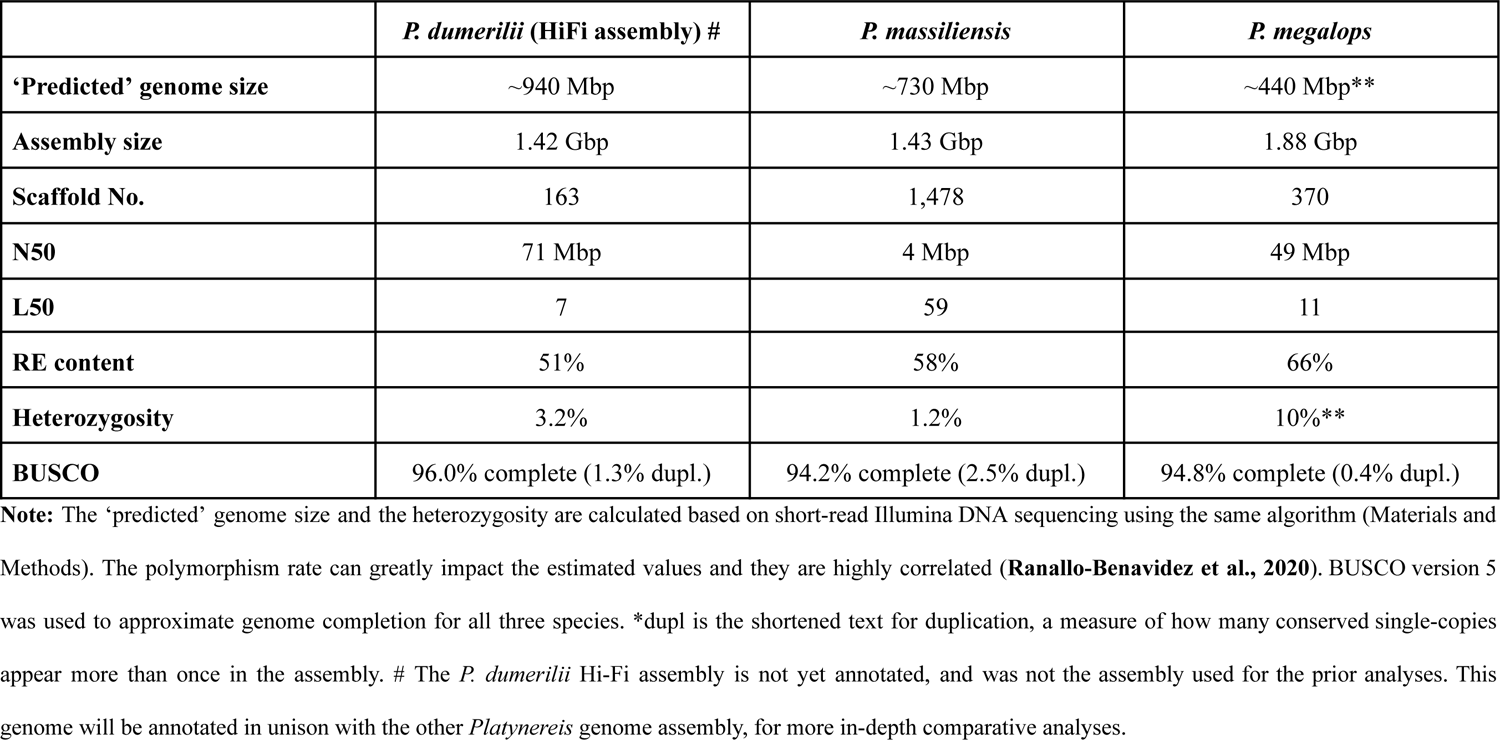
Assembly features of *Platynereis* genomes.

The previous analyses on the first version of the *P. dumerilii* genome revealed RE content to be a major player in genome size differences. To explore if this is also the case between closely related species, REs were identified, revealing that their content scaled with genome size in the three *Platynereis* species (Table 4 - 5). Of all the REs that scaled according to genome size, the Unclassified elements appeared to be the most dominant (Table 5), comprising 31% of the total genome length in *P. dumerilii* and *P. massiliensis* and 38% of *P. megalops* genome. All other REs were distributed similarly across the different species, suggesting that these elements do not change rapidly.

**Table 5.** RE distribution in *Platynereis* species.

|  | *P. dumerilii* | *P. massiliensis* | *P. megalops* |
| --- | --- | --- | --- |
| **Total genome RE content** | **50.38%** | **58.40%** | **65.96%** |
| Retroelements | 14.83% | 20.92% | 20.56% |
| *SINEs* | *0.16%* | *0.63%* | *0.28%* |
| *LINEs* | *5.00%* | *7.22%* | *7.99%* |
| *LTR elements* | *9.67%* | *13.07%* | *12.28%* |
| DNA transposons | 2.79% | 5.06% | 5.66% |
| Rolling-circles | 0.12% | 0.36% | 0.26% |
| Unclassified | 30.93% | 30.56% | 38.19% |
| Total interspersed repeats** | 48.56% | 56.54% | 64.41% |
| Small RNA | 0.36% | 0.59% | 0.11% |
| Satellites | 0.23% | 0.03% | 0.001% |
| Simple-repeats | 1.00 | 1.03% | 1.08% |
| Low-complexity | 0.11% | 0.10% | 0.09% |

**Note:** The total interspersed repeats include all REs that are not found in tandem, which include all Retroelements, DNA transposons, Rolling-circles and Unclassified repeats.

All three *Platynereis* species show overall conservation of scaffold homology, which likely extends to chromosomes. However, several breaks in collinearity within homologous regions could be observed indicative of their evolutionary distance (Fig. 7). Additionally, some translocations between scaffolds can be observed for *P. megalops*. While this suggests that overall, no major rearrangements of karyotypes had happened in these three species, further sequencing efforts are required to test chromosomal conservation and prevalence of inter-chromosomal translocations in this clade.

**Figure 7.**
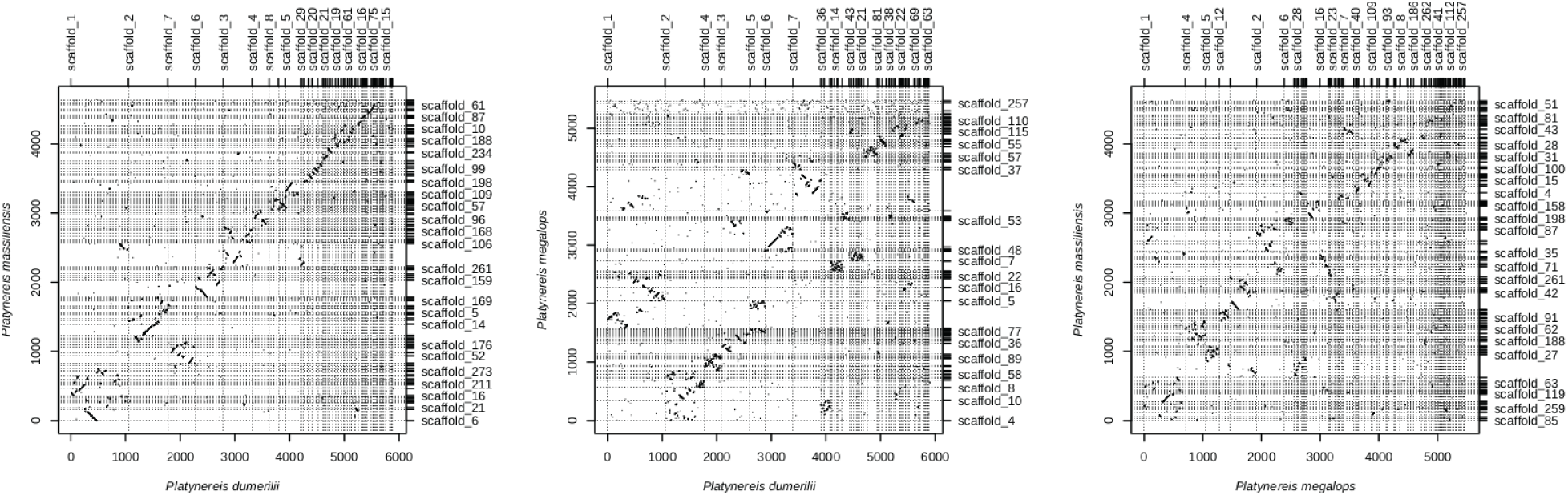
Comparison of three *Platynereis* species. Oxford dotplot comparison of the three *Platynereis* species genome assemblies. White homologies of most of the scaffolds can be identified, within scaffold inversions are common and present in all species.

### Insights into chromosomal evolution in annelids

Conserved chromosomal-level conservation (macro-synteny) can be traced over very large phylogenetic distances spanning the animal tree of life (Putnam et al., 2007, 2008; Simakov et al., 2022). Previous studies showed that Bilateria linkage groups (BLGs, (Simakov et al., 2022)) can be used to represent karyotypes of many animal genomes. Such algebraic operations involve duplications of ancestral chromosomal elements (e.g., in vertebrates (Simakov et al., 2020)) or various fusion processes (Simakov et al., 2020, 2022) with and without mixing of genes (Fig. 8A). We sought to explore the chromosomal-level organization of the *P. dumerilii* genome in the context of annelids, and more generally other Spiralia assemblies currently available and their BLG representation. In total, we analyzed 51 spiralian genomes at near-chromosomal level resolution, including 25 annelids, nine mollusks, six flatworms, four bryozoans, two nemerteans, three rotifers, a brachiopod and a phoronid (Table 6). Of particular interest to us was the identification of chromosomal fusion-with-mixing events that have been suggested to comprise strong irreversible synapomorphic characters (Simakov et al., 2020, 2022) (Fig. 8A), informative for validation of phylogenetic relationships (Schultz et al., 2023).

**Figure 8.**
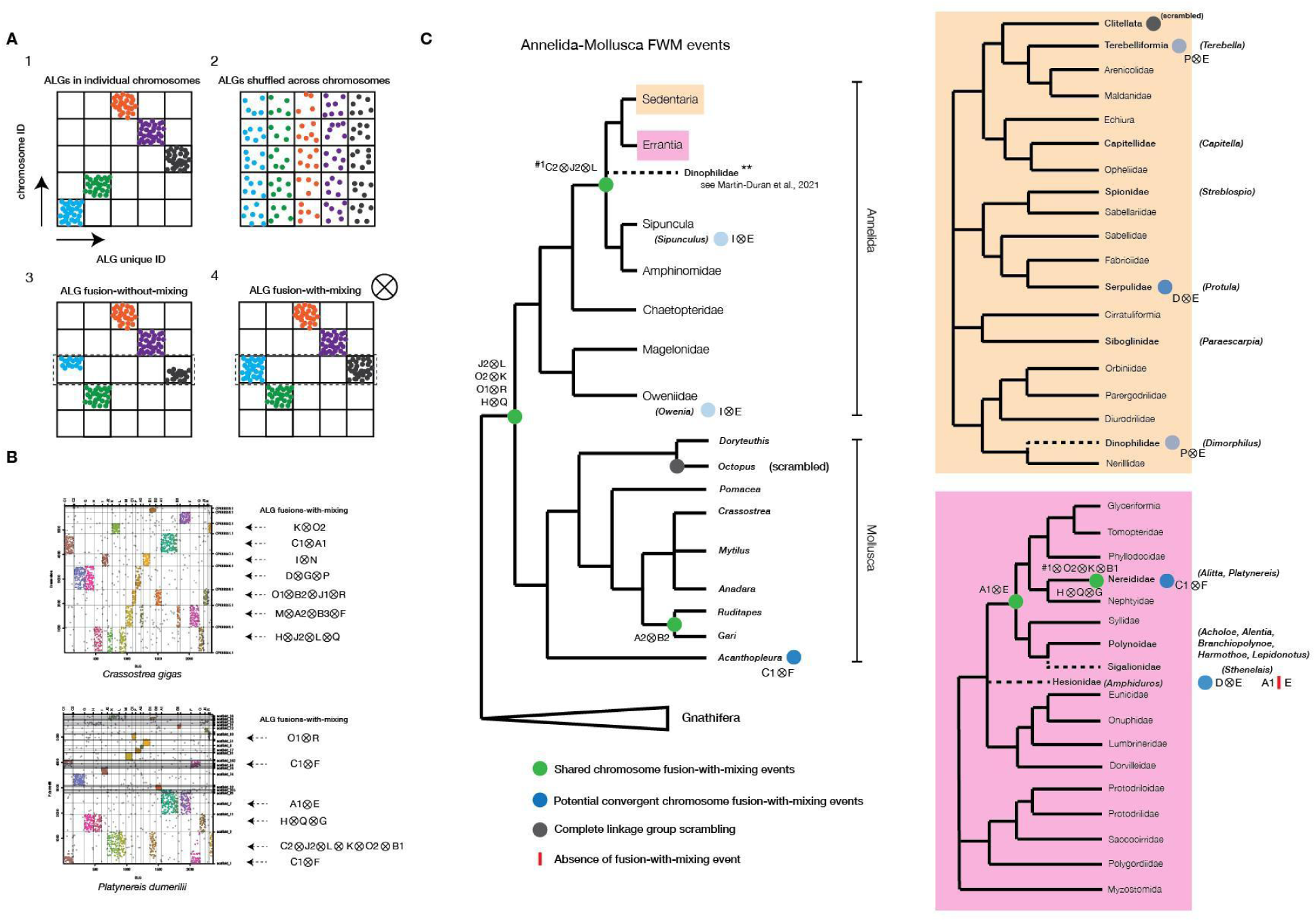
Bilaterian Ancestral Linkage Group (bALG) fusion-with-mixing events towards and within Errantia. **A,** potential linkage group evolutionary patterns described in animals. **B**, example annotation of bALGs and fusion-with-mixing (FWM) events detected in scaffolds or annotated chromosomes of *C.gigas* and *P. dumerilii* (this study). **C**, mapping of FWM events detected in this study onto the most up-to-date annelid-mollusc tree. Highlighted are the FWM events detected in annelid species belonging to specific groups within Sedentaria (orange) and Errantia (pink).

**Table 6.** Analyses of Spiralian genomes at near-chromosomal level resolution.

| Animal | Species | Karyotype | ALGs (total of 24 elements) | ALG fusions |
| --- | --- | --- | --- | --- |
| Annelids | *Acholoe squamosa* | - | all | A1⊗E, + ‘spiralian’-fusions (J2⊗L⊗C2) |
|  | *Alentia gelatinosa* | - | all | A1⊗E, I⊗N, + ‘spiralian’-fusions (H⊗Q⊗J2⊗L⊗C2, O2⊗K⊗M) |
|  | *Alitta virens* | - | all | C1⊗F, A1⊗E, + ‘spiralian’-fusions (H⊗Q⊗G and J2⊗L⊗O2⊗K⊗C2⊗B1) |
|  | *Amphiduros pacificus* | - | all | A2⊗J1, M⊗B1, D⊗G, + ‘spiralian’-fusions (J2⊗L⊗C2⊗I, H⊗Q⊗B2, O1⊗R⊗N) |
|  | *Aporrectodea icterica* | - | none | scrambled |
|  | *Bimastos eiseni* | - | none | scrambled |
|  | *Branchellion lobata* | - | none | scrambled |
|  | *Branchiopolynoe longqiensis* | - | all | M⊗I, C1⊗H, A1⊗E, + ‘spiralian’-fusions (J2⊗L⊗C2) |
|  | *Capitella teleta* | 10 | all | I⊗G, + ‘spiralian’-fusions (J2⊗L⊗C2) |
|  | *Dimorphilus gyrociliatus* | 12 | missing: K, A2, M, R, O2, C2 | NxI, A1xH, B2xB3, J1⊗F, B1⊗D, O1⊗G, C1⊗Q, P⊗E<br>‘spiralian’-fusions (J2⊗L only) |
|  | *Harmothoe impar* |  | all | A1⊗E, + ‘spiralian’-fusions (J2⊗L⊗C2) |
|  | *Helobdella robusta* | 9 | none | scrambled |
|  | *Lamellibrachia satsuma* |  | all | A2xM, A1xD, B2⊗J1, D⊗P, + ‘spiralian’-fusions (O1⊗R⊗B3 and J2⊗L⊗C2) |
|  | *Lepidonotus clava* | - | all | A1⊗E, + ‘spiralian’-fusions (J2⊗L⊗C2) |
|  | *Lumbricus rubellus* | - | missing: H, L, C2, J1, B2, C1, P, I, O2, R, F, M, G, A1, A2, Q, N, B3, B1, E, O1, D, J2, K | scrambled |
|  | *Metaphire vulgaris* | - | none | scrambled |
|  | *Owenia fusiformis* | 12 | all | C1⊗A2, B1⊗B3⊗C2, I⊗E, + ‘spiralian’-fusions (O1⊗R⊗M and J2⊗L⊗P⊗B2⊗J1) |
|  | *Paraescarpia ochinospica* | - | all | ExB1, A2xM, B2⊗J1, D⊗P, + ‘spiralian’-fusions (O1⊗R⊗B3 and J2⊗L⊗C2) |
|  | *Piscicola geometra* | 16 | none | scrambled |
|  | *Platynereis dumerilii* | 14 | all | C1⊗F, A1⊗E, + ‘spiralian’-fusions (H⊗Q⊗G and J2⊗L⊗O2⊗K⊗C2⊗B1) |
|  | *Protula sp. h YS-2021* | - | all | C1⊗A1, B1⊗B2, I⊗A2, B3⊗J1⊗G, D⊗E, + ‘spiralian’-fusions (J2⊗L⊗C2⊗P, O2⊗K⊗F) |
|  | *Sipunculus nudus* | - | all | J1⊗M, I⊗E, + ‘spiralian’-fusions (J2⊗L⊗C2) |
|  | *Sthenelais limicola* | - | all | C1⊗M⊗B1, D⊗E, A1⊗G, J1⊗B3⊗N, + ‘spiralian’-fusions (J2⊗L⊗I⊗C2, H⊗Q⊗F⊗B2, O2⊗K⊗P) |
|  | *Streblospio benedictii* | 11 | all | M⊗G, C1⊗B2, P⊗N, A1⊗B3, A2⊗D, + ‘spiralian’-fusions (O1⊗R⊗E, O2⊗K⊗J1, J2⊗L⊗C2⊗F) |
|  | *Terebella lapidaria* | - | all | G⊗F + ‘spiralian’-fusions (J2⊗L⊗C2, H⊗Q-P-E* ‘mixing-without-fusion’) |
| Brachiopods | *Lingula anatina* | 10 | all | Only the J2⊗L and H⊗Q spiralian fusions |
| Bryozoans | *Bugulina stolonifera* | - | missing: A2, B1, R, B2 | C2⊗A1, C1⊗D⊗G⊗E⊗B3, N⊗F, I⊗L⊗O1⊗Q (only J2⊗L⊗J1⊗I⊗O1⊗Q and O2⊗K⊗P ‘spiralian’-fusions) |
|  | *Membranipora membranica* | - | missing: J1, B2, C2, R, A2, B1, B3 | C1⊗D⊗G⊗E, N⊗F, I⊗L⊗H⊗Q (only the O2⊗K, H⊗Q⊗P ‘spiralian’-fusions) |
|  | *Cryptosula palasiana* |  | missing: R, B2, O1, B1, J2, B3, A2 | D⊗G⊗E⊗C1, C2⊗A1, I⊗L, N⊗F (only the O2⊗K⊗P, H⊗Q, ‘spiralian’-fusions) |
|  | *Watersipora subatra* | - | missing: A2, B1, B2, J1, R | C1⊗D⊗G⊗E⊗B3, N⊗F, C2⊗A1, O1⊗H, I⊗L⊗Q (only the L⊗J2⊗I⊗Q⊗O1, O2⊗K⊗P ‘spiralian’-fusions) |
| Mollusks | *Acanthopleura granulata* | - | all | D⊗P, C1⊗F, + ‘spiralian’-fusions (O1⊗R⊗I) |
|  | *Anadara kagoshimensis* |  | all | P⊗B3 + ‘spiralian’-fusions |
|  | *Crassostrea virginica* | 10 | all | B1⊗C2, M⊗A2⊗B3⊗F, C1⊗A1, I⊗N⊗D⊗G⊗P + ‘spiralian’-fusions (O1⊗R⊗B2⊗J1, H⊗Q⊗J2⊗L) |
|  | *Doryteuthis pealeii* |  | missing: J2, B3, O2, R | scrambled (J1⊗L, C2⊗I, D⊗G, D⊗A2, K⊗M, G⊗A1, G⊗F, E⊗H, L⊗F) |
|  | *Gari tellinella* |  | all | A2⊗B2, + ‘spiralian’-fusions |
|  | *Mytilus edulis* |  | all | D⊗A2, N⊗A1, I⊗C1, M⊗B1⊗B2, + ‘spiralian’-fusions (J2⊗L⊗E) |
|  | *Octopus sinensis* |  | C1, D, G, I, M, P, N, B1, B2, A1 and F | no ‘spiralian’-fusion |
|  | *Pomacea canaliculata* | 14 | all | G⊗J1, P⊗B2⊗C2, I⊗B1, + ‘spiralian’-fusions (H⊗Q⊗B3, O1⊗R⊗F) |
|  | *Ruditapes philippinarum* | 19 | all | A2⊗B2, + ‘spiralian’-fusions |
| Nemerteans | *Lineus longissimus* |  | all | C1⊗G, + ‘spiralian’-fusions |
|  | *Notospermus geniculatus* | 19 | all | C1⊗G, + ‘spiralian’-fusions (J2⊗L) |
| Phoronids | *Phoronis australis* |  | missing:<br>C2, J2, K, B2,<br>F, Q, J1, R and<br>O2 | fragmented assembly |
| Platyhelminthes | *Clonorchis sinensis* | 28 | missing: H, B2,<br>C1, C2, R, O2,<br>I, G, M, F, Q,<br>B1, K, J2, O1,<br>E | scrambled (A1⊗N, B3⊗A2⊗J1, L⊗D) |
|  | *Echinococcus granulosus* | 9 | missing: B1,<br>B3, N, E, O1,<br>J2, Q, A2, P, I,<br>R, O2, G, M,<br>H, L, J1, B2,<br>C2 | scrambled (K⊗D) |
|  | *Hymenolepis microstoma* | 6 | missing: G, I, P,<br>R, O2, J1, B2,<br>C2, H, L, K,<br>O1, E, J2, D,<br>B1, N, B3, Q,<br>A2 | scrambled + (A1⊗M) |
|  | *Schistosoma japonica* | 8 | missing: J2, E,<br>B1, B3, Q, A2,<br>G, M, R, O2, I,<br>P, C1, B2, J1,<br>C2, H | scrambled |
|  | *Schmidtea mediterranea* | 4 | none | scrambled |
|  | *Taenia multiceps* | - | K, O1, A1 and<br>F | scrambled |
| Rotifers | *Adineta vaga* | 6 | none | scrambled |
|  | *Brachionus rubens* | - | I, M, A2, N,<br>B2, B3, A1 and<br>F | A2⊗N⊗B2⊗B3, I⊗M |
|  | *Proales similis* | - | missing: A2,<br>B3, N, B1, O1,<br>D, J2, K, H, L,<br>C2, J1, P, O2,<br>R, G | scrambled |

### BLG configurations towards and within Errantia

Consistent with previous findings in other animal lineages, the majority of annelid genomes showed retention of the BLG complement with a high degree of chromosome conservation. Clitellata are an exception (n = 7) (Table 6; Supplementary Fig. 9) with BLG ‘scrambling’ (Fig. 8C; Supplementary Fig. 9), consistent with earlier sequencing results and recent studies (Lewin et al., 2024; Schultz et al., 2024; Simakov et al., 2013; Vargas-Chávez et al., 2024). For the non-clitellate species with clear BLG conservation we annotated additional BLG fusions-with-mixing events. For example, most annelids (except clitellates) possessed a J2⊗L⊗C2 fusion-with-mixing (Table 6, Fig. 8C), with only *Owenia* lacking this specific event (Fig. 8C), which supports *Owenia* (and Paleoannelida) representing the sister group to other annelids (note that J2⊗L alone represents a shared spiralian fusion (Simakov et al., 2022)). We also noted that *Dimorphilus* did not show statistically significant enrichment of the C2 BLG associated genes, likely due to faster evolutionary rates in genes and chromosomes in *Dimorphilus* (Table 6, Supplementary Fig. 9B).

We found a potential synapomorphic fusion-with-mixing event A1⊗E within errant annelids including Nereididae (n = 2) and Polynoidae (n = 5), belonging to the taxonomic order Phyllodocida. We note however that Hesionidae (n = 1) and Sigalionidae (n = 1), both thought to belong to the same taxonomic order of Phyllodocida (Weigert & Bleidorn, 2016), lacked this fusion-with-mixing event, suggesting placement of Sigalionidae outside the group comprising Nereididae and Polynoidae. Alternatively, A1⊗E could have fused independently in Nereididae and Polynoidae. Corroborating this, we did identify several potential independent fusion-with-mixing events on the annelid tree. For instance, the two lineages Oweniidae (n = 1) and Sipuncula (n = 1) both showed a I⊗E fusion-with-mixing event (Fig. 8C; Table 6). Between Hesionidae (n = 1) and Serpulidae (n = 1), the D⊗E fusion is shared; however this is only a single fusion character that unites two distinct lineages belonging to Errantia and Sedentaria (Weigert & Bleidorn, 2016), thus it has higher chance of being a convergent fusion. Within Sedentaria, Dinophilidae (n = 1) and Terebelliformia (n = 1) shared the P⊗E fusion event, despite belonging to distinct lineages (Fig. 8C). Single convergent fusion-with-mixing events, while rare, are not unlikely to be observed at such vast evolutionary distances (Schultz et al., 2023). Within Errantia, *Platynereis* and its relatively closely related genus *Alitta* showed several shared fusion-with-mixings, including C1⊗F, A1⊗E, and addition of several other fusions on-top of the pre-existing ‘spiralian’-fusions (G to H⊗Q, and O2⊗K⊗C2⊗B1 to J2⊗L). This suggests that the reduced chromosomal complement within this taxon can be at least partially explained by these chromosomal fusions.

## Conclusion

Despite its prominent research history, *Platynereis* has so far been lacking an adequate genome resource. Advances in sequencing technology and computational sequence analysis have been catalysts in achieving the goal of building the current resource. Together, the data and analyses presented, provide the reference genome assembly and annotation of a highly sought after spiralian model species and its two sibling species. While in the draft state, the assembly will clearly benefit from further chromosomal-scale sequencing efforts of closely related populations and species to disentangle the high genetic variation and produce a usable linkage map. This will in turn allow for the study of its evolutionary resilience and change at the cell-type level.

## Materials and Methods

The laboratory culture reference version of the *P. dumerilii* genome assembly is available at Genbank under accession number GCA_026936325.1 (WGS Project: JAPTHN01), single-individual assembly is available under accession number GCA_043381215.1. *P. megalops* and *P. massiliensis* genome assemblies from single individuals are available under Genbank accessions GCA_043380565.1 and GCA_043380595.1, respectively.

### DNA extraction and sequencing

For the polymorphic *Platynereis dumerilii* strain genome assembly (GCA_026936325.1 (WGS Project: JAPTHN01)) thousands of sibling worms from 2 fertilizations at the Özpolat Lab cultures in Woods Hole, MA, USA were used. These cultures originated from the polymorphic strains from Guillaume Balavoine’s laboratory in France. Worms were starved for 6 days in their culture boxes, then transferred into new boxes with clean sea water and starved for another 3 days. Worms were then collected into sterilized beakers, washed with about 2 L of 0.22 um filter-sterilized sea water, and transferred into 15 mL falcon tubes after washing. Excess water was removed as much as possible, samples were weighed and then flash frozen in liquid nitrogen for 2 minutes (tube 1 weighed 10.3 grams in 3 mL volume, tube 2 weighed 9.1 grams in 2 mL volume and tube 3 weighed 11.1 grams in 4 mL volume). Tubes were immediately transferred onto dry ice and shipped overnight to Dovetail Genomics (currently called Cantata Bio). Cantata Bio processed *P. dumerilii* samples for PacBio Sequel II CLR DNA sequencing.

An additional *P. dumerilii* sample was prepared for both Illumina sequencing using *P. dumerilii* cultured worms in Heidelberg, Germany. We used a protocol adapted from Oxford High Molecular Weight (HMW) extraction protocol from The Jackson Laboratory. Briefly, a single *P. dumerilii* sexually mature male (post sperm release) was incubated in 10mL lysis buffer (5M NaCl, 1M Tris pH8.0, 0.5M EDTA pH8.0, 10% SDS, 100mg/mL RNAse, Nuclease free H_2_O) for 1 hour at 37°C. 50-500 µL of Proteinase K was added to the lysis buffer and after mixing, sample was incubated at 50°C overnight (∼12-16 hours). DNA precipitation and elution were performed using the QIAGEN MaXtract High Density protocol kits (Cat. No. / ID: 129073). DNA was eluted in 300uL of TE pH8.0 buffer and stored at 4°C or −80°C, for long-term storage. All our DNA Illumina sequencing was performed in EMBL, Heidelberg at the Genomics Core Facility. For the genome size estimates and polishing Illumina data, DNA was extracted from a single ‘wild-type’ lab culture male (after sperm release) using the HMW QIAGEN Phenol-Chloroform kits. Approximately 500 nanograms (ng) – 1 microgram (µg) of DNA was used for library preparations and quality control. This sample was then sequenced using two HiSeq4000 lanes with 150 PE (thus a total of 300 bases). All protocols yielded good quality DNA at A260/230 and A260/230 ratios greater than 1.8.

### Genome size measurements

We estimated genome size using k-mer distributions and DNA quantity. To calculate k-mer distributions from Illumina 150PE sequenced (see below) data, we first trimmed high-quality reads based on length using Trimmomatic (Bolger et al., 2014) (command/parameters used ‘*java -jar trimmomatic-0.38.jar PE-threads 16 illumina_reads_R1.fastq illumina_reads_R2.fastq paired_R1.fq unpaired_R1.fq paired_R2.fq unpaired_R2.fq LEADING:3 TRAILING:3 SLIDINGWINDOW:4:15 MINLEN:105*’). Although the data was of high quality prior to trimming (evaluated using FastQC (Andrews, 2010)), we nevertheless trimmed data to ensure that only the best quality reads were retained for analysis. Duplicate reads were then discarded using SuperDeduper (Petersen, 2015) – re-implemented as part of – command/parameters: ‘*hts_SuperDeduper −1 trimmed_paired_R1.fq −2 trimmed_paired_R2.fq -f deduped*’). Deduplicated reads were further normalized (k-mer coverage to 100 times) using BBNorm from the BBTools suite (version 37.68; command/parameters ‘*bbnorm.sh prefilter=t usejni=t in=deduped_R1.fastq in2=deduped_R2.fastq out=normalized_R1.fq out2=normalized_R2.fq target=100 min=5*’). Counting of k-mers, size 21bp, was performed with Jellyfish version 2.2.7 (Marçais & Kingsford, 2011) using the commands: ‘*jellyfish count -t 16 -C -m 21 -s 6G -o 21mer_out --min-qual-char=? normalized_R1.fq normalized_R2.fq*’ followed by ‘*jellyfish histo -o 21mer_out.histo 21mer_out*’. We used Smudgeplot (version 0.2.1) and GenomeScope (version 2.0) (Ranallo-Benavidez et al., 2020) to estimate ploidy, heterozygosity and genome size respectively, from the normalized Illumina DNA-sequencing dataset. To estimate ploidy and heterozygosity, the commands ‘*smudgeplot.py cutoff 21mer_out.histo L*’ and ‘*smudgeplot.py cutoff 21mer_out.histo U*’ were executed so as to determine the lower and upper coverage cutoffs/thresholds. ‘*jellyfish dump -c -L $L -U $U 21mer_out | smudgeplot.py hetkmers -o kmer_pairs*’ was the command used to extract heterozygous k-mer pairs. Plots were derived using ‘*smudgeplot.py plot kmer_pairs_coverages.tsv -o pdum*’. The GenomeScope command executed to estimate genome size: ‘*genomescope2 -i 21mer_out.histo -o output -k 21*’.

We also stained dissociated *Drosophila melanogaster* embryos and *P. dumerilii* male and female adults into nuclei and stained them with DAPI to estimate DNA content via FACS. Briefly, frozen *Drosophila* embryos (stored at −80°C) were resuspended in 500µL ice-cold lysis buffer (10mM Tris-Cl pH 8.0, 10mM NaCl, 0.2% IGEPAL CA-630 and 1X cOmplete PI) and carefully ground using pre-chilled metal pestles until embryos were completely lysed. After incubating on ice for 15 mins, the homogenized suspension was centrifuged at 2,000 x *g* at 4°C for 10 mins, and the supernatant discarded. Remaining pellet was washed at least once with 500µL ice-cold lysis buffer, and later with 500µL ice-cold PBTriton 0.1%. Nuclei were then mechanically extracted using a series of 20G and 22G syringes, carefully pipetting the mixture. Solution was left in 4°C for no more than a week for experiments. Nuclei were then counted using beads, and then an equivalent DAPI:cell number staining ratio was added so that all nuclei across different samples had equal amounts of DAPI stain. A similar approach was taken for the *P. dumerilii* samples with the exceptions that animals were homogenized using 250mM sucrose, 25mM KCl, 5mM MgCl_2_ 10mM Tris-HCl pH 8.0 and 0.1% Nonidet P40/IGEPAL solution. After homogenizing the samples followed by low-speed centrifugation at 100 x *g* for 1 min at 4°C, the supernatant was saved and washed at least once with ice-cold homogenization buffer and at least twice with ice-cold 1X PBS with centrifugation at 400-500 x *g* for 4-7 mins each. Nuclei suspensions in 1X PBS were then filtered using 40µM Flowmi strainers and 10µM filters.

### Genome assembly

Three different long-read genome assembly algorithms were tested against our PacBio Sequel II CLR data; CANU (Koren et al., 2017), FLYE (Kolmogorov et al., 2019) and wtdbg2 (Ruan & Li, 2020) (Mutemi, 2023). We compared all assemblies, and found that CANU returned the more contiguous and complete genome assembly (Mutemi, 2023). The specific parameters used for CANU (version 2.1) were: ‘*canu -minReadLength=1000 -minOverlapLength=500 -genomeSize=1g -pacbio pacbio_sequel_II_CLR_subreads.fastq.gz -useGrid=false*’. Iterative purging was performed using purge_dups (Guan et al., 2020) and purge_haplotigs (Roach et al., 2018) (Mutemi, 2023 for protocol details). Throughout the purging iterations, we ran BUSCO (Simão et al., 2015) (version 5.0.0) to evaluate likely genome completeness: ‘*busco -i canu_asm.fa --config config.ini -l metazoa_odb10 -m genome –long -c 24*’. The final assembly was subjected to LINKS scaffolding (Warren et al., 2015) (see Mutemi, 2023 and below for recipe details), followed by Illumina-read polishing via POLCA (Zimin & Salzberg, 2020) with the command ‘*polca.sh -a links_canu_16.scaffolds.fa -r “trimmed_paired_R1.fq trimmed_paired_R2.fq” -t 32 -m 1G*’; approximately ∼2 million substitution errors and ∼1.8 million insertion/deletion errors were corrected, giving a final consensus quality of 99.74. We upload the assembly to NCBI under the GenBank assembly ID GCA_026936325.1 (whole genome sequence [WGS] project JAPTHN01) to facilitate its further exploration, annotation and quality maintenance, in hopes that it can be a useful resource for a broader audience, and not only the *Platynereis* community.

### *k*-mer pairs and Hi-C scaffolding

After assembling and polishing the genome (contigs=1007, N50=2.4 and size=1.46 Gbp), we subjected the assembly through iterative scaffolding methods. Firstly we made use of LINKS (Warren et al., 2015), which searched for *k*-mers of length 25 bp in the genome assembly as well as the raw PacBio Sequel II CLR datasets. The k-mer pairs used for scaffolding the contigs from the raw reads were sampled at different intervals (ranging from 200 to 1 bp) and over several distances (spanning from 10 to 200 Kbp); ‘*LINKS -f asm.fa -s long_reads.txt -d 10000-100000 -t 200-1 -k 25 -b links_pdum -v 1*’ (see Mutemi, 2023 and (Warren et al., 2015) for detailed iterative protocol). The LINKS scaffolded genome was then subjected to Hi-C scaffolding via SALSA2 (Ghurye et al., 2019), with 10 iterations. The Hi-C plot (Fig. 2) was generated using the ‘convert.sh’ script in the SALSA2 pipeline and visualized using Juicebox (Robinson et al., 2018).

A single *P. dumerilii* Hi-C library was prepared by Dovetail Genomics. For this dataset, the Hi-C library was prepared from hundreds of progenies from two parents (one male ♂and one female ♀). Briefly, chromatin was fixed in place with formaldehyde in the nucleus and then extracted. Fixed chromatin was digested with *DpnII*, the 5’ overhangs filled in with biotinylated nucleotides, and then free blunt ends were ligated. After ligation, crosslinks were reversed, and the DNA purified from proteins. Purified DNA was treated to remove biotin that was not internal to ligated fragments. The DNA was then sheared to ∼350 bp mean fragment size and a sequencing library was generated using Illumina-compatible adapters. Biotin-containing fragments were isolated using streptavidin beads before PCR enrichment of the library. The library was sequenced on three different Illumina lanes; a HiSeqX platform and two more HiSeq4000 lanes.

### Repeat modeling and evolutionary analysis

We modeled and masked REs using RepeatModeler (version 2.02) (Flynn et al., 2020) and RepeatMasker (version) (Tarailo-Graovac & Chen, 2009), on the scaffolded genome. We first built a database of likely REs using Dfam release 3.3 (Storer et al., 2021) with the commands ‘*BuildDatabase -name pdum scaffolds.fasta*’ and ‘*RepeatModeler -database pdum -pa 6 -LTRStruct > run.out*’. REs were then masked using RepeatMasker (version 4.1.1) (Tarailo-Graovac & Chen, 2009) with the commands: ‘*RepeatMasker-lib pdum-families.fa -pa 10 -dir pdum_scaffolds_masked -xsmall -gff -cutoff 250 -xm scaffolds.fasta*’. The same commands were executed when re-annotating REs for the other annelid genomes. The RMouttobed.pl script (Kapusta, 2017) was used to convert the *.out file from the RepeatMasker command into a *.bed file. This file was then used to extract length statistics and compare inter- and intra-genic occupancy of repeat elements using custom R scripts.

Mapping of quality-trimmed RNA-seq reads (see below) to an annotated ‘repeat_element.gtf’ file was performed via STAR (version 2.7.1a) (Dobin et al., 2012), using the commands: ‘*STAR --runThreadN 24--runMode genomeGenerate --genomeDir Reindex/ --genomeFastaFiles pdum_scaffolds.masked.fa --sjdbGTFfile pdum_repeats.gtf --limitSjdbInsertNsj 2637930*’ followed by ‘*STAR --runThreadN 24--genomeDir Reindex/ --readFilesIn trimmedRNA_paired_R1.fq trimmedRNA_paired_R2.fq --quantMode TranscriptomeSAM GeneCounts*’. The ‘ReadsPerGene.out.tab’ output file was further analyzed for RE expression counts in different loci. We considered only those loci whose read counts amounted to greater than or equal to 15 (Chen et al., 2016).

### Gene modeling and annotation

Transfer-RNA genes were predicted using tRNAscan-SE (version 2.0.7) (Chan et al., 2019; Lowe & Eddy, 1997) using the commands: ‘*tRNAscan-SE -EHQ -o# -f# -m# -s# -a# -l# --detail --thread 16 -p pdum_v2_tRNA pdum_scaffolds.fasta*’ followed by ‘*EukHighConfidenceFilter --result pdum_v2_tRNA.out --ss pdum_v2_tRNA.ss -p pdum_v2_tRNAQC -o eukqualfilt_pdumv2_tRNAs*’. Ribosomal RNA genes were predicted *de novo* from the assembly using barrnap (version 0.9) (Seemann, 2018), via the commands: ‘*barrnap --kingdom euk --threads 6 --reject 0.1 scaffolds.fasta --outseq pdum_rRNA.fa*’. We further mapped partial *P. dumerilii* rRNA sequences previously identified (Hui et al., 2007), and found consistent ribosomal gene loci, with the exception of novel 5S rRNA genes found with the barrnap models. Short PE Illumina RNA-seq reads were quality trimmed using Trimmomatic (version 0.38) with the commands ‘*java -jar trimmomatic-0.38.jar PE -threads 16 pdum_totalRNA_R1.fqpdum_totalRNA_R2.fq trimmedRNA_paired_R1.fq trimmedRNA_unpaired_R1.fq trimmedRNA_paired_R2.fq trimmedRNA_unpaired_R2.fq LEADING:3 TRAILING:3 SLIDINGWINDOW:4:15 MINLEN:45*’. These reads were then mapped to the scaffolded and masked genome using STAR (version 2.7.1a) (Dobin et al., 2012) with the commands: *STAR --runThreadN 16--runMode genomeGenerate --genomeDir index/ --genomeFastaFiles pdum_scaffolds.fasta,* followed by *STAR --runThreadN 16 --genomeDir index/ --readFilesIn trimmed_paired_R1.fq trimmed_paired_R2.fq*. Nanopore and PacBio transcriptomic long-reads were mapped to the same genome using Minimap2 (version 2.17) (Li, 2018); ‘*minimap2 -ax splice:hq -uf pdum_scaffolds.fasta pacbio_isoseq.fa > pb.sam*’ and ‘*minimap2 -ax splice pdum_scaffolds.fasta nanopore.fq > ont.sam*’.

StringTie (version 2.1.7) (Kovaka et al., 2019; M. Pertea et al., 2015) was used to reconstruct transcripts from the sorted. BAM mapping files generated for both short- and long-reads; ‘*stringtie -o *.gtf*sorted.bam --conservative -p 16*’, using the ‘-L’ option for long-reads. Using GFF-Utilities (G. Pertea & Pertea, 2020), we grouped overlapping transcripts from the three datasets (i.e. Illumina PE, PacBio and Nanopore) into loci using ‘*gffcompare -i {gtf_list.txt}*’ and ‘*gffcompare gffcmp.combined.gtf*’ - repeated at least three times to completely collapse overlapping exons/transcripts. Transcripts were extracted via ‘*gffread -w pdum_transcripts.fa -g scaffolds.fasta gffcmp.combined.gtf*’ and BUSCO values were calculated (using the transcriptome mode) giving a BUSCO score of: C, 96.7% [S: 44.0%, D: 52.7%], F, 0.6% and M, 2.7% from a Metazoa list of 954 genes. To convert to the Ensembl Gene-Transfer-Format (GTF), the AGAT tool (version 0.5.1) command ‘agat_sp_ensembl_output_style.pl’ was used (Dainat, 2020). Transcripts were extracted from the genome *.gtf using TransDecoder (version 5.5.0) ‘util’ scripts: ‘*perl ∼/util/gtf_genome_to_cdna_fasta.pl pdum_ensembl_genome.gtf scaffolds.fasta > pdum_transcripts.fasta* and *perl ∼/util/gtf_to_alignment_gff3.pl pdum_ensembl_genome.gtf > pdum_transcripts.gff3*’. Protein-coding genes were then retrieved via TransDecoder (version 5.5.0) using ‘*TransDecoder.LongOrfs -t pdum_transcripts.fasta* and *TransDecoder.Predict -t pdum_transcripts.fasta*’. Finally a genome-based coding region annotated file was built using ‘*perl∼/util/cdna_alignment_orf_to_genome_orf.pl pdum_transcripts.transdecoder.gff3 pdum_transcripts.gff3 pdum_transcripts.fasta > pdum_transcripts.transdecoder.genome.gff3* ‘. In total, we modelled 166,199 mRNA transcripts likely originating from 69,573 loci. Of the 166,199 transcripts, 72,852 had multiple exons. 24,237 loci (of the 69,573) contained multiple transcripts with (∼ 2.4 transcripts/loci), suggesting that approximately 35% of *P. dumerilii* genes contain more than one isoform. From the 166,199 transcripts (incl. isoforms), 93,240 had predicted ORFs; amounting to a total of 28,985 likely protein-coding genes. Selecting the longest peptide sequences for each transcript with an ORF, resulted in a total of 78,322 sequences. We subsequently annotated these protein sequences using orthology- (i.e. EGGNOGMAPPER version 2.1.5 (Cantalapiedra et al., 2021; Huerta-Cepas et al., 2019)), revealing only 37,664 with annotations, and 5,418 protein isoforms having a unique annotation.

For miRNA annotation and analysis, several developmental samples were pooled and subjected to single-end sequencing at ∼80bp length. RNA was extracted from several *P. dumerilii* developmental stages (24, 36, 48, 72, and 144 hours post-fertilization [hpf]), a minimum of three replicates for each developmental stage, using the Direct-zolTM RNA MiniPrep (Cat. No.: R2050). Sequencing libraries were prepared with the NextFlex smRNA kit. The 3’ adapter sequence ‘TGGAATTCTCGGGTGCCAAGG’ was trimmed using AdapterRemoval (version 2.3.2) (Schubert et al., 2016) using the commands: ‘*AdapterRemoval --file1 pdum_total_smallRNAs.fq --adapter1 TGGAATTCTCGGGTGCCAAGG --output1 pdum_trim_smallRNAs.fq*’, with additional random bases appearing immediately 5’ and 3’ to the insert removed using ‘*seqtk trimfq -b 4 -e 4 pdum_trim_smallRNAs.fq > pdum_trim_clip_smallRNAs.fq*’. miRNAs were predicted using miRDeep2 (Friedländer et al., 2012) with the commands: ‘*bowtie-build pdum_scaffolds.fasta pdum*’ to generate the index, ‘*mapper.pl pdum_trim_clip_smallRNAs.fq -e -h -I -j -m -l 18 -p pdum -s pdum_all_filt_collapsed.fa-t pdum_all_collapsed_genome.arf -v*’ to process and map reads to the genome and ‘*miRDeep2.pl pdum_all_filt_collapsed.fa pdum_scaffolds.fasta pdum_all_collapsed_genome.arf none cte.fas none 2>report.log*’. This identified 587 miRNA ‘unique’ gene locations, of which many could be grouped into miRNA gene clusters. Conserved miRNA sequences were identified using MirMachine (Umu et al., 2023). In addition, each output sequence from the MirDeep2 prediction was searched in MirGeneDB 2.1. miRNA gene names were thus assigned according to the best hits (i.e., lowest E value and/or highest Bits score) and by comparison to correspondent miRNA gene families of related species. For each selected organism, miRNAs genome coordinates were downloaded from MirGeneDB 2.1 (mirgenedb.org/browse). Genes were then sorted according to their genomic position and considered to be clustered when they were within 50KB (Baskerville and Bartel, 2005).The same approach was used to identify the clusters in *P. dumerilii*, considering the genome location of each annotated miRDeep2-predicted miRNA. A table summarising the details of the conserved miRNA clusters in each species was generated (Supplementary Files).

### Gene-content evolution analyses

For protein-coding gene count and statistics (i.e. exon, intron etc.), *.gff3 files for *Helobdella*, *Capitella* and *Dimorphilus* were downloaded from the NCBI Assembly database. AGAT (version 0.5.1) was used to quickly test for .gff3 file compatibility and basic statistics (Dainat, 2020). The longest isoforms per gene locus were extracted via the AGAT and GFF-Utilities tools using ‘*agat_sp_keep_longest_isoform.pl -gff transcripts.fasta.transdecoder.genome.gff3 -o single_isoform.transdecoder.genome.gff3* followed by *gffread -w pdum_single_isoform.transdecoder.fa -g scaffolds.fasta single_isoform.transdecoder.genome.gff3*’. The calculated BUSCO completion for this single isoform/gene fasta file stood at C, 92.1% [S: 88.4%, D: 3.7%], F, 0.6% and M, 7.3% from a Metazoa list of 954 genes; missing ∼4.6% of genes from the original proteome file (see above). For the comparative analyses, the *Capitella*, *Helobdella* and *Dimorphilus* CDS *.fasta files were downloaded from the GenBank Assembly database. For the *Streblospio* genome, we first extracted the longest protein-coding isoform/gene from the *.gff3 file, and extracted the transcripts (similar to the workflow for *Platynereis*) and filtered the non-coding tRNAs using pyfaidx (commands: ‘*faidx streblospio_single_isoform.cds.fa -v-g “_nc_”*’. To identify orthologous groups, we used OrthoFinder (version 2.5.4) (Emms & Kelly, 2015) using the default settings commands: ‘*orthofinder -f metazoa_proteomes_folder*’. We annotated these orthology groups using eggNOG-mapper (version 2.1.5) (Cantalapiedra et al., 2021) for each species to further explore if these species-specific orthogroups fall into specific functional categories.

### Variation analyses

*P. dumerilii* animals were sampled from the different sites with guided experts from the local marine stations. RNA was collected and sequenced using Illumina PE, and only quality-trimmed-paired reads were used for further analyses. We sampled the datasets to 20 million read pairs using ‘*seqtk sample -s10 RNA_trimmed.paired.R.fq 20000000 > RNA_trimmed.paired.sampled.R.fq*’, and mapped the data using STAR (Dobin et al., 2012) (version 2.7.9a). Variants were identified using bcftools (version 1.13) (Li, 2011) using the commands ‘*bcftools mpileup -Ou -f scaffolds.fasta rna_aligned.sortedByCoord.out.bam | bcftools call -mv -Ob -o calls.bcf*’. Preliminary descriptions of variants were performed using the SnpEff & SnpSift tool (version 5.0) (Cingolani et al., 2012). A *P. dumerilii* specific database was built using the commands: ‘*java -jar snpEff.jar build -v pdumv2.0*’. SNP and In/Del quantification was summarized using the commands ‘*java -jar snpEff.jar eff pdumv2.0*’. To count number of SNPs/In(Del)s per gene, we prepared genome annotation and variant call *.BED files (from *.GFF3 and *.VCF respectively) using the BEDOPS (version 2.4.39) (Neph et al., 2012) ‘*convert2bed*’ and ‘*vcf2bed*’ commands. We then summarized the counts of SNPs and In(Del)s using the BEDTOOLS (version 2.30.0) ‘*annotate*’ command.

The various *P. dumerilii* samples’ transcriptomes were built using RNA-Bloom (version 1.3.1) (Nip et al., 2020) with the commands: ‘*rnabloom --left <site_platynereis_sample>.R1.fq –right <site_platynereis_sample>.R2.fq -rcr -ntcard -outdir <site_sample>*’. Transcriptomes were subsequently filtered for sequences greater than 1000 bp, and Orthofinder (version 2.5.4) (Emms & Kelly, 2015) was used to identify orthology groups and generate a ‘species’ tree using: ‘*orthofinder -f variants_transcriptome_folder -M msa -A mafft -T fasttree -d -S blast_nucl*’. To test which genes showed site-specific variants, the eggNOG-mapper (version 2.1.5) (Cantalapiedra et al., 2021; Huerta-Cepas et al., 2019) annotated genes - commands: ‘*emapper.py -m diamond --itype CDS -i pdum.fasta -o emapper_output --cpu 50*’ - were intersected with the positions at which variants were detected.

### Genome assembly for P. massiliensis and P. megalops

*P. massiliensis* adult samples were collected from Roscoff, France in 2020, whereas *P. megalops* samples were collected from Woods Hole at the Marine Biological Laboratory (MBL) USA, in the summer of 2021. *P. megalops*, sexually mature animals were cultured and assessed for sexual behaviors by Dr. B. Duygu Özpolat and her team, prior to concluding if they were *P. megalops* or *P. dumerilii*. A single male was used for DNA sequencing. *P. massiliensis* samples were collected by Dr. Conrad Helm and colleagues, and observed in the laboratory conditions to confirm (i.e. tubes with sexually mature females and fertilized embryos also encased in the same tubes).

The Omni-C library was prepared using the Dovetail® Omni-C® Kit according to the manufacturer’s protocol. Briefly, the chromatin was fixed with disuccinimidyl glutarate (DSG) and formaldehyde in the nucleus. The cross-linked chromatin was then digested *in situ* with DNase I. Following digestion, the cells were lysed with SDS to extract the chromatin fragments and the chromatin fragments were bound to Chromatin Capture Beads. Next, the chromatin ends were repaired and ligated to a biotinylated bridge adapter followed by proximity ligation of adapter-containing ends. After proximity ligation, the crosslinks were reversed, the associated proteins were degraded, and the DNA was purified then converted into a sequencing library using Illumina-compatible adaptors. Biotin-containing fragments were isolated using streptavidin beads prior to PCR amplification. The libraries were sequenced on an Illumina NextSeq P3 platform to generate ∼400 million for each library (multiplexed) 2 x 150 bp read pairs.

The *P. megalops* (version 1.0) and *P. dumerilii* (version 2.0) and the *P. massiliensis* (version 1.0) genomes were assembled using PacBio HiFi reads, sequenced at ∼50X read depth, assuming 1 Gbp genome size. For all PacBio HiFi assemblies, highly accurate consensus sequences were produced from the *.subreads.bam files using the Circular Consensus Sequencing workflow from the PacBio bioconda tools with the commands: ‘ccs *subreads.bam platynereis_spp_ccs.fastq.gz --min-rq 0.99--reportFile=ccs_report.csv --num-threads 96’. Adapter sequences and were removed from the consensus reads using *Cutadapt* (version 1.18) with the commands ‘cutadapt -b TAGAGAGAGAAAAGGAGGAGGAGGCAACAACAACAACTCTCTCTA -bATCTCTCTCAACAACAACAACGGAGGAGGAGGAAAAGAGAGAGAT -oplatynereis_spp_ccs.clean.fastq.gz platynereis_spp_ccs.fastq.gz --cores=50’. The consensus reads were then assembled via the *hifiasm* algorithm (**Cheng et al., 2021**) with the commands: ‘hifiasm -o platynereis_spp-hifi.asm -t 48 platynereis_spp_ccs.fastq.gz -l 3 -s 0.55’. Genomes were scaffolded first using either Hi-C (*P. dumerilii*) or Omni-C via the Arima and SALSA2 pipelines, as aforementioned (**Ghurye et al., 2019**). REs were modelled and the respective genomes masked as described earlier for *P. dumerilii*.

### Synteny and gene-content evolution analyses

A set of *Branchiostoma floridae* highly conserved protein-coding genes were searched for using BLAST+ (version 2.12.0) against the hard-masked genome versions (i.e. all REs converted to ‘N’ nucleotides). Nucleotide databases for the hard-masked annelid genomes were built using the commands ‘*makeblastdb-in <hard-masked.fasta> -parse_seqids -dbtype nucl -out <database_name>*’. The conserved *B. floridae* proteins were then searched using ‘*tblastn -query <B.floridae_proteins.pep> -db <database_name> -out<blast_output_name> -max_target_seqs 1 -outfmt 6 -evalue 1e-2 -num_threads 24*’. The blast output file was then parsed for synteny analysis using custom scripts; the BLAST+ output result was processed using the following ‘*perl makeMap_Blast.pl <blast.outfmt6.output>. > blast.outfmt6.output.chrom*’ ‘*perl prepMsynt2.pl blast.outfmt6.output.chrom. threeway_final.allmbh.clus > blast.outfmt6.output.3waymbh_final.msynt*’.

We accessed publicly available assembled annelid genomes (n = 22, plus our *P. dumerilii* assembly) (at either chromosome or near-chromosome resolution) available via the NCBI Assembly database (Materials and Methods). This included an annelid genome from the Oweniidae (*Owenia fusiformis*), which together with Magelonidae comprises the Paleoannelida – earliest major branch of Annelida – another from the Sipuncula lineage (*Sipunculus nudus*) which together with Amphinomidae form a sister group to the Pleistoannelida (Errantia and Sedentaria) (Weigert & Bleidorn, 2016). From the Errantia lineage, two Nereididae (*Alitta virens* and *Platynereis dumerilii*), five Polynoidae (*Acholoe squamosa*, *Alentia gelatinosa*, *Branchiopolynoe longqiensis*, *Harmothoe impar* and *Lepidonotus clava*), a Hesionidae species (*Amphiduros pacificus*) and a Sigalionidae species *Sthenelais limicola*, and a total of 12 Sedentaria with six from Clitellata (*Bimastos eiseni*, *Branchellion lobata*, *Helobdella robusta*, *Lumbricus rubellus*, *Metaphire vulgaris* and *Piscicola geometra*) and a single representative from each of Terebelliformia (*Terebella lapidaria*), Capitellidae (*Capitella teleta*), Spionidae (*Streblospio benedictii*), Serpulidae (*Protula*), Siboglinidae (*Paraescarpia*) and Dinophilidae (*Dimorphilus gyrociliatus*).

## Notes

### Competing Interest Statement

The authors have declared no competing interest.

### Summary of Updates

NCBI and GenBank accession numbers for genome assemblies. Added single individual sequencing results. Updated the results in the miRNA section. Updated methods. Updated Figure 1 Updated References

