## Supplementary File 03 for "A genome resource for the marine annelid *Platynereis dumerilii*"

### microRNA Annotation Compendium
### *Platynereis dumerilii*

| miRNA name | Mature sequence | Star sequence | pre-miRNA | Gene coordinate |
|---|---|---|---|---|
| Pdu-Bantam-PX_3p | UGAGAUCAUGGUGAAAACUAAU | ugguuuucacaauggucuauaucaga | ugguuuucacaauggucuauaucagaguauauucaaaacccugagaucauggugaaaacuaau | scaffold_2:72666494..72666557:- |
| Pdu-Bantam-PY_3p | UGAGAUCAUUGUGAAAACUGAUU | cugguuuucacauuggucuucc | cugguuuucacauuggucuuccagagugaguuucucgucugagaucauugugaaaacugauu | scaffold_2:72665913..72665975:- |
| Pdu-Let-7_5p | UGAGGUAGUAGGUUGUAUAGUA | cuguacaaccuucuagcuuucc | ugagguaguagguuguauaguaagagaacuacaccuuuuucuggcggacuguacaaccuucuagcuuucc | scaffold_4:25097122..25097192:- |
| Pdu-Mir-1_3p_+_ | UGGAAUGUAAAGAAGUAUGUAG | acauacuucuucauauaccaua | acauacuucuucauauaccauaucuuuacugauguauggaauguaaagaaguauguag | scaffold_1:162427564..162427622:+ |
| Pdu-Mir-1_3p-5p_-_ | UGGUAUAUGAAGAAGUAUGUGA | acauacuucuuuacauuccaua | acauacuucuuuacauuccauacaucaguaaagauaugguauaugaagaaguauguga | scaffold_1:162427562..162427620:- |
| Pdu-Mir-10-P1_5p | UACCCUGUAGAUCCGAAUUUGU | aaauucgagucugcggaguguug | uacccuguagauccgaauuuguguauaaaaugacaacaaauucgagucugcggaguguug | scaffold_27:3434438..3434498:+ |
| Pdu-Mir-10-P2_5p | AACCCGUACAACCGAACUUGUG | caagcucgcucuuacgggccug | aacccguacaaccgaacuugugucguacgaauaaaucacaagcucgcucuuacgggccug | scaffold_4:25098345..25098405:- |
| Pdu-Mir-10-P3_5p-3p | UCCCUGAGACCCUAACUUGUGA | acagguugaggucuugggacua | ucccugagacccuaacuugugaagaaaaacgucacagguugaggucuugggacua | scaffold_4:25095716..25095771:- |
| Pdu-Mir-10-P4-1_3p-5p_+_ | AGAAGCUCGGUUCUACAGGUA | acccuguagacccggguuugugu | acccuguagacccggguuuguguuguucagagcaaaucacagaagcucgguucuacaggua | scaffold_27:3633349..3633410:- |
| Pdu-Mir-10-P4-1_5p_-_ | UACCUGUAGAACCGAGCUUCUG | caaacccgggucuacaggguau | uaccuguagaaccgagcuucugugauuugcucugaacaacacaaacccgggucuacaggguau | scaffold_27:3633349..3633412:+ |
| Pdu-Mir-10-P4-2_5p | AACCCUGUGGAUACGGGCUUGU | aagcucguuuuuacaagguguc | aacccuguggauacgggcuugugucgauuacaucaacaagcucguuuuuacaagguguc | scaffold_27:3503211..3503270:- |
| Pdu-Mir-10-P5_5p | UUACCCUGUAGAACCGAGCGAGUA | ccacucaguucacagggucauu | uuacccuguagaaccgagcgaguauguauucauuaguaccacucaguucacagggucauu | scaffold_27:2892358..2892418:+ |
| Pdu-Mir-10-P6_5p | UUUACCCUGUGAAUAGAGAGUG | gucucauuucuuggggguaagca | uuuacccugugaauagagagugggguauuucauucccagucucauuucuuggggguaagca | scaffold_27:2135863..2135923:+ |
| Pdu-Mir-10-P7_5p | CACCCUGUAGAACCGAGCUUGU | caggcugguuacaccgggucaa | cacccuguagaaccgagcuugugcugaauucaaaacacaggcugguuacaccgggucaa | scaffold_27:3180449..3180508:+ |
| Pdu-Mir-1175_3p | UGAGAUUCAACUCCUCCAACUGC | aguggagagaguucuaucucauc | aguggagagaguucuaucucaucgucucaguucaaucuggugagauucaacuccuccaacugc | scaffold_1:113487497..113487560:+ |
| Pdu-Mir-12_3p-5p | CGUGCCUUUUGUGAUUCUCUUG | ugaguauuacaucagguacuga | ugaguauuacaucagguacugaguguuuauaagacguccgugccuuuugugauucucuug | scaffold_7:46912750..46912810:- |
| Pdu-Mir-124_3p-5p | UAAGGCACGCGGUGAAUGCCA | aguguucacuguguacgccuuggu | aguguucacuguguacgccuugguguguugguaacuauuaaggcacgcggugaaugcca | scaffold_1:63283530..63283588:- |
| Pdu-Mir-133_3p | UUGGUCCCCUUCAACCAGCUGU | agcugguugaaauagggccaaau | agcugguugaaauagggccaaauugucugauuuucugucauuugguccccuucaaccagcugu | scaffold_1:162444773..162444836:+ |
| Pdu-Mir-153_5p-3p | CGAGCUUUUGUGAUUUGCAAUU | uugcauagucacaaaagugauc | cgagcuuuugugauuugcaauuguauuguggacaaauugcauagucacaaaagugauc | scaffold_1:61020209..61020267:- |
| Pdu-Mir-184_3p | UGGACGGAGAACUGAUAAGGGC | ccuucucacuuguucguccggu | ccuucucacuuguucguccggucuuauauuugaguacuggacggagaacugauaagggc | scaffold_1:167014350..167014409:+ |
| Pdu-Mir-190_5p | AGAUAUGUUUGAUAUAUUUGGUGG | accauguaucaaacaugucacg | agauauguuugauauauuugguggguuuuuauaaaggaucuaccauguaucaaacaugucacg | scaffold_1:167624155..167624217:+ |
| Pdu-Mir-193-P2_3p | CAAUGCCCUAUGAAAUCCUAAA | ugggauuucugggggcauccugug | ugggauuucugggggcauccugugauauugaacugacaaugcccuaugaaauccuaaa | scaffold_186:387560..387618:- |
| Pdu-Mir-1986_5p-3p | CAAGGAUCCUGGGGAAGCCCCC | uggauuucccaugauccguaac | caaggauccuggggaagcccccuuccaucaaugguggauuucccaugauccguaac | scaffold_10:42047757..42047813:+ |
| Pdu-Mir-1987_3p-5p | ACUGCCAGAUGUCAUGUUGUGCA | cauaacuggcaccugggcauguug | cauaacuggcaccugggcauguugcucuucacucaacugccagaugucauguugugca | scaffold_1:66332418..66332476:- |
| Pdu-Mir-1989-1-a_5p | UCAGCUGUAAAGAUGCCUUCUU | ggagccauuuuaacugcugaca | ucagcuguaaagaugccuucuugaaauucauacaaggagccauuuuaacugcugaca | scaffold_10:38602058..38602115:- |
| Pdu-Mir-1989-2-a_5p | UCAGCUGUAAAGAUGCCUUCUU | ggagucauuuuaacugcugaca | ucagcuguaaagaugccuucuugaaauuuaaacaaggagucauuuuaacugcugaca | scaffold_10:38602337..38602394:- |
| Pdu-Mir-1989-3-a_5p | UCAGCUGUAAAGAUGCCUUCUU | gagacauuuuaacugcugacau | ucagcuguaaagaugccuucuugaaauuuacaaaaggagacauuuuaacugcugacau | scaffold_10:38602627..38602685:- |
| Pdu-Mir-1989-4-b_5p | UCAGCUGUAACGAUGCCUUCUU | ggagucaucuuaacugcugaca | ucagcuguaacgaugccuucuugaaauuuaaacaaggagucaucuuaacugcugaca | scaffold_10:38602893..38602950:- |
| Pdu-Mir-1989-5-c_5p | UCAGCUGUCACGAUGCCUUCUU | gggggcauuuuaacugcugaca | ucagcugucacgaugccuucuugucauuuaccgaagggggcauuuuaacugcugaca | scaffold_10:38603309..38603366:- |
| Pdu-Mir-1989-6-d_5p | UCAGCUGUCGCGAUGCCUUCUU | gaagguguuaccacugcugaca | ucagcugucgcgaugccuucuuuucacaauuucuagcaagaagguguuaccacugcugaca | scaffold_10:38603735..38603796:- |
| Pdu-Mir-1990-1orP1_3p_+_ | UGGGACUAUGUCAACUUACAAC | uggaaguuaacguagucccggg | uggaaguuaacguagucccggguguguuuauucccugggacuaugucaacuuacaac | scaffold_10:42046321..42046378:- |
| Pdu-Mir-1990-1orP1_5p-3p_-_ | UGUAAGUUGACAUAGUCCCAGG | cgggacuacguuaacuuccagc | uguaaguugacauagucccagggaauaaacacacccgggacuacguuaacuuccagc | scaffold_10:42046323..42046380:+ |
| Pdu-Mir-1990-2orP2_5p | UGUAAGUUAACGUAGUCCAAGGUU | uugagauuacgauaacuuuccau | uguaaguuaacguaguccaagguuuaugaaaggcaaacuugagauuacgauaacuuuccau | scaffold_18:7382654..7382715:+ |
| Pdu-Mir-1992-v1_5p-3p | CAUUAGUGGAUAACGGCUGGUA | ucagcaguuguaccacugaugug | cauuaguggauaacggcugguaguaacauuacgucugccuaucagcaguuguaccacugaugug | scaffold_12:23661448..23661512:- |
| Pdu-Mir-1992-v2_3p | CAUUAGUGGAUAACGGCUGGUA | cucagcacuaauccaucugug | cucagcacuaauccaucugugcgucuuccugacucuggcacuggcagcacauuaguggauaacggcuggua | scaffold_12:23661490..23661561:- |
| Pdu-Mir-1993_3p | UAUUAUGCUGUUAUUCACGAGA | ucgggaauaucaguguucuaugcc | ucgggaauaucaguguucuaugccguuuguugaagguauuaugcuguuauucacgaga | scaffold_1:57292928..57292986:- |
| Pdu-Mir-1995_5p-3p | GUACAUCUCACAUUGUGACCAU | ggucucugugugaaaugucgau | guacaucucacauugugaccaugugacuuaacaauggucucugugugaaaugucgau | scaffold_5:31347920..31347977:- |
| Pdu-Mir-1996-1 | CUCAAGUGAGGUCAGUGCUACU | gggcacugcccucaacugaagc | cucaagugaggucagugcuacuguucuugaaccaggggcacugcccucaacugaagc | scaffold_1:189982417..189982474:- |
| Pdu-Mir-1996-2 | AUCAAGUGAGGUCAGAUCUUGG | agggucauggucucaauugaccc | aucaagugaggucagaucuugggugacaugggggcgccagggucauggucucaauugaccc | scaffold_1:189983200..189983260:- |
| Pdu-Mir-1997_3p | UCUGCAGGUUCACAUCAGCCCCA | gggcgaugggguucucgcagggu | gggcgaugggguucucgcagggugagauaaaacuauucugcagguucacaucagcccca | scaffold_4:4379290..4379348:+ |
| Pdu-Mir-1998_3p | UUGAACGCAGAGAUGUACAUCA | auguauaucccuacguccacacu | auguauaucccuacguccacacuucccucuccaguugaacgcagagauguacauca | scaffold_2:43820342..43820398:+ |
| Pdu-Mir-2000_5p | UGACUAAAAGUAUUGAAGGCUU | gucuucacuacuuuuaguuugg | ugacuaaaaguauugaaggcuuuaagucaaaucucuguuaaagucuucacuacuuuuaguuugg | scaffold_2:43826989..43827053:+ |

| | | | | |
|---|---|---|---|---|
| Pdu-Mir-210-1_3p-5p | CUUGUGCGUGUGACAGUGACAAU | ccgucacuggcuccgcgcaaaga | ccgucacuggcuccgcgcaaagaguagaaugaugccccucuugugcgugugacagugacaau | scaffold_10:1472007..1472069:- |
| Pdu-Mir-210-2_3p | UUGUGCGUGGGACAGCGGCCCGC | caccguugcuucaugcccagcg | caccguugcuucaugcccagcguuaaauucaagucguugugcgugggacagcggcccgc | scaffold_10:1593967..1594026:- |
| Pdu-Mir-210-3_3p | UUGUGCGUGGAACAGCGACCAU | gucacuggccuuggcacaagca | gucacuggccuuggcacaagcacuuacguuuuaauguguugugcguggaacagcgaccau | scaffold_10:1599739..1599799:- |
| Pdu-Mir-216-P1_5p | UAAUCUCAGCUGGUAAAAGUGAG | ccaguuacuagcugagguuaca | uaaucucagcugguaaaagugagugcaugacuucuccaguuacuagcugagguuaca | scaffold_7:46914986..46915043:- |
| Pdu-Mir-216-P2_5p | AAAUAUCAGCUGGUAAUUCUGAG | caggauucgagcugauauuggg | aaauaucagcugguaauucugagugguuugaucuaguccucaggauucgagcugauauuggg | scaffold_7:46915860..46915922:- |
| Pdu-Mir-219-1_5p | UGAUUGUCCAAACGCAUUUCUUG | agaacuguuuuggacaucagu | ugauuguccaaacgcauuucuuguuguagcugaccaagaacuguuuuuggacaucagu | scaffold_6:22743322..22743380:+ |
| Pdu-Mir-219-2_5p | UGAUUGUCCAAACGCAUUUCUUG | agaacuguuuuggacaucagu | ugauuguccaaacgcauuucuuguuguagcugaccaagaacuguuuuuggacaucagu | scaffold_8:20782311..20782369:- |
| Pdu-Mir-22-P1_3p | AGCUGCCUGGUGAAGAGCUGUC | cggccuuucuccaggcaggacuuu | cggccuuucuccaggcaggacuuuguguuucugacuaaagcugccuggugaagagcuguc | scaffold_2:75490842..75490902:- |
| Pdu-Mir-22-P2_3p-5p | GAGCUGCCAAGUGAAGGGCUGU | aguuccuucuccuggcugcugucccc | aguuccuucuccuggcugcugucccuaugcuaugggcgggagcugccaagugaagggcugu | scaffold_2:75489903..75489964:- |
| Pdu-Mir-242-1a_5p | UUGCGUAGGCGUUGUGCACAGA | uguguauuuugccuacgcaaauu | uugcguaggcguugugcacagauuugaucacgcgagugggucuguguauuuugccuacgcaaauu | scaffold_7:29826410..29826474:+ |
| Pdu-Mir-242-2b_5p | UUGCGUAGGCGUUGUGCACAGA | uguguauuuuaccuacgcaaau | uugcguaggcguugugcacagauuugaucaugaagguggucuguguauuuuaccuacgcaaau | scaffold_7:29826776..29826839:+ |
| Pdu-Mir-242-3c_5p | UUGCGUAGGCGUUGUGCACAGA | uguguauuuugccuacgcaauu | uugcguaggcguugugcacagauuugaccaacuaagugggucuguguauuuugccuacgcaauu | scaffold_7:29827091..29827154:+ |
| Pdu-Mir-242-4a_5p | UUGCGUAGGCGUUGUGCACAGA | uguguauuuugccuacgcaaauu | uugcguaggcguugugcacagauuugaucacagaggugguucuguguauuuugccuacgcaaauu | scaffold_7:29836783..29836847:+ |
| Pdu-Mir-252-P1_5p | CUAAGUACUAGCGCCGCAGGA | cugcuguccuagugcuuaaug | cuaaguacuagcgccgcaggaguuccuuuuuaagccuccugcuguccuagugcuuaaug | scaffold_5:66482390..66482449:- |
| Pdu-Mir-252-P2_5p | CUAAGUAGUAGCGCCGCAGGUA | ccugcacccugcugcuuauca | cuaaguaguagcgccgcagguaguugucaacuauaccugcacccugcugcuuauca | scaffold_5:66483205..66483261:- |
| Pdu-Mir-2685_3p | UAACUCAGUCAGAUCACAGUC | cagugguauggcucgaguugua | cagugguauggcucgaguuguagagacguaacauauaacucagucagaucacaguc | scaffold_30:5202777..5202833:+ |
| Pdu-Mir-2689_3p | UAUCCUGGCCUGCAAGUGCACA | ugcauucgcaugucuggaugca | ugcauucgcaugucuggaugcaauuacauuugcauuuguauccuggccugcaagugcaca | scaffold_49:2560194..2560254:- |
| Pdu-Mir-2691-1_3p | AUUUGCAAUGUAUCACUGCCCG | ggcgaugucacauugcagucg | ggcgaugucacauugcagucgugugugguuuaaaaucgauuugcaauguaucacugcccg | scaffold_12:16062900..16062959:+ |
| Pdu-Mir-2691-2_3p | AUUUGCAAUGAAUCACUGCCCA | gguggcgauucuuugcagucg | gguggcgauucuuugcagucguguaaaauggaagacgauuugcaaugaaucacugccca | scaffold_12:26857282..26857341:+ |
| Pdu-Mir-2691-3_3p | AUUUUGCAUCGUAUCACUGCCGG | gguagcgauucccugacaaaaucu | gguagcgauucccugacaaaaucuucauauaguuagauuuugcaucguaucacugccgg | scaffold_27:5652967..5653026:- |
| Pdu-Mir-2691-4_3p | GUUUUGCAUGGUAUCACUGCUC | guagugauacuuugcacaaaucg | guagugauacuuugcacaaaucguaaagaaacaucgguuuugcaugguaucacugcuc | scaffold_44:156337..156395:+ |
| Pdu-Mir-2691-5_3p | GACGGUGAAACUUGUGCAAAACC | uuuugcaaaguaucacagcuua | gacggugaaacuugugcaaaaccuuacauccaguugguuuugcaaaguaucacagcuua | scaffold_4:75689505..75689564:- |
| Pdu-Mir-2692_3p | CCAGUCAAUGUUGACACCACCGCU | ggugaugucccauugacuuuggug | ggugaugucccauugacuuuggugggauuuuguaaaacaccagucaauguugacaccaccgcu | scaffold_10:42044737..42044800:+ |
| Pdu-Mir-2705-1_3p | UCUGCAAGGUAAGUGCUGUCU | gcggugcuuaccuuugcagcug | gcggugcuuaccuuugcagcugaugauuugccaucugcaagguaagugcugucu | scaffold_1:66337315..66337369:- |
| Pdu-Mir-2705-2_3p | UCUGCAACGUAAGUGCUGUCA | acagcgcuaugccuuuauugcaacc | ucugcaacguaagugcugucauggaaacagcgcuaugccuuuauugcaacc | scaffold_6:5069891..5069942:+ |
| Pdu-Mir-2707_3p-5p | AAUACUUAUUUUGCUUCUGACAGA | ugucaagcagauuaaguauugu | ugucaagcagauuaaguauuguugguaauggaauaaaauacuuauuuugcuucugacaga | scaffold_26:3813112..3813171:+ |
| Pdu-Mir-277_3p | UAAAUGCAUUAUCUGGUAUGUA | cauaucagaaaugcacuuugca | cauaucagaaaugcacuuugcauguuacuauaaaaguguaaaugcauuaucugguaugua | scaffold_4:52020820..52020880:+ |
| Pdu-Mir-278_3p | UCGGUGGGACUUUCGUUCGUUC | acgaaugaaauucucgacaggguc | acgaaugaaauucucgacagggucgauucacaagcgaucgguggggacuuucguucguuc | scaffold_2:55309987..55310046:+ |
| Pdu-Mir-279-1a_3p | UGACUAGAUCCACACUCAUCC | augacuguaggucuaguccaug | augacuguaggucuaguccauguuuaaugucaugacuagauccacacucaucc | scaffold_9:2657710..2657763:- |
| Pdu-Mir-279-1b_3p | UGACUAGAUCCACACUCAUCC | ggugagugggauuuuggucaug | ggugagugggauuuuggucauguuuuuuaauaucuucaugacuagauccacacucaucc | scaffold_1:184772430..184772489:- |
| Pdu-Mir-279-2a_5p-3p | AUGACUGUAGGUCUAGUCCAUG | ugacuagauacacacucaucc | augacuguaggucuaguccauguuuaaugucaugacuagauacacacucaucc | scaffold_3:17716402..17716455:+ |
| Pdu-Mir-279-2b_5p | AUGACUGUAGGUCUAGUCCAUG | ugauuagucaaacacucauagc | augacuguaggucuaguccauguuuaauaucaugauuagucaaacacucauagc | scaffold_3:90160574..90160628:- |
| Pdu-Mir-281_3p-5p | UGUCAUGGAGUUGCUCUCUUUA | aagggagcauucuuggacagu | aagggagcauucuuggacaguggucaaucaaaaacugucauggaguugcucucuuua | scaffold_22:3227625..3227682:- |
| Pdu-Mir-29-P1_3p | UAGCACCAUUUGAAAUCAGUUU | gcuggguucuucuggugcugga | gcuggguucuucuggugcuggauaccauaacucuagcaccauuugaaaucaguuu | scaffold_11:27895590..27895645:- |
| Pdu-Mir-29-P2_5p-3p | CUGGUCUCAAGUGGUGGAUAGA | uagcaccauuugaaaucagu | cuggucucaagugguggauagaggucugaaaacugucuagcaccauuugaaaucagu | scaffold_11:27907196..27907253:- |
| Pdu-Mir-2-oAa1_3p | UAUCACAGCGUGCUUUGAUGAGCU | cucgcgaagucgcugugccagg | cucgcgaagucgcugugccagguuuuaacgaguccuaucacagcgugcuuugaugagcu | scaffold_3:67088839..67088898:+ |
| Pdu-Mir-2-oBb1_3p-5p | UAUCACAGCAGCGCUUUGAUGAGUC | ggcaucaaggguucugugaaaug | ggcaucaaggguucugugaaaugcugaauuguaccauaucacagcagcgcuuugaugaguc | scaffold_3:67088402..67088463:+ |
| Pdu-Mir-2-oCc-a1_3p | UAUCACAGCCAGCUUUGAUGAGC | cccaucuuaguggcugugggugug | cccaucuuaguggcuguggugugcauuuaugccgacauaucacagccagcuuugaugagc | scaffold_3:67088097..67088157:+ |
| Pdu-Mir-2-oDc-b1_3p | UAUCACAGCCAGCUUUGAUGAGC | ccaucuaaguggcugugaugug | ccaucuaaguggcugugauguggguguugcgaugcauaucacagccagcuuugaugagc | scaffold_3:67087900..67087958:+ |
| Pdu-Mir-2-oEd1_3p | UAUCACAGCCAGCUUUGACGAGC | cgucacaguggcugugaagug | cgucacaguggcugugaagugguuagaugaacgcauaucacagccagcuuugacgagc | scaffold_3:67087490..67087548:+ |
| Pdu-Mir-2-oFe1_5p | UAUCACAGCCUUGCUUGGAUCAUA | cgguccaagugguugagauagc | cgguccaagugguugagauagcuguuuucucuaaacuggggguaucacagccuugcuuggaucaua | scaffold_3:67087095..67087160:+ |
| Pdu-Mir-2-oGc-c1_3p | UAUCACAGCCAGCUUUGAUGAGC | caucaauccuggcauguuaua | caucaauccuggcauguuauaguauccacagucgagcuaucacagccagcuuugaugagc | scaffold_3:67086276..67086336:+ |
| Pdu-Mir-2-oA2_5p-3p | UGGCCAAGGGGGCUGUGAUGUG | uaucacagcccaugcuuuggucaaa | uggccaagggggcugugauguguugaaaugcgucauaucacagcccaugcuuuggucaaa | scaffold_3:66418561..66418621:+ |
| Pdu-Mir-2-oB2_3p | UAUCACAGCCCGCUUUGACAAGU | guguugaagugguugugaugug | guguugaagugguugugauguguuguacuaguguccauaucacagcccgcuuugacaagu | scaffold_3:66418750..66418810:+ |
| Pdu-Mir-2-oC2_3p | UAUCACAGCCAAUUUGACAGCC | cuguuggauuuguugugacuaug | cuguuggauuuguugugacuauguauaaggcaucauaucacagccaauuugacagcc | scaffold_3:66419071..66419128:+ |
| Pdu-Mir-2-oD2_3p | UAUCACAGCCCGCUUUGGUCAUA | cggucaaaguggcugagauaug | cggucaaaguggcugagauauguuuuuguguucguaucacagcccgcuuuggucaua | scaffold_3:66419294..66419351:+ |
| Pdu-Mir-2-oE2_3p | UAUCACAGCCAGAUUUGGUAUAUC | agccaaaucggcugugaaaug | agccaaaucggcugugaaauguuauaucaacugagucguaucacagccagauuugguauauc | scaffold_3:66419578..66419640:+ |
| Pdu-Mir-31_5p | AGGCAAGAUGUUGGCAUAGCUGA | agcucugucccauguugccacc | aggcaagauguuggcauagcugagucacguggcgcgucagcucugucccauguugccacc | scaffold_1:92405032..92405092:+ |
| Pdu-Mir-315_5p | UUUUGAUUGUUGCUCAGAAAGCC | cuaucggguaguaaucaaaaaa | uuuugauuguugcucagaaagccguugugacugacuggcuaucggguaguaaucaaaaaa | scaffold_11:16298118..16298178:+ |
| Pdu-Mir-317_3p | UGAACACAGCUGGUGGUAUCUCUUU | agaguacuccagcguggucgca | agaguacuccagcguggucgcauguuguuaaagugugaacacagcugguggguaucucuuu | scaffold_4:52017422..52017482:+ |
| Pdu-Mir-34_5p | UGGCAGUGUGGUUAGCUGGUUGU | aaccacuaucugcccugucuua | uggcaguggguuagcugguugugugucgcguagaaauguuacaaccacuaucugcccugucuua | scaffold_4:52046695..52046759:+ |
| Pdu-Mir-36_5p-3p | GGUGAGUGGAUACUCAGGUGGUG | ucaccggguauacauucauccg | ggugaguggauacucaggugguguuccaauugacgucaucaccggguauacauucauccg | scaffold_9:2652939..2652999:- |
| Pdu-Mir-375_3p | UUUGUUCGUCCGGCUCGCGUUA | augugagcuauucguacagagc | augugagcuauucguacagagcuuauuucucuugcuuuguucguccggcucgcguua | scaffold_31:4109488..4109545:- |

| | | | | |
|---|---|---|---|---|
| Pdu-Mir-67_3p-5p | ACCUUAUUCAGUGGUGUUGUGGU | ucacaaccugcaugaaugaggua | accuuauucagugguguuguggugagguuaaaaagcaucacaaccugcaugaaugaggua | scaffold_2:34906472..34906532:+ |
| Pdu-Mir-7_5p | UGGAAGACUAGUGAUUUUGUUGUU | caaugaaacacuaucuuccaua | uggaagacuagugauuuuguuguuguucgcugaccuaacaaugaaacacuaucuuccaua | scaffold_1:24169743..24169803:- |
| Pdu-Mir-71_5p | UGAAAGACAUGGGUAGUGAGAUG | gcucgcuacccugucuuuccag | ugaaagacauggguagugagauggagagaaagaaacagcucgcuacccugucuuuccag | scaffold_3:67085297..67085356:+ |
| Pdu-Mir-750_3p | CCAGAUCUAACUCUUCCAGCUCA | cgcuggaagcuuaggucuccgca | cgcuggaagcuuaggucuccgcagugucauuccugccagaucuaacucuuccagcuca | scaffold_1:113485471..113485529:+ |
| Pdu-Mir-76_3p-5p | UUCGUUGUCGUCGAAACCUGCCU | acagguuucacgauuuucgaacau | acagguuucacgauuuucgaacauaagacaauuuuuauauguucguugucgucgaaaccugccu | scaffold_58:1786823..1786887:+ |
| Pdu-Mir-8_3p | UAAUACUGUCAGGUAAAGAUGUU | caucuuacugggcagcauuaga | caucuuacugggcagcauuagacuugguagccauacucuaauacugucagguaaagauguu | scaffold_13:14140837..14140898:- |
| Pdu-Mir-87_3p | GUGAGCAAAGUUUCAGGUGUGU | cugccugaaauuuucgcucagaccu | cugccugaaauuuucgcucagaccuguaaucaaaaggugagcaaaguuucaggugugu | scaffold_1:126238902..126238960:+ |
| Pdu-Mir-9_5p | UCUUUGGUUAUCUAGCUGUAUGA | auaaagcuagguuaccaaagcu | ucuuugguuaucuagcuguaugagugaagauauacuucauaaagcuagguuaccaaagcu | scaffold_1:38229931..38229991:+ |
| Pdu-Mir-92-1_3p | AAUUGCACUUGUCCCGGCCUGC | agguugugacagcugcaauaugg | agguugugacagcugcaauauggucuuuacaauccaauugcacuugucccggccugc | scaffold_1:194081866..194081923:- |
| Pdu-Mir-92-2_3p | AAUUGCACUUGUCCCGGCCUGC | cuggugggacuuaugcaacuug | cuggugggacuuaugcaacuugauuuucaaccacaaauugcacuugucccggccugc | scaffold_1:194083611..194083668:- |
| Pdu-Mir-92-3_3p | AAUUGCACUUGUCCCGGCCUGC | gggucgugauuugugcacuuuug | gggucgugauuugugcacuuuuguugucacucagucaaauugcacuugucccggccugc | scaffold_1:194084057..194084116:- |
| Pdu-Mir-96-P1_5p | UUUGGCACCAGUGGAAUAGUCAC | gaaauuccauuggugacauaug | uuuggcaccaguggaauagucacuuguuuuaaaaugugaaauuccauuggugacauaug | scaffold_1:138807632..138807691:- |
| Pdu-Mir-96-P2_5p | CUUGGCACUGGUAGAAUUCACUGA | agggauucuagagugcucaaaa | cuuggcacugguagaauucacugaguguauuugaccaucagggauucuagagugcucaaaa | scaffold_11:45340966..45341027:- |
| Pdu-Mir-96-P3_5p | AAUGGCACUGGUAGAAUUCACGG | gugggucuucuggugccgugca | aauggcacugguagaauucacgguucuuuaagaauuccgugggucuucuggugccgugca | scaffold_7:11760213..11760273:- |
| Pdu-Mir-96-P1bx_or_P1dx_5p | AUUUGGCACUAGUGGAAUGGUC | gaguuucauuaaugccaacacc | auuuggcacuaguggaauggucaucuguuauccgaugugaguuucauuaaugccaacacc | scaffold_39:3782468..3782528:- |
| Pdu-Mir-96-P1by_or_P1dy_5p | AUUUGGCACGAGUGGAAUAUUCG | guuauccaccugugccauauaa | auuuggcacgaguggaauauucguaucucucaccgguuauccaccugugccauauaa | scaffold_39:3780733..3780790:- |
| Pdu-Mir-1994_3p | missing but found in previous prediction/genome assembly | | | |
| Pdu-Mir-2001_5p-3p | missing but found in previous prediction/genome assembly | | | |
| Pdu-Mir-33_3p | missing but found in previous prediction/genome assembly | | | |
| Pdu-Mir-137 | Missing but putative gene locus found | | | |

### Bilateria

- **Let-7:**
  - **Bilaterian-specific**
  - **Paralogues/Orthologues:**
    - **Protostomes:** Most of them have **1 copy - Let-7_5p**
      - **Cte, Lgi, Cgi, Lan**
    - **Efe** has 5 copies:
      - **P10_5p**
      - **P11_5p**
      - **P12_5p**
      - **P13_5p**
      - **P14_5p**
    - Other paralogues appeared in some Protostomes and Vertebrates
  - **Seed:**
    - **5p_GAGGUAG** is almost the only one present
  - **Gene Clusters:**
    - Bilaterian-conserved **- Mir-10-P2_Let-7_Mir-10-P3**
  - **Pdu:**
    - **Pdu-Mir-Let-7_5p -** U**gagguag**UAGGUUGUAUAGUA
      - **3p* -** cuguacaaccuucuagcuuucc
      - The sequence shows complete conservation with several Let-7_5p pf pther organisms
      - 3p* also shows complete allignments with 3p* of several organisms

    **> Lgi-Let-7_3p***
    Length=22

Score = 41.7 bits (22),  Expect = 1e-06
Identities = 22/22 (100%), Gaps = 0/22 (0%)
Strand=Plus/Plus

```
Query  1   CTGTACAACCTTCTAGCTTTCC  22
           ||||||||||||||||||||||
Sbjct  1   CTGTACAACCTTCTAGCTTTCC  22
```

- **Gene Clusters:**
    - **Bilaterian-conserved Mir-10-P2_Let-7_Mir-10-P3** Cluster is present in pdu with the same orientation

- # Mir-1:
    - **Bilaterian-specific**
    - **Paralogues/Orthologues:**
        - **Protostomes:** Most of them have **1 copy - Mir-1_3p**
            - **Cte, Lgi, Cgi, Lan**
        - **Efe** has 4 copies:
            - **P5_3p**
            - **P6_3p**
            - **P7_3p**
            - **P8_3p**
        - Other paralogues appeared in **vertebrates**
    - **Seed:**
        - **3p_GGAUGU** is the only seed
            - Apparently **3p_mature arm** is the only one used
    - **Gene Clusters:**
        - Bilaterian-conserved **Mir-1_Mir-133**
            - **In Cte**: Mir-133_Mir-1 (within 6000ntds)
    - **Pdu:**

    Mir-1 gene might be trascribed in both directions (**Plus and Minus strands**). However, in the 2020SmallRNAseq the mature and star read counts of the minus strand are very low (19 and 1 respectively.
- **Pdu-Mir-1_3p (Plus-strand) -** U**ggaaugu**AAAGAAGUAUGUAG
    - **5p* -** acauacuucuucauauaccaua
    - Sequence shows complete conservation with several mir-10_3p of other organisms
    - Also 5p* shows almost complete conservation with several 5p*

**> Cli-Mir-1-P4_5p***
Length=23

Score = 36.2 bits (19),  Expect = 6e-05
Identities = 22/23 (96%), Gaps = 1/23 (4%)
Strand=Plus/Plus

```
Query  1   ACATACTTCTTCATATACC-ATA  22
           ||||||||||||||||||| |||
Sbjct  1   ACATACTTCTTCATATACCCATA  23
```

**> Ete-Mir-1-P2_5p***
Length=23

Score = 25.1 bits (13),  Expect = 0.13
Identities = 20/23 (87%), Gaps = 1/23 (4%)
Strand=Plus/Plus

```
Query  1   ACATACTTCTTCATATACC-ATA  22
           ||||||||||| || |||| |||
```

Sbjct  1   ACATACTTCTTTATGTACCCATA  23

- **Pdu-Mir-1_3p-5p (Minus-strand) -** U**ggaaugu**AAAGAAGUAUGUAG
  - **5p -** a**cauacuu**cuucauauaccaua

- **Both arms** might be **active.**
  - In the **2019SmallRNAseq-2022MirDeep2-2022Genomev2.1** the read counts for this gene are very low (~20 for 3p and 1 for 5p)

- **3p_Seed** sequence is **not conserved**
- 3p_mature shows quite good Plus-Minus alignments with several Mir-1_5p*
  - **Cli-Mir-1-P4_5p***

Length=23

Score = 36.2 bits (19),  Expect = 6e-05
Identities = 19/19 (100%), Gaps = 0/19 (0%)
Strand=Plus/Minus

Query  2   GGTATATGAAGAAGTATGT  20
|||||||||||||||||||
Sbjct  19  GGTATATGAAGAAGTATGT  1
**> Gga-Mir-1-P4_5p***
Length=23

Score = 30.7 bits (16),  Expect = 0.003
Identities = 16/16 (100%), Gaps = 0/16 (0%)
Strand=Plus/Minus

Query  5   ATATGAAGAAGTATGT  20
||||||||||||||||
Sbjct  16  ATATGAAGAAGTATGT  1

- Also 5p* shows good allignments with several **Mir-1-pre** and **Mir-1_3p**

**> Xtr-Mir-1-P2_pre**
Length=61

Score = 41.7 bits (22),  Expect = 1e-06
Identities = 22/22 (100%), Gaps = 0/22 (0%)
Strand=Plus/Minus

Query  1   ACATACTTCTTTACATTCCATA  22
||||||||||||||||||||||
Sbjct  59  ACATACTTCTTTACATTCCATA  38
**> Lgi-Mir-1_3p**
Length=22

Score = 38.1 bits (20),  Expect = 2e-05
Identities = 20/20 (100%), Gaps = 0/20 (0%)
Strand=Plus/Minus

Query  1   ACATACTTCTTTACATTCCA  20
||||||||||||||||||||
Sbjct  20  ACATACTTCTTTACATTCCA  1

- **Gene Clusters:**
  - Bilaterian-conserved **Mir-1_Mir-133**
    - **In Pdu:** Mir-1-3p(Plus and Minus strand)_Mir-133 (within 7.000 ntds)

- **Mir-10:**

- **Eumetazoa**
- **Paralogues:**
    - **P1_5p**,**P2_5p**,**P3_5p** are **Bilaterian-specific**
    - **P4_3p** is **Protostome-specific**
        - In Protostomes this copy shifted to the **3p_mature** arm
    - **P5_5p**, **P6_5p** are **Lophotrocozoa-specific**
    - **P7** is **Annelid** (Note:another Paralogue 7 is in Brachiostoma Floridae, but must be a different one). But also in Amphiouxus (most probably an indipendent paralogue)

- **Efe** as further duplications of almost all paralogues

- **Seeds:**
  **P1_5p** CCCUGUA/ACCCUGU
  **P2_5p -** ACCCGUA
  **P3_5p -** CCCUGAG
  **P4_3p -** AAGCUCG
  **P5_5p**, **P6_5p -** UACCCUG/UUACCCU
  **P7_5p -** Cte_ACCCUGU/Efe_ACACUGU
- **Gene Clusters:**
  **Bilaterian-conserved** gene cluster:
  **Mir-10-P2_Let-7_Mir-10-P3**

- **In Pdu:**
  **All Paralogues** appear to be present in Pdu, except for P6 which has not been found yet.
- **Pdu-Mir-10-P1_5p -** U**acccugu**AGAUCCGAAUUUGU
    - **3p*** - aaauucgagucugcggaguguug
    - 5p_mature sequence shows complete conservation with several other 5p_mature
    - 3p* shows mild conservation (~10 ntds) with some 3p* of other animals
- **Pdu-Mir-10-P2_5p -** A**acccgua**CAACCGAACUUGUG
    - **3p*** - caagcucgcucuuacgggccug
    - 5p_mature sequence shows almost complete conservation with several other P2-5p_mature. Complete only with Cte and Efe.
    - 3p* shows mild conservation with 3p* of other organisms
- **Pdu-Mir-10-P3_5p-3p -** U**cccugag**ACCCUAACUUGUGA
    - **3p* -** acagguugaggucuugggacua

- 3p_arm might also be active. **Both-arms might be** active maybe. In the 2019SmallRNAseq-MirDeep2-2019Genome prediction the reads count are in the same order of magnitude (~16.000vs10.000)
- 5p_mature sequence shows complete conservation with several others
- 3p* star also shows good allignments with some P3_pre and 3p* sequences

- **Pdu-Mir-10-P4-1_3p-5p_+_ -** A**gaagcuc**GGUUCUACAGGUA -Version1-2022Genome
    - **5p*** - acccuguagacccggguuugugu

*A**gaagcuc**GGUUCUACAGGUU -Version2-2019Genome*

- ***5p**** *- acccuguagacccgguuugugu*

- **Plus-Strand_Conserved - Not main strand used**
- 5p_arm might also be active. **Both-arms** might be **active** maybe.  In the 2019SmallRNAseq-MirDeep2-2019Genome prediction the reads count are in the same order of magnitude (~2.600vs1.200).

- **Pdu-Mir-10-P4-1_5p_-_ -** U**accugua**GAACCGAGCUUCUG -Version1-2022Genome
  - **3p*** - caaacccggucuacaggguau -Version1-2022Genome

- *A**accugua**GAACCGAGCUUCUG -Version2-2019Genome*
  *caaacccggucuacaggguau -Version2-2019Genome*

- **Minus-strand - Main strand used**
- According to 2019smallRNAseq this is the main mature sequence used from the Mir-10-P4 gene locus. From the 2019SmallRNAseq-2022MirDeep2-2022Genome prediction, more than 2x10^6 5p_mature reads are found.

- **Pdu-Mir-10-P4-1_5p Plus** and **Minus Strand in Platy:** Probably in Pdu both genomic strands are transcribed:
  - **Seq** A**gaagcuc**GGUUCUACAGGUU is the "original" conserved 3p_mature (Plus-Strand)
    - This strand has a conserved seed.
  - **Seq** A**accugua**GAACCGAGCUUCUG is 5p-mature (Minus-Strand)
    - This sequence does not have a conserved seed and its activity is probably Pdu-specific.
    - This is the Minus strand of the Pdu-Mir-10-P4_5p

```
> Cte-Mir-10-P4_3p
Length=22

 Score = 34.4 bits (18),  Expect = 3e-04
 Identities = 20/21 (95%), Gaps = 0/21 (0%)
 Strand=Plus/Minus

Query  1   TACCTGTAGAACCGAGCTTCT  21
           |||||||||||| ||||||||
Sbjct  21  TACCTGTAGAACAGAGCTTCT  1
```

- **Gene Structure:**

| | |
|---|---|
| 5p_Mature | |
| **5p-TACCTGTAGAACCGAGCTTCTG**TGATTTGCTCTGAACAACA**CAAACCCGGGTCTACAGGGTAT-3p** | P4_-_ |
| 3p-**ATGGACATCTTGGCTCGAAGA**CACTAAACGAGACTTGT**TGTGTTTGGGCCCAGATGTCCCA**TA-5p | _+_ |
| 3p-5p_Mature | |
| 5p-**ACCCTGTAGACCCGGGTTTGTGT**TGTTCAGAGCAAATCAC**AGAAGCTCGGTTCTACAGGTA**-3p | P4_+_ |

| | |
|---|---|
| 3p-**TGGGACATCTGGGCCCAAACACA**ACAAGTCTCGTTTAGTG**TCTTCGAGCCAAGATGTCCAT**-5p | _-_ |

- **Pdu-Mir-10-P4-2_5p -** A**acccugu**GGAUACGGGCUUGU
    - **3p*** - aagcucguuuuuacaagguguc

- **Arm-shift** from 3p to **5p_mature.**
- **5p_mature** shows **good partial allignment** with some Mir-10-P4_5p* of other species.
- -3p_star shows also some good partial allignments with some Mir-10-P4_3p_mature of other species.

```
> Tca-Mir-10-P4_5p*
Length=23

 Score = 25.1 bits (13),  Expect = 0.19
 Identities = 17/19 (89%), Gaps = 0/19 (0%)
 Strand=Plus/Plus

Query  2   ACCCTGTGGATACGGGCTT  20
           ||||||| ||| |||||||
Sbjct  2   ACCCTGTAGATCCGGGCTT  20

> Lpo-Mir-10-P4j_5p*
Length=23

 Score = 25.1 bits (13),  Expect = 0.19
 Identities = 17/19 (89%), Gaps = 0/19 (0%)
 Strand=Plus/Plus

Query  2   ACCCTGTGGATACGGGCTT  20
           ||||||| ||| |||||||
Sbjct  2   ACCCTGTAGATCCGGGCTT  20

> Lpo-Mir-10-P4n_3p
Length=23

 Score = 28.8 bits (15),  Expect = 0.015
 Identities = 18/19 (95%), Gaps = 1/19 (5%)
 Strand=Plus/Plus

Query  1   AAGCTCGTTTTTACAAGGT  19
           ||||||||||||||| |||
Sbjct  2   AAGCTCGTTTTTACA-GGT  19
```

- **Pdu-Mir-10-P5_5p -** U**uacccug**UAGAACCGAGCGAGUA
    - **3p* -** ccacucaguucacagggucauu
    - 5p-mature shows complete conservation with Cgi, Cte and Efe.
    - 3p* does not show any conservation with other 3p*

- **Pdu-Mir-10-P6_5p -** U**uuacccu**GUGAAUAGAGAGUG
    - **3p*** - gucucauuucuugggguaagca

    **5p_mature** shows few partial allignments with some other species. The sequence probably changed a lot in P. dumerilii. However,
    **5p_seed** is conserved.

The 3p* sequence shows no significant allignment.

```
> Npo-Mir-10-P6_5p*
Length=23

 Score = 25.1 bits (13),  Expect = 0.19
 Identities = 13/13 (100%), Gaps = 0/13 (0%)
 Strand=Plus/Plus

Query  2   TTACCCTGTGAAT  14
           |||||||||||||
Sbjct  2   TTACCCTGTGAAT  14

> Efe-Mir-10-P6_5p
Length=22

 Score = 21.4 bits (11),  Expect = 2.5
 Identities = 13/14 (93%), Gaps = 0/14 (0%)
 Strand=Plus/Plus

Query  3   TACCCTGTGAATAG  16
           |||||||||| |||
Sbjct  2   TACCCTGTGAGTAG  15

> Efe-Mir-10-P6_3p*
Length=22

 Score = 23.3 bits (12),  Expect = 0.70
 Identities = 16/18 (89%), Gaps = 0/18 (0%)
 Strand=Plus/Minus

Query  2   TTACCCTGTGAATAGAGA  19
           |||||| | |||||||||
Sbjct  20  TTACCCAGCGAATAGAGA  3

> Lgi-Mir-10-P6_5p
Length=23

 Score = 19.6 bits (10),  Expect = 9.0
 Identities = 12/13 (92%), Gaps = 0/13 (0%)
 Strand=Plus/Plus

Query  2   TTACCCTGTGAAT  14
           ||||||||| |||
Sbjct  2   TTACCCTGTTAAT  14
```

- **Pdu-Mir-10-P7_5p -** C**acccugu**AGAACCGAGCUUGU
- **3p* -** caggcugguuacaccgggucaa
- Seq **5p_**A**acccugu**GGAUACGGGCUUGU - **Pdu-Mir-P4_5p/Pdu-Mir-P7_5p/Pdu-Mir-P1x_5p:**
  - **3p* -** aagcucguuuuuacaagguguc
  - Possible homo-/para-logies are with:
    - **Mir-10-P4-5p***, P4 has 3p-arm mature in all other animals. Pdu has also an active P4_3p. However it might have a P4 copy, using the 5p.
      - In favour of this paralogy, is the fact that many other P4_5p* have a good allignment.
      - Also the 3p*-arm shows the 2 top allignments two P4_3p

**> Tca-Mir-10-P4_5p***
Length=23

Score = 25.1 bits (13), Expect = 0.13
Identities = 17/19 (89%), Gaps = 0/19 (0%)
Strand=Plus/Plus

```
Query  2   ACCCTGTGGATACGGGCTT  20
           ||||||| ||| |||||||
Sbjct  2   ACCCTGTAGATCCGGGCTT  20
```

**> Isc-Mir-10-P4_3p**
Length=23

Score = 23.3 bits (12), Expect = 0.46
Identities = 17/19 (89%), Gaps = 1/19 (5%)
Strand=Plus/Plus

```
Query  1   AAGCTCGTTTTTACAAGGT  19
           |||||||||| |||| |||
Sbjct  2   AAGCTCGTTTCTACA-GGT  19
```

- **Cte-Mir-10-P7_5p**, shares the same seed ACCCUGU and 5 mismatches in the rest of the seq;

**> Bfl-Mir-10-P7_5p**
Length=24

Score = 25.1 bits (13), Expect = 0.13
Identities = 19/22 (86%), Gaps = 0/22 (0%)
Strand=Plus/Plus

```
Query  1   AACCCTGTGGATACGGGCTTGT  22
           |||||||||||| ||  |||||
Sbjct  1   AACCCTGTGGATCCGATCTTGT  22
```

- **Cte-Mir-10-P1_5p**, has the same Seed and same mature arm with 5 mismatches with Cte_Mir-10-P1-5p. However, another Mir-10-P1 paralogue is already present in Pdu.
    - Is not in the top allignments

- **(!)** Seq C**uacgcau**CUUUGGGUCUGAAAUG:
    - Aligns with the central/final part of Mir-10-P2_3p* of several vertebrates. It looks like there are no significant Alignments with Mir-10-P2 of protostomes.
    - It might be due to chance or convergence.
    - **Arm-shift** from 5p to **3p-mature arm**
        - The 5p* shows no significant allignments with miRNAs of other species

**> Sko-Mir-10-P2_3p***
Length=22
Score = 23.3 bits (12), Expect = 0.59
Identities = 14/15 (93%), Gaps = 0/15 (0%)
Strand=Plus/Plus

```
Query  4   CGCATCTTTGGGTCT  18
           ||||||| |||||||
Sbjct  7   CGCATCTCTGGGTCT  21
```

- **Gene Clusters:**
    - Bilaterian-conserved gene cluster:
      **Pdu-Mir-10-P3_Pdu-Let-7_Pdu-Mir-10-P2** within 2000ntds
- All the other gene sit on the same scaffold, within 1,6x10^6 ntds

- # Mir-124:
  - **Bilaterian-specific**
  - **Paralogues:**
    - **Many Protostomes** and other organisms have **1 copy - Mir-124_3p**
      - **Few** of these organisms switched to the **5p_mature arm**
    - **Some Protostomes** have indipendent duplications/Paralogues:
      - **Efe** has **4 copies:**
        - P5_3p
        - P6_3p
        - P7_3p
        - P8_3p
        - P9_3p
      - **Lgi** has **2 copies:**
        - P10_3p
        - P11_3p
      - **Cgi** has **2 copies:**
        - P12_3p
        - P13_3p
    - **Vertebrates:** P1. P2, P3, P4 paralogues appeared in vetebrates
  - **Seeds:**
  **3p_mature -** AAGGCAC - is the **most common** with probably very few variants
  **5p_mature -** GUGUUCA
- **Gene Clusters:**
  - No known conserved gene cluster found
- **Pdu:**
  - **Pdu-Mir-124_3p-5p -** U**aaggcac**GCGGUGAAUGCCA
    - **5p* -** aguguucacuguguacgccuuggu

- 3p_ mature shows **complete conservation** across all taxa
- 5p* also shows good conservation with some lophotrochozoans 5p* - Lgi and Lan
- **Note: Both arms** might be **active**. However, in the 2019SmallRNAseq it looks like, in earlier stages (<=48hpf) the two sequences are almost equivalent in numbers. At 6dpf the 3p_mature is more abundant.

- **?Pdu-Mir-124-B_3p?** - U**aaggcac**GCGGUGAAUGCCA
  - **5p* -** guguucacugguguacgccuuggu (Only predicted)

- **3p_mature** sequence is identical to the other gene.
  - From the 2019SmallRNAseq-MirDeep2 is not clear wether this is a false positive or a real gene. The star read count is only predicted and the MirDeep2 score is low.

- **Gene Clusters:**
  - No gene clusters found

- # Mir-133 Family:
  - **Bilaterian-specific**
  - **Paralogues:**

    - **Protostomes:** most have **1 copy - Mir-133_3p**
    - **Cgi** has **2 copies:**
        - **v1_3p -** seed UUGGUCC
        - **v2_3p -** seed UGGUCCC
    - **Efe** appears to have both arms active: **Efe-Mir-133_3p/5p**
    - **Vertebrates:** Many copies appeared with v1(seed_UUGGUCC ) and v2(seed_UGGUCCC):
        - P1, P2, P3 and P4.

- **Seeds:**
    - 3p_UGGUCCC is the main seed. Also present in v2 of vertebrates and Cgi
    - 3p_UUGGUCC is less common seed (probaby seed-shift). Also present in v1 of vertebrates and Cgi.
    - Efe has **both arms** active with:
        - **Mir-133_3p** seed_UGGUCCC
        - **Mir-133_5p** seed_GCUGGCU

- **Gene Clusters:**
    - **Bilaterian-conserved:** Mir-1_Mir-133 cluster
        - **in Cte:** Mir-133_Mir-1 (within 6000 ntds)
- **Pdu:**
    - **Pdu-Mir-133_3p -** U**uggucccc**CUUCAACCAGCUGU
        - **5p*** - agcugguugaaauagggccaaau
        - 3p_mature shows complete conservation with several bilaterians
        - 5p* shows almost complete conservation with Lgi, Lan and Cte. Good with mir-133-pre of other protostomes. Few with 5p* of other organisms

- **Gene Clusters:**
    - **Bilaterian-conserved:** Mir-1_Mir-133 cluster
        - **in Pdu:** Mir-133_Mir-1(Plus and Minus strand) (within 17000 ntds)

- # Mir-153 Family:
    - **Bilaterian-specific**
    - **Paralogues:**
        - **Protostomes** have just **one copy**
        - All paralogues are **vertebrate-specific**
        - Mir-153 appears to have mostly a **3p-mature** sequence
            - Few exception switched to 5p-mature sequence
        - **Efe**: 2 indipendent paralogues in Efe
            - **P5_3p**
            - **P6_3p**
        - **Cte** has just one copy
    - **Seeds:**
        - 3p_seed **UGCAUAG**
            - Few exception switched to the 5p-seed CAUUUUU/UCAUUUU
    - **Gene Clusters:**
        - No known conserved miRNA clusters found
    - **Pdu:**
        - **Pdu-Mir-153_5p-3p** - C**gagcuuu**UGUGAUUUGCAAU

        - 3p*_a**ugcauag**ucacaaaagugauc

- **Arm-Switch** from **3p arm** to **5p-mature** or **both arms** are **active**. However, in the 2019smallRNAseq the two sequences are almost in the same of magnitude, the ratio Mature/Star reads is ~10^4/10^3. this trend is seen in all the stages.
- The 3p* sequence has total identity with other Mir-153_3p, including the seed UGCAUAG

```
> Lgi-Mir-153_3p
Length=22

 Score = 39.9 bits (21),  Expect = 5e-06
 Identities = 21/21 (100%), Gaps = 0/21 (0%)
 Strand=Plus/Plus

Query  2   TGCATAGTCACAAAAGTGATC  22
           |||||||||||||||||||||
Sbjct  2   TGCATAGTCACAAAAGTGATC  22
```

- **Gene Clusters:**
    - No gene clusters found

- # Mir-184:
    - **Bilaterian-specific**
    - **Paralogues:**
        - **Bilaterians:** Most of them present **1 copy: Mir-184_3p**
        - **Cte** has **2 copies:**
            - **P3_3p**
            - **P4_3p**
                - they appear to be independent duplications from oter organisms
        - **Efe** has **4 copies:**
            - **o1_3p**
            - **o2_3p**
            - **o3_3p**
            - **o4_3p**
        - **Lgi** has **2 copies:**
            - **P5_3p**
            - **P6_3p**
        - **Cgi** has **44 copies:**
            - Named from P7 to P15. Each of these has several duplications
                - Few exceptions use the **5p_mature** arm
    - **Seeds:**
        - 3p_GGACGGA appears to be the only one used.
        - Few exceptions are only seen in some copies of Cgi which use the 5p_mature (seed_CUUGUCA/UUGUCAC)
    - **Gene Clusters:**
        - Not known conserved clusters
        - **In cte:** Made by **mir-184-P3** and **P4** (within 200ntds)
    - **Pdu:**
        - **Pdu-Mir-184_3p -** U**ggacgga**GAACUGAUAAGGGC
            - **5p* -** ccuucucacuuguucguccggu

        - 3p_mature shows complete conservation between many Bilaterians
        - 5p* shows good conservation between Cte, Lgi and Cgi (~18/19). Less with other organisms

- # Mir-190:
    - **Bilaterian-specific**
    - **Paralogues:**
        - **Most Bilaterians** appear to have **1 copy: Mir-190_5p**
        - **Efe** has **4 copies:**
            - **P3_5p**
            - **P4_5p**
            - **P5_5p**
            - **P6_5p**
        - **Lan** has **2 copies:**
            - **P3_5p**
            - **P4_5p**
        - **Lgi** has **1 gene** which appears to be **processed in 2 ways**:
            - **Mir-190-v1_5p -** seed_GAUAUGU
            - **Mir-190-v2_5p -** seed_AUAUGUU
    - **Seeds:**
        - 5p_mature - GAUAUGU is the only seed with very few exceptions that use 3p_mature
    - **Gene Clusters:**
        - Not known conserved clusters
        - **In Cte: mir-190** is probably in the same chromosome as the Mir-184-P3/P4 cluster. However, the distance between these genes is over 80.000 ntds

- **Pdu:**
    - **Pdu-Mir-190_5p -** A**gauaugu**UUGAUAUAUUUGGUGG
        - **3p* -** accauguaucaaacaugucaug
        - 5p_mature shows complete conservation with Efe, Cte, Cgi, Lgi.
            - almost complete conservation with several other bilaterians
        - 3p* is only predicted from the genome ad no reads were found in the 2020SmallRNAseq. However, it shows almost complete conservation with Lgi-v2_3p* (18ntds). Some conservation ( around 13 ntds), but in Plus/minus oritentation, with 5p and precursors of several bilaterians.

- # Mir-193:
    - **Bilaterian-specific**
    - **Paralogues:**
        - **P1_3p and P2_3p:**
            - **Bilateria-specific**
            - Both present in **all Bilaterians**
            - **3p-arm** appears to be the original **mature** with main seed **AAUGCCC**

            - Several indipendent duplications in many specie adopted the **5p_mature arm**, thus switching to the seeds GGGGUUU/GGGUCUU/...
    - **Cte** has the 2 paralogues:
        - P1_3p -A**acuggcc**CGUCAAGUCCCUCC-
        - P2_3p -G**aaugccc**UUUCAAAUCCUGGG-
    - **Efe** has **8 copies of P2** and **2 copies of P1**.
        - All of them appear to have kept the **3p_mature arm**
        - P1-g/h
        - P2c-v1/v2
        - P2d-v1/v2
        - P2e-v1/v2
        - P2f-v1/v2
    - **Lgi** and **Cgi** have the two **P1_3p** and **P2 _3p**
    - **Lan** has a duplication of both **P1** and **P2:**
        - P1i_3p
        - P1j_3p
        - P2i_3p
        - P2j_3p
- **Seeds:**
    - **P1_3p and P2_3p:**
        - ACUGGCC/AAUGCCC/ACUGGCC are the most common
        - 5p_arm paralogues show GGGGUUU/GGGUCUU/GGGAUUU/...
- **In Pdu:**
    - **P1_3p -** <u>Not yet found</u>
    - **Pdu-Mir-193-P2_3p -** C**aaugccc**UAUGAAAUCCUAAA
        - **5p* -** ugggauuucugggggcauccugug
        - 3p_mature shows complete identity with P2_3p of few distant specie (Pfl, Sko). Almost complete identity with many other specie P2_3p ( ~ 16-20 ntds).
        - 5p* also shows good identity with Cgi and Lan (~20ntds). Quite good with many others P2_5p* ( ~ 14 ntds)

**> Pfl-Mir-193-P2_3p**
Length=22

Score = 39.9 bits (21),  Expect = 5e-06
Identities = 21/21 (100%), Gaps = 0/21 (0%)
Strand=Plus/Plus

```
Query  2   AATGCCCTATGAAATCCTAAA  22
           |||||||||||||||||||||
Sbjct  2   AATGCCCTATGAAATCCTAAA  22
```

**> Lan-Mir-193-P2j_3p**
Length=22

Score = 30.7 bits (16),  Expect = 0.003
Identities = 20/22 (91%), Gaps = 0/22 (0%)
Strand=Plus/Plus

```
Query  1   CAATGCCCTATGAAATCCTAAA  22
           |||||||||  |||||||||||
Sbjct  1   CAATGCCCTGCGAAATCCTAAA  22
```

- # Mir-153 Family:
  - **Bilaterian-specific**
  - **Paralogues:**
    - **Protostomes** have just **one copy**
    - All paralogues are **vertebrate-specific**
    - Mir-153 appears to have mostly a **3p-mature** sequence
      - Few exception switched to 5p-mature sequence
    - **Efe**: 2 indipendent paralogues in Efe
      - **P5_3p**
      - **P6_3p**
    - **Cte** has just one copy
  - **Seeds:**
    - 3p_seed **UGCAUAG**
      - Few exception switched to the 5p-seed CAUUUUU/UCAUUUU
  - **Gene Clusters:**
    - No miRNA clusters
  - **Pdu:**
    - **Pdu-Mir-153_5p**_C**gagcuuu**UGUGAUUUGCAAU
      - 3p*_a**ugcauag**ucacaaaagugauc
      - Switched from **3p arm** to **5p-mature**.
      - The 3p* sequence has total identity with other Mir-193_3p, including the seed UGCAUAG

    **> Lgi-Mir-153_3p**
    Length=22

    Score = 39.9 bits (21),  Expect = 5e-06
    Identities = 21/21 (100%), Gaps = 0/21 (0%)
    Strand=Plus/Plus

    ```
    Query  2   TGCATAGTCACAAAAGTGATC  22
               |||||||||||||||||||||
    Sbjct  2   TGCATAGTCACAAAAGTGATC  22
    ```

- In the 2020smallRNAseq the ratio Mature/Star reads is ~10^4/10^3
- In cluster with unknown, probably not significant sequence.

- # Mir-2001:
  - **Bilaterian-specific**
    - Present in **Protostomes** and **Ambulacrarians**. Apparently **lost in Chordates**
  - **Paralogues:**
    - **1 copy** present in most of the species:
    - **Lan** has **2 copies:**
      - **P1_5p**
      - **P2_5p (only predicted)**
    - **Efe** apparently lost the gene
  - **Seeds:**
    - **5p_seed** are mainly in seed-shift between **UUGUGAC/UGUGACC**
    - **Cte** and **Cgi** have UUGUGAC
    - **Lgi** and **Lan** have UGUGACC

- **Gene Clusters:**

- **Bilaterian-Conserved gene cluster: Mir-252-P1/P2_Mir-2001**
- **In Cte:** Mir-2001_~2000ntds_Mir-252-P2_~1000ntds_Mir-252-P1
  -

- **In Pdu:**
  - **1 copy** is present:
    - **Pdu-Mir-2001_5p** - U**ugugacc**GUUACAAUGGGCA
      - 3p* - cucauuguuacguugacaacu
    - In **Seed-shift respect to Cte** in 5'-direction.
    - **Note:** in the 2019SmallRNAseq the two arms reads are in the same order of magnitude (5p/3p ~ 9000/1000). So, also the 3p* sequence might have some activity. However, in all developmental stages, 5p_sequence is always between 5-10 times higher.
    - **5p_mature** has almost complete identity to other species.
    - **3p*_star** is not conserved

- **Gene Clusters:**
  - **Bilaterian-Conserved gene cluster: Mir-252-P1/P2_Mir-2001**
  - Mir-2001_~500ntds_Mir-252-P2_~1000ntds_Mir-252-P1

- # Mir-210 Family:
  - **Bilaterian-specific**
  - **Paralogues:**
    - **Most Bilaterian:** apart form few exceptions have **one gene - Mir-210**
    - Some Bilaterians show indipendent duplications.
    - **Cte** has **2 copies:**
      - **P5_5p/3p**
      - **P6_3p**
    - **Efe** has **1 copy**
  - **Seeds and arms:**
    - **5p_arm** is the most common.
    - Most common **5p_seed** is UGUGCGU. Some Bilaterians seed-shited to 3p'-direction in UUGUGCG
    - **Cte-3p_seed** is GGUCAUU
  - **Gene Clusters:**
    - **Cte** has an **indipendent Mir-210-P5/P6** gene cluster

- **In Pdu:**
  - 3 copies appear in Pdu. The relationship with the copies of Cte is unclear.

- **Pdu-Mir-210-1_3p-5p -** C**uugugcg**UGUGACAGUGACAAU
  - 5p* - ccgucacuggcuccgcgcaaaga

- **Both arms** might **be active** - In the 2019SmallRNAseq-Mirdeep2-2019Genome prediction In all developmental stages the difference between 3p and 5p is less then an order of magnitude. Ranging from 2 to 5 fold higher.
- 3p_mature shows almost complete identity with Cte-Mir-210_P5 and Lan-Mir-210. The identity goes down to (16/16ntds)(1-16ntd) with several other species.
- 3p_seed shows a **seed shift** in 3'-direction **as Cte-Mir-210-P5** and some other bilaterians.

> **Cte-Mir-210-P5_3p**
Length=24
Score = 41.7 bits (22), Expect = 1e-06
Identities = 22/22 (100%), Gaps = 0/22 (0%)
Strand=Plus/Plus

```
Query  2   TTGTGCGTGTGACAGTGACAAT  23
           ||||||||||||||||||||||
Sbjct  2   TTGTGCGTGTGACAGTGACAAT  23
```

- **Pdu-Mir-210-2_3p -** U**ugugcgu**GGGACAGCGGCCCGC
  - 5p* - caccguugcuucaugcccagcg
  - 3p_mature shows partial allignment with Cte-Mir-210-P5 (14/15)(1-15/2-16). Partial allignments also with several other bilaterians (17-18 ntds)(2-19)

> **Lgi-Mir-210_3p**
Length=22
Score = 32.5 bits (17), Expect = 9e-04
Identities = 17/17 (100%), Gaps = 0/17 (0%)
Strand=Plus/Plus

```
Query  1   TTGTGCGTGGGACAGCG  17
           |||||||||||||||||
Sbjct  1   TTGTGCGTGGGACAGCG  17
```

> **Cte-Mir-210-P5_3p**
Length=24
Score = 23.3 bits (12), Expect = 0.52
Identities = 14/15 (93%), Gaps = 0/15 (0%)
Strand=Plus/Plus

```
Query  1   TTGTGCGTGGGACAG  15
           ||||||||| |||||
Sbjct  2   TTGTGCGTGTGACAG  16
```

- **Pdu-Mir-210-3_3p** - U**ugugcgu**GGAACAGCGACCAU
  5p* - gucacuggccuuggcacaagca
- 3p_mature shows almost 5'-side allignment with molluscs (Cgi-16/16 -- Lgi 17/18). Partial allignments at 5'-side with several other bilaterians (~15ntds with 1-2 Gaps after seed sequence)

> **Cgi-Mir-210_3p**
Length=21

Score = 30.7 bits (16), Expect = 0.003
Identities = 16/16 (100%), Gaps = 0/16 (0%)
Strand=Plus/Plus

```
Query  1   TTGTGCGTGGAACAGC  16
           ||||||||||||||||
Sbjct  1   TTGTGCGTGGAACAGC  16
```

> **Cte-Mir-210-P6_3p**
Length=22

Score = 25.1 bits (13), Expect = 0.13
Identities = 17/19 (89%), Gaps = 0/19 (0%)
Strand=Plus/Plus

```
Query  2   TGTGCGTGGAACAGCGACC  20
```

```
              ||||||| ||||||||||
Sbjct  2   TGTGCGTAAAACAGCGACC  20
```

- **Gene Clusters:**
  - All 3 copies appear in the same scaffold:
    - Two genes from a cluster within ~5800 ntds:
      - Mir-210-2_Mir-210-3
      - Mir-210-1 is more then 100.000 ntds far from this cluster.

- # Mir-216 Family:
  - **Bilaterian-specific**
  - **Paralogues:**
    - **P1_5p and P2_5p** are **Bilaterian-specific**.
    - **Cte** has a P1 duplication.
      - **P1c_5p**
      - **P1d_3p**
        - P1d shifted from 5p to **3p-mature** with seed AAGUUAC
    - **Efe** has a P2 duplication:
      - **P2a_5p**
      - **P2b_5p**
  - **Seeds**
    - **Mir-193-P1/P2_5p** has AAUCUCA/AAUAUCA as main seeds
  - **Gene Clusters:**
    - **Mir-12_Mir-216 Protostome-conserved** Cluster in Cte
      - **Cte:** is organized this way **Mir-12_Mir-193-P1d_Mir-193-P2_Mir-193-P1c**

- **Pdu:**
  - Has the 2 conserved bilaterian-paralogues, **P1 and P2**.
    - **Pdu-Mir-216-P1_5p -** U**aauauca**GCUGGUAAAAGUGAG
    - **Pdu-Mir-216-P2_5p -** A**aauauca**GCUGGUAAUUCUGAG

- **Gene Cluster:**
  Conserved **Mir-12_Mir-216 Protostome-cluster**:
- **Mir-12_Mir-216-P1_Mir-216-P2** within ~ 2300 ntds

- # Mir-219 Family:
  - **Bilateria-specific**
  - **Orthologues/Paralogues:**
    - **Protostomes**: most appear to have just one copy - **Mir-219_5p**
    - **Efe** posses **3 copies:**
      - **P3_5p**
      - **P4_5p**
      - **P5_5p**
    - **Lan** has both strands active
    - **Vertebrates: P1_5p** and **P2_5p** copies appeared in Vertebrates
  - **Seeds:**
    - **5p_**seed**_GAUUGUC** is the most common 5p_seed in **protostomes**

        - Other organisms appear to have minor variations respect to the 5p_seed of protostomes
        - no evident arm-switches in any organism
    - **Gene Clusters:**
        - Not known
    - **Pdu:**
        - **2** copies appear to be present in Pdu. However, they have identical 5p-3p-Hairpin sequence, so they might represent an artifact duplication of the genome. They might sit on repetitive element regions.
        - The two copies have been called **1** and **2**
        - **Pdu-Mir-219-1_5p -** U**gauuguc**CAAACGCAUUUCUUG - scaffold_6:22743322..22743380:+
            - **3p* -** agaacuguuuuggacaucagu

▪ 5p shows almost complete allignment (only 1ntd change respect to all other sequences) with many mir-219 from other species.
▪ 3p* also shows great allignments with some 3p of other organisms

```
> Lan-Mir-219_3p
Length=22

 Score = 28.8 bits (15),  Expect = 0.010
 Identities = 20/22 (91%), Gaps = 1/22 (5%)
 Strand=Plus/Plus

Query  1   AGAACTGTTT-TGGACATCAGT  21
           |||||||| | |||||||||||
Sbjct  1   AGAACTGTGTATGGACATCAGT  22
```

- **Pdu-Mir-219-2_3p -** U**gauuguc**CAAACGCAUUUCUUG - scaffold_8:20782311..20782369:-
    - **5p* -** agaacuguuuuggacaucagu

- **No Gene Cluster detected**

- ## Mir-22 Family:
    - **Bilaterian-specific** family
    - **Paralogues:**
        - 2 copies appeared in **Bilaterians**
            - **P1_3p**
            - **P2_3p**
        - **Efe** has a duplication of P1 and **lost P2**:
            - **P1c_3p**
            - **P1d_3p**
        - **Cte** has both Bilaterians copies
    - **Seeds:**
        - **3p_seed** is AGCUGCC for both copies
        - Looks like **all bilaterians** use the **3p_arm**
        - Many **Lophotrocozoans** on the database (incl. Lan, Lgi, Cgi, Efe, Cte, Efe) have a **seed-shift in 5'-**direction of **P1 paralogue**, thus changing to **GCUGCCU**
    - **Gene Clusters:**

        - **Conserved Bilaterian** Mir-22 copies cluster Mir-22-P1_Mir-22-P2
            - **in Cte:** Mir-22-P1_Mir-22-P2 within 500 ntds

- **In Pdu:**
    - Both Bilaterian conserved copies are present:
        - **Pdu-Mir-22-P1_3p** - A**gcugccu**GGUGAAGAGCUGUC
            - **5p*** - cggccuuucuccaggcaggacuuu
            - **Seed shift** in 5'-direction like other Lophothrocozoans
        - **Pdu-Mir-22-P2_3p-5p -** G**agcugcc**AAGUGAAGGGCUGU
            - **5p*** - aguuccuucuccuggcugcuguccc

- **Both arms** might **be active.** In the 2019SmallRNAseq-2020MirDeep2-2019Genome prediction the 5p arm shows higher reads in early stages (24-36hpf), from 2 to 5 fold. Almost equivalent reads in 48hpf. Then Higher reads of the 3p in later stages (72hpf-6dpf). However, the difference in later stages is is about 2-3 fold, never higher than an order of magnitude

- **Gene Clusters:**
    - Bilaterian conserved Mir-22 Cluster: Mir-22-P1_Mir-22-P2 within 1000ntds

- # Mir-242 Family:
    - **Bilaterian-specific** family
        - However apparently lost in many lineages as Chordates and Ecdysozoans. Present in Ambulacrarians and Lophothrocozoans.
    - **Paralogues:**
        - One copy in Bilaterians
        - **Lgi** has 2 copies:
            - P1_5p
            - P2_5p
        - **Cte** has one copy.
        - **Efe** apparently **lost** the gene
    - **Seeds:**
        - **5p_seed** is **UGCGUAG** is the most common. 5p_arm_mature is the only one used.
        - Some Ambulacraria seed shifted in **5' direction** to GCGUAGG
    - **Gene Clusters:**
        - No Known conserved **gene clusters**

- **In Pdu:**
    - According 2019SmallRNAseq-2022MirDeep2-2022Genome there might be 4 copies of the genes in a cluster in scaffold_7.
        - First three copies separeted by 300 ntds between each of them. The last copy ~9000 ntds further.
        - All copies present the same 5p_mature sequence and almost identical 3p*_star sequences (with max 2 ntd changes between them). Also the middle region of the precursor presents few variations. It might be an artefact of the prediction. However it cannot be excluded that this are 4 indipendent copies.
        - In general all four sequences have very low Star read counts. 5-6 orders of magnitude less than the mature read count.
        - Nomenclature:

    - Numbers 1,2,3,4 identify the 4 sequences
    - Letters a,b,c indentify the 3 different 3p* sequences. mir-242-1 and 4 have the same 3p* sequence

- **Pdu-Mir-242-1a_5p** - U**ugcguag**GCGUUGUGCACAGA
    - 3p* - uguguauuuugccuacgcaaauu

- **Pdu-Mir-242-2b_5p** - U**ugcguag**GCGUUGUGCACAGA
    - 3p* - uguguauuuuaccuacgcaaau

- **Pdu-Mir-242-3c_5p** - U**ugcguag**GCGUUGUGCACAGA
    - 3p* - uguguauuuugccuacgcaauu

- **Pdu-Mir-242-4a_5p** - U**ugcguag**GCGUUGUGCACAGA
    - 3p* - uguguauuuugccuacgcaaauu

- **Gene Clusters:**
    - See above: might be a Pdu-specific cluster..
    - Pdu-Mir-242-1a_~300ntds_Pdu-Mir-242-2b_~300ntds_Pdu-Mir-242-3c_~9000_Pdu-Mir-242-4a

- ## Mir-252 Family:
    - **Bilaterian-specific** family
    - **Paralogues:**
        - **2 copies** appeared in **Bilaterians:**
            - **P1_5p**
            - **P2_5p**
        - **Lan** has a duplication of both copies:
            - P1a_5p
            - P1b_5p
            - P2a_5p
            - P1b_5p
        - **Chordates**, appear to have lost Mir-252 family
    - **Seeds:**
        - **5p_seed** UAAGUAC is the most common. Ecdysozoans show a seed shift in 5'-direction.
        - **All bilaterians** use the **5p_arm_mature**
    - **Gene Clusters:**
        - **Bilaterian-conserved** cluster: Mir-252-Paralogues_Mir-2001
        - **In Cte:** Mir-2001_Mir-252-P2_Mir-252-P1 within ~3000ntds

- **In Pdu:**
    - Both bilaterians paralogues are present:
        - **Pdu-Mir-252-P1_5p -** C**uaaguac**UAGCGCCGCAGGA
            - **3p*** - cugcuguccuagugcuuaaug
        - **Pdu-Mir-252-P2_5p** - C**uaaguag**UAGCGCCGCAGGUA
            - **3p*** - ccugcacccugcugcuuauca

- **Gene Clusters:**
    - Has the entire **Bilaterian-conserved cluster:**

**Mir-2001_Mir-252-P2_Mir-252-P1** within ~1500ntds

- Note: in the Genme assembly v.2.1 (2022) Mir-2001 is no longer present.
- ## Mir-281 Family:
  - **Bilaterian-specific** family
  - **Paralogues:**
    - Appear to be indipendent in each clade:
      - **Cte** has 1 gene **3p_mature**
      - **Efe** has 3: **P5_3p/P6_3p/P7_5p**
      - **Lgi** has 2: **P8_3p/P9_3p**
      - **Cgi** has 1: **3p_mature**
      - **Lan** has 2: **P9_5p/P10_5p**
    - Mainly **insects** appear to have switched to the **5p_arm**
    - It looks like there have been main changes in different clades in the use of **3p** or **5p arm**
  - **Seeds**:
    - **3p_seed** is mainly **GUCAUGG**
    - **5p_seed** is mainly **AGAGAGC**
  - **Pdu:**

  So far, it appears to have just one.
- **Pdu-Mir-281_3p-5p -** U**gucaugg**AGUUGCUCUCUUUA
  - **5p -** a**agggagc**auucuuggacagu - **-** The 5p-seed has changed

- **Both arms** are **active.** In the 2019SmallRNAseq-2022Mirdeep2-2022Genome the two arms reads are in the same order of magnitude.
- The 5p_seed has a mutation A->G respect to the conserved seed, AG**G**GAGC. The same Seed is also present in the P7 paralogue of Efe.

- ## Mir-29 Family:
  - **Bilaterian-specific** family
  - **Paralogues:**
    - **P1_3p** and **P2_3p** are **Bilaterian-specific**
      - **Protostomes:** duplications of these paralogues appaeared in Pancrustacea and other species/clades
      - The most common active sequence is the **3p_mature.** Few examples of switches to the 5p_mature
    - **Cte** has the two P1_3p and P2_3p
      - They form the **Mir-29-P1/P2 Cluster**
    - **Efe** has duplicated both paralogues
      - P1e_3p
      - P1f_3p
      - P2c_3p
      - P2d_3p
    - **Cgi** has the two P1_3p and P2_3p
    - **Lgi** has P1_3p but lost P2_3p
    - **Lan** duplicated both paralogues:
      - **P1g_3p**
      - **P1h_3p**
      - **P2g_3p**
      - **P2h_3p**
  - **Seeds:**
    - **3p_seed** is mainly **AGCACCA**

- **Gene Clusters:**
  - **Cte:** has the **Mir-29-P1/P2 Cluster**

- **Pdu:**
  Pdu has the 2 bilaterian-specific conserved parlogues in cluster:

- **Pdu-mir-29-P1_3p -** U**agcacca**UUUGAAAUCAGUUU
  - 5p* - gcuggguucuucuggugcugga

- **3p-mature** shows **complete allignments** with mir-29-P1 of many other organisms
- **5p*-star** shows only partial allignments with other mir-29-P1.

- **Pdu-Mir-29-P2_5p-3p** - C**uggucuc**AAGUGGUGGAUAGA
  - **3p-** u**agcacca**uuugaaaucagu
  -
  - **Arm-shift** from 3p to 5p_mature or **Both arms** might be **active.**
    In the 2019SmallRNAseq-2020Mirdeep2-2019Genome the 5p_mature and 3p* counts are in the same order of magnitude. However, the abbundancy in all samples at all stages is always higher for the 5p_arm.

- **5p-mature** shows **almost complete identity** with some other **5p*_star**
- **3p*-star** shows **complete identity** with a lot other **3p_mature** of other species.

- Pdu might have numerous duplications in the same contig. However it might be an artefact of the prediction and/or of the genome assembly. This still need to be further investigated.
  - Anyway for these duplications there is only an identity with mir-29 only at the level of the seed sequences. The rest of the sequence does not give significant allignments with other mir-29

- **Gene Clusters:**
  - **Bilaterian-conserved** Mir-29 Paralogues cluster
    - **In Cte:**
      - **Mir-29-P1_~800ntds_Mir-29-P2**
    - **In Pdu:**
      - **Mir-29-P1_~11000ntds_Mir-29-P2**

- # Mir-31 Family:
  - **Bilaterian-Specific**
  - **Orthologues/Paralogues:**
    - Very conserved miRNA. Almost identical sequence between all bilaterians.
    - Most Bilaterians appear to have only one copy:
      - **Cte** has 1
      - **Efe** has 1
    - **Drosophilas** have 2 copies:
      - P1_5p
      - P2_5p
    - Some species have **apparently indipendent paralogues**:

        - **Echinoderm P. miniata** has 4:
            - P3_5p
            - P4_5p
            - P5a_5p
            - P5b_5p
        - **Lan** has 2:
            - P6_5p
            - P7_5p
        - **Chelicerata Crab** (L. Polyphemus) has 4
            - P13_5p
            - P14_5p
            - P15_5p
            - P16_5p
        - **Planaria Sme** has 2:
            - P19_5p
            - P20_5p
    - **Seeds/Arms:**
        - The only seed used is **5p_GGCAAGA**
    - **Clusters:**
        - No known conserved gene clusters

- **Pdu:**
    - **Pdu-Mir-31_5p -** AGGCAAGAUGUUGGCAUAGCUGA
        - 3p* - agcucugucccauguugccacc

- Prediction_2.0-2022-Genome3.0
- Complete conservation with many other species

```
> Cte-Mir-31_5p
Length=23

 Score = 43.6 bits (23),  Expect = 6e-07
 Identities = 23/23 (100%), Gaps = 0/23 (0%)
 Strand=Plus/Plus

Query  1   AGGCAAGATGTTGGCATAGCTGA  23
           |||||||||||||||||||||||
Sbjct  1   AGGCAAGATGTTGGCATAGCTGA  23

> Cgi-Mir-31_5p
Length=23

 Score = 43.6 bits (23),  Expect = 6e-07
 Identities = 23/23 (100%), Gaps = 0/23 (0%)
 Strand=Plus/Plus

Query  1   AGGCAAGATGTTGGCATAGCTGA  23
           |||||||||||||||||||||||
Sbjct  1   AGGCAAGATGTTGGCATAGCTGA  23
```

- **Gene Clusters:**
    - No gene clusters found

- # Mir-315 Family:
  - **Bilaterian-specific**
  - **Orthologues/Paralogues:**
    - All Bilaterians appear to have only one copy
    - **Cgi** has 2 copies:
      - Mir-315-v1
      - Mir-315-v2
  - **Seeds:**
    - 5p_seed UUUGAUU is the only seed present.
  - **Gene Clusters:**
    - No known gene clusters

- **In Pdu:**
  - **Pdu-Mir-315_5p -** U**uuugauu**GUUGCUCAGAAAGCC
    - **3p* -** cuaucggguaguaaucaaaaaa

- # Mir-33 Family:
  - **Bilaterian-specific**
  - **Orthologues/Paralogues:**
    - **Protostomes -** most, appear to have just **one copy: Mir-33_5p**
    - **Cte** and **Efe** have **1 copy**
    - **Lan** has 2 copies:
      - **P7_5p**
      - **P8_5p**
    - In Vertebrates appeared **P1 and P2 paralogues**.
  - **Seeds:**
    - **5p_ seed -** UGCAUUG - is the most common
    - **5p-**mature appears to be the most commonly used sequence
  - **Gene Clusters:**
    - Not known

- **Pdu:**
  Appear to have **1 copy**
- **Pdu-Mir-33_3p -** C**aaugcuu**CUGCAGUGCAAUCA
  - **5p* -** g**ugcauug**uaguugcauugca
  - **Arm-shift** from 5p to **3p-mature**
  - Also 5p* arm shows great conservation with 5p of other organisms

```
> Cte-Mir-33_5p
Length=21

 Score = 39.9 bits (21),  Expect = 5e-06
 Identities = 21/21 (100%), Gaps = 0/21 (0%)
 Strand=Plus/Plus

Query  1   GTGCATTGTAGTTGCATTGCA  21
           |||||||||||||||||||||
Sbjct  1   GTGCATTGTAGTTGCATTGCA  21
```

- **Gene Clusters:** not found

- # Mir-34 Family:
  - **Bilaterian-specific**

- **Orthologues/Paralogues:**
    - Most paralogues appeared in Vertebrate
    - Most Protostomes appear to have only one copy
        - **Lan** has **2 copies**
- **Seeds:**
    - **5p_seed** GGCAGUG is the main seed

- **Gene clusters:**
    - **Protostome conserved** cluster Mir-34_Mir-277-Mir-317:
        - **In Cte:** Mir-34_~400ntds_Mi-277-P2_~100ntds_Mi-277-P1_~150ntds_Mir-317

- **In Pdu:**
    - **Pdu-Mir-34_5p -** U**ggcagug**UGGUUAGCUGGUUGU
        - **5p*** - aaccacuaucugcccugucuua
    - **Gene Clusters:**
        - **In Pdu:**
            - Protostome conserved cluster Mir-34_Mir-277-Mir-317
            - Mir-317_~3500ntds_Mir-277_~26.000ntds_Mir-34

-
- # Mir-375 Family:
    - **Bilaterian-specific**
    - **Orthologues and paralogues:**
        - Most species appear to have only one copy
        - Cgi has **2 copies**
            - P3_3p
            - P4_3p
    - **Seeds:**
        - **3p_seed** UUGUUCG is the only seed, except for Cgi.

- **Gene Clusters:**
    - Not known
- **In Pdu:**
    - **Pdu-Mir-375_5p -** U**uuguucg**UCCGGCUCGCGUUA
        - 5p* - augugagcuauucguacagagc

- **Gene Clusters:**
    - Not found

- # Mir-7 Family:
    - **Bilaterian-specific**
    - **Orthologues and paralogues:**
        - Most Protostomes in the database show only one copy
        - Most duplications appeared in vertebrates
        - **Efe** has **3 copies**:
            - P7_5p
            - P8_5p
            - P9_5p

- **Seeds:**
    - **5p-seed** GGAAGAC is the only seed

- **Gene Clusters:**
    - No Known conserved gene clusters

- **In Pdu:**
    - **Pdu-Mir-7_5p** - U**ggaagac**UAGUGAUUUGUUGUU
        - **3p*** - caaugaaacacuaucuuccaua

- 5p_mature sequence shows complete allignment with many other organisms

- **Gene clusters:**
    - No known gene clusters

- ## Mir-71 Family:
    - **Bilaterian-specific**
    - **Orthologues and paralogues:**
        - Most of the species present ony one copy
        - On the side of **Deuterostomes is present only in Ambulacraria**
        - **Efe** has **3 copies:**
            - **P1_5p**
            - **P2_5p**
            - **P3_5p**
        - **Lan** has 2 copies:
            - **P4_5p**
            - **P5_5p**

- **Seeds:**
    - Main **5p_seed** is **GAAAGAC** with few exceptions

- **Gene clusters:**
    - Protostomes conserved cluster **Mir-2_Mir-71:**
      **In Cte:**
- **Cluster Mir-2 + Mir-71:**
    - Contains 4 Mir-2 copies + Mir-71 within <600ntds
        - Mir-71_Mir-2-o37_Mir-2-o42_Mir-2-o38_Mir-2-o36

- **In Pdu:**
    - **Pdu-Mir-71_5p -** U**gaaagac**AUGGGUAGUGAGAUG
        - **3p* -** gcucgcuacccugucuuuccag

- **Version_1:** from 2019SmallRNAseq-2022Mirdeep2-2022Genome
- **5p_mature** shows **complete allignment** with many other organisms
- **3p_star** shows partial allignment (~14-16ntds) with many other organisms

- **Note:** in the 2019SmallRNAseq-2022Mirdeep2-2022Genome the sequence shows very high read counts (>10^6) with star reads 2 orders of magnitude lower

- ***Version2:*** from 2019SmallRNAseq-2020Mirdeep2-2019Genome
  ***Pdu-Mir-71_5p -*** *U**gaaagac**AUGGGUAGUGGAUG*

- ***3p* -** cucgcuacccuguuuccagg*

- *5p_mature sequence shows almost complete allignment with many other organisms*
- ***Note:** in the 2019SmallRNAseq prediction the sequence shows very low mature read counts (>30) and no star read count. Also th MirDeep2 score is low*

- **Gene Clusters:**
    - Protostomes conserved cluster Mir-2_Mir-71:
        - Is composed of 5 or 6 Mir-2 family genes + Mir-71:
            - Pdu-Mir-2-oA1_Pdu-Mir-2-oB1_Pdu-Mir-2-oC1_Pdu-Mir-2-oD1_Pdu-Mir-2-oE1_?Pdu-Mir-2-oF1?_Pdu-Mir-71
                - All within <4000ntds

- # Mir-76 Family:
    - **Bilaterian-specific**
    - **Orthologues and paralogues:**
        - Most of the species contain only one copy
        - On the side of **Deuterostomes is present only in Ambulacraria.**
        - **Lan** has **2 copies:**
            - **P1_3p**
            - **P2_3p**
    - **Seeds:**
        - **3p_seed UCGUUGU** appears to be the only one present.
    - **Gene Clusters:**
        - No known conserved gene clusters
    - **In Pdu:**
        - **Pdu-Mir-76_3p-5p -** U**ucguugu**CGUCGAAACCUGCCU
            - 5p - acagguuucacgauuuucgaacau

- **3p_mature** sequence shows **complete allignment** with other organisms.
- **Both arms** might be **active** maybe
    - In the 2019SmallRNAseq the the 5p and 3p reads are in the same order of magnitude. The 3p appears to be always higher in all developmental stages. But only 2-3folds higher.
    - Both arms could be active

- **Gene Clusters:**
    - No gene Clusters found

- # Mir-8 Family:
    - **Bilaterian-specific**
    - **Orthologues and paralogues:**
        - Most Protostomes in database appear to have **one copy**
        - Most Paralogues appeared in Vertebrates
        - **Efe** has **3 copies:**
            - **P4_3p**
            - **P5_3p**

        - **P6_3p**
  - **Seeds:**
    - **3p_seed AAUACUG** is the main seed for most organisms
  - **Gene clusters:**
    - No Known conserved gene clusters

- **In Pdu:**
  - **Pdu-Mir-8_3p -** U**aauacug**UCAGGUAAAGAUGUU
    - 5p* - caucuuacugggcagcauuaga

- 3p_mature shows almost **complete allignments** with many other organisms

- **Gene clusters:**
  - No gene clusters found

- ## Mir-9 Family:
  - **Bilaterian-specific**
  - **Orthologues and paralogues:**
    - Many Protostomes show just one copy
    - **Cte** has **1 copy**
    - **Lgi** has **1 copy**
    - **Lan** has **1 copy**
    - **Cgi** has **3 copies:**
      - **P12_5p**
      - **P13_5p**
      - **P14_5p**
    - **Efe** has **2 copies:**
      - **P15_5p**
      - **P16_5p**
  - **Seeds:**
    - **5p_seed CUUUGGU** is the main seed. Shared in almost all protostomes.

- **Gene Custers**
  - No known conserved gene clusters

- **In Pdu:**
  - Pdu-Mir-9_5p - U**cuuuggu**UAUCUAGCUGUAUGA
    - 3p* - auaaagcuagguuaccaaagcu

- **5p_mature** shows **complete allignments** with many other organisms

- **Gene Clusters:**
  - No gene cluster found

- ## Mir-92 Family:
  - **Bilaterian-conserved**
  - **Paralogues:**
    - Almost all organisms show duplications of the gene.

            - However, It is not clear either from phylogenetic or syntenic information how many Mir-92 genes were present in the last common ancestor of bilaterians and how the vertebrate Mir-92s relate to the invertebrate Mir-92s and thus these multiple paralogues in invertebrates are classified here as orphans pending additional data.
        - **Cgi** has **4 copies**
        - **Lgi** has **5 copies**
        - **Lan** has **5 copies**
        - **Cte** has **3 copies**
        - **Efe** has **2 copies**
    - **Seeds:**
        - **3p_seed AUUGCAC** is the main seed, with few exceptions:

- **Gene clusters:**
    - No known conserved gene clusters.
    - In Cte the 3 copies are in cluster:
      Mir-92-o37_Mir-92-o35_Mir-92-o36 (within 1000ntds)

- **In Pdu:**
    - 3 copies are present. They all show the same mature sequence but slightly different star sequences. They are all 3 in a cluster, like the 3 copies of Capitella.

- **Pdu-Mir-92-1_3p** - A**auugcac**UUGUCCCGGCCUGC
    - **5p*** - agguugugacagcugcaauaugg
- **Pdu-Mir-92-2_3p -** A**auugcac**UUGUCCCGGCCUGC
    - **5p*** - cuggugggacuuaugcaacuug
- **Pdu-Mir-92-3_3p** - A**auugcac**UUGUCCCGGCCUGC
    - **5p* -** gggucgugauuugugcacuuuug

- 3p_mature sequence shows complete allignments with othe organisms.

- **Note:**
    - The sequences have been named oA, oB, oC according to order in which they appear in the gene cluster. The 'o' stands for 'orphan' as annotated in the MirGeneDB database since phylogenetic informations are not complete.

- **Gene Clusters:**
    - No known conserved gene clusters (lack of Phylogenetic information). However, Pdu shows a cluster of 3 Mir-92 copies, as in Cte
        - **Pdu-Mir-92-1_Pdu-Mir-92-2_Pdu-Mir-92-3** all within (<2100ntds)
        - Mir-92-o37_Mir-92-o35_Mir-92-o36 (within 1000ntds)

- ## **Mir-96 Family**:
    - **Blaterian-conserved**
    - **Paralogues:**
        - **P1_5p**, **P2_5p, P3_5p** are **bilaterian-specific**
            - have mostly the seed UUGGCAC
            - **P3_5p** shifted to UGGCACU and some to AUGGCAC (as Platy)

- **Annelids** (Cte and Efe) appear to have **lost P3**.
    - **Cte** posses only **P1_5p** and **P2_5p.**
    - **Efe** posses **P1_5p** and a duplicated **P2_5p** in:
        - **P2d_5p**
        - **P2e_5p**
    - All paralogue have the **same seed**
- **Molluscs** (Cgi and Lgi) have all 3 paralogues and a duplicaton of the P1 paralogue:
    - **P1a_5p -** seed UUGGCAC
    - **P1b_5p** - seed UUUGGCA (seed shift in 3p-direction)
- **Cgi** presents also a duplication of the P2 paralogue:
    - **P2h_5p**
    - **P2i_5p**
- **Lan** possess all the **3 paralogues**

- **Seeds:**
    - **P1_5p, P2_5p,** have mostly the seed **UUGGCAC**
    - **P3_5p** mutated to **AUGGCAC** in many organisms (Protostomes included)
    - **Molluscs_P1b_5p** seed shifted in 3p-direction to **UUUGGCA**

- **Gene Clusters:**
    - No known conserved gene clusters

- **In Pdu:**
    - **All 3 parlogues** are present.
        - **P1_5p** - C**uuggcac**UGGCGGAAUAAUCAC
            - **3p* -** gguguuccccugguggcaaaca

- Version 1 From the 2019SmallRNAseq-2020Mirdeep2-2019Genome
    - Shows **complete and almost complete allignment** with many organisms

```
> Cgi-Mir-96-P1i_5p
Length=23
Score = 43.6 bits (23),  Expect = 6e-07
 Identities = 23/23 (100%), Gaps = 0/23 (0%)
 Strand=Plus/Plus
Query  1   CTTGGCACTGGCGGAATAATCAC  23
           |||||||||||||||||||||||
Sbjct  1   CTTGGCACTGGCGGAATAATCAC  23

> Lgi-Mir-96-P1i_5p
Length=22
Score = 41.7 bits (22),  Expect = 2e-06
 Identities = 22/22 (100%), Gaps = 0/22 (0%)
 Strand=Plus/Plus
Query  1   CTTGGCACTGGCGGAATAATCA  22
           ||||||||||||||||||||||
Sbjct  1   CTTGGCACTGGCGGAATAATCA  22

> Lan-Mir-96-P1_5p
Length=23
Score = 38.1 bits (20),  Expect = 3e-05
 Identities = 22/23 (96%), Gaps = 0/23 (0%)
 Strand=Plus/Plus
Query  1   CTTGGCACTGGCGGAATAATCAC  23
           ||||||||||||||||| |||||
Sbjct  1   CTTGGCACTGGCGGAATGATCAC  23

> Cte-Mir-96-P1_5p
Length=23
```

Score = 38.1 bits (20), Expect = 3e-05
Identities = 22/23 (96%), Gaps = 0/23 (0%)
Strand=Plus/Plus
Query 1 CTTGGCACTGGCGGAATAATCAC 23
||||||||||||||||| |||||
Sbjct 1 CTTGGCACTGGCGGAATTATCAC 23

- 3p* shows only one partial allignment with Cte:

> Cte-Mir-96-P1_3p*
Length=22
Score = 25.1 bits (13), Expect = 0.19
Identities = 13/13 (100%), Gaps = 0/13 (0%)
Strand=Plus/Plus
Query 3 TGTTCCCCTGGTG 15
|||||||||||||
Sbjct 3 TGTTCCCCTGGTG 15

*From <https://mirgenedb.org/blast>*

-

- **P1_5p** - U**uuggcac**CAGUGGAAUAGUCAC
  - **3p* -** gaaauuccauuggugacauaug

- Version 2 from the 2019SmallRNAseq-2022Mirdeep2-2022Genome
  - Shows **almost complete allignment** with fewer organisms respect to version 1

> Cgi-Mir-96-P1h_5p
Length=23
Score = 23.3 bits (12), Expect = 0.80
Identities = 16/18 (89%), Gaps = 0/18 (0%)
Strand=Plus/Plus
Query 1 TTTGGCACCAGTGGAATA 18
|||||||| ||||||||
Sbjct 2 TTTGGCACTTGTGGAATA 19

- 3p* does not show any significant allignment with 3p* of other organisms

- **P2_5p** - C**uuggcac**UGGUAGAAUUCACUGA
  - **3p* -** agggauucuagagugcucaaaa
- **P3_5p** - A**auggcac**UGGUAGAAUUCACGG
  - **3p* -** gugggucuucuggugccgugca
- **2 other copies**, together in a cluster are also present. The paralogy relationship of these copies is not clear.
  - **Pdu-Mir-96-P1**(**bx** or **dx**)**_5p** - A**uuuggca**CUAGUGGAAUGGUC
    - **3p*** - gaguuucauuaaugccaacacc
  - **Pdu-Mir-96-P1**(**by** or **dy**)**_5p - A**uuuggca**CGAGUGGAAUAUUCG
    - **3p*** - guuauccaccugugccauauaa
  - The Seed is **UUUGGCA** for both. So it might be a shift to the 3'-direction, as it has happened in the P1b Moluscs' paralogue. However, probably the appearence of these copies is indipendent from the one of the molluscs, since is not detected in the other two Annelids. Still, it might be that P1 duplicated in Lophotrocoans into P1a and P1b (with P1b seed-shifting to UUUGGCA from UUGGCAC), P1b was then lost in the two Anellids (and maybe brachiopods), Cte and Efe, but further duplicated in Pdu (and maybe other anellids).
  - Efe instead duplicated the P2 paralogue in P2d and e.
- In Conclusion Pdu might have in total **5 copies** of the Mir-96 family of which:
  - **3 copies P1** of which:

- **Scenario 1:**
  **P1a, P1bx, P1by**
  P1a and P1b, originated from Lophotrocozoans. P1b then further duplicated in Pdu (and/or other Anellids) into P1bx and P1by
- **Scenario 2:**
  **P1c, P1dx, P1dy**
  The 3 copies appeared indipendently in *Pdum* from Molluscs (and/or other Lophotrocozoans). Probably in the following order. P1 duplicated into P1c and d. P1d seed-shifted and then further duplicated to P1dx and P1dy, forming a gene cluster.
- **P2**
- **P3**

- **Gene Clusters:**
  - A pdu specific **Mir-96-P1-Paralogues** might be present, containing:
    - Pdu-Mir-96-P1(bx or dx)_Pdu-Mir-96-P1(by or dy) - (within 1800ntds)

### Protostomia

- ## Bantam:
  - **Protostome-specific**
  - **Paralogues:**
    - Paralogues appeared indipenently just in some clades
    - Most Protostomes have just 1 copy
      - **Lan, Lgi, Cgi, Dme, Cte** posses just one copy
    - **Efe** has 3 copies:
      - **P6_3p**
      - **P7_3p**
      - **P8_3p**
  - **Seeds**
    - Most common **3p_mature_seed** is **GAGAUCA**
  - **Gene Clusters:**
    - No known conserve gene clusters
  - **Pdu:**
    - **2 Paralogues** in a cluster, appear to be present in Pdu.
      - There are no known relationship with paralogues of other species. Neither obvious relationships with the ones of the annelid Efe.

- **Pdu-Bantam-PX_3p -** U**gagauca**UGGUGAAAACUAAU
  - **5p* -** ugguuuucacaauggucuauaucaga
    Has almost complete identity to Cte-Bantam
    **> Cte-Bantam_3p**
    Length=23

    Score = 36.2 bits (19), Expect = 6e-05
    Identities = 21/22 (95%), Gaps = 0/22 (0%)
    Strand=Plus/Plus

```
Query  1   TGAGATCATGGTGAAAACTAAT  22
           ||||||||| ||||||||||||
Sbjct  1   TGAGATCATTGTGAAAACTAAT  22
```

**> Lgi-Bantam_3p**
Length=23

Score = 30.7 bits (16), Expect = 0.003
Identities = 20/22 (91%), Gaps = 0/22 (0%)
Strand=Plus/Plus

```
Query  1   TGAGATCATGGTGAAAACTAAT  22
           ||||||||  ||||||||||||
Sbjct  1   TGAGATCACTGTGAAAACTAAT  22
```

- 5p* also shows quite good alignments with many 5p* of other organisms

**> Lan-Bantam_5p***
Length=24
Score = 27.0 bits (14), Expect = 0.055
Identities = 14/14 (100%), Gaps = 0/14 (0%)
Strand=Plus/Plus

```
Query  1   TGGTTTTCACAATG  14
           ||||||||||||||
Sbjct  1   TGGTTTTCACAATG  14
```

**> Cgi-Bantam_5p***
Length=23
Score = 25.1 bits (13), Expect = 0.20
Identities = 15/16 (94%), Gaps = 0/16 (0%)
Strand=Plus/Plus

```
Query  1   TGGTTTTCACAATGGT  16
           ||||||||| ||||||
Sbjct  2   TGGTTTTCATAATGGT  17
```

- **Pdu-Bantam-PY_3p -** U**gagauca**UUGUGAAAACUGAUU
    - **5p* -** cugguuuucacauuggucuucc

  Shows complete identity to **Lan/Cgi/Tca/…-Bantam**

**> Lan-Bantam_3p**
Length=23

Score = 43.6 bits (23), Expect = 4e-07
Identities = 23/23 (100%), Gaps = 0/23 (0%)
Strand=Plus/Plus

```
Query  1   TGAGATCATTGTGAAAACTGATT  23
           |||||||||||||||||||||||
Sbjct  1   TGAGATCATTGTGAAAACTGATT  23
```

**> Cte-Bantam_3p**
Length=23

Score = 36.2 bits (19), Expect = 7e-05
Identities = 19/19 (100%), Gaps = 0/19 (0%)
Strand=Plus/Plus

```
Query  1   TGAGATCATTGTGAAAACT  19
           |||||||||||||||||||
Sbjct  1   TGAGATCATTGTGAAAACT  19
```

  5p* shows quite good identity with 5p* of other organisms

- **Gene Cluster:**

**Pdu-specific Bantam MiRNA cluster:**

- **Pdu-Bantam-PX_Pdu-Bantam-PY** within ~600ntds

- # Mir-1175:
  - **Prostostome-specific**
  - **Paralogues:**
    - Most Protostomes have **1 copy -** Mir-1175_5p
    - Few Protostomes started using **both arms -** Mir-1175_5p and Mir-1175_3p
    - **Efe** has **2 copies**:
      - **P1_5p**
      - **P2_5p**
  - **Seeds:**
    - **5p_mature -** GAGAUUC
    - **3p_mature -** AGUGGAG
  - **Gene Clusters:**
    - **Protostome-conserved** cluster: Mir-750_Mir-1175
      - **in Cte:** Mir-750_Mir-1175 (within 1200 ntds)
  - **Pdu:**
    - **Pdu-Mir-1175_3p -** U**gagauuc**AACUCCUCCAACUGC
      - **5p* -** aguggagagaguucuaucucauc

- **3p_mature** sequence shows **complete conservation** with other 5p_mature of many other protostomes.
- **5p*** sequence also shows **complete conservation** with many other 5p* of other protostomes: Lgi, Cte, Cgi, Efe and Lan
- Pdu does not seem to use also the 5p-arm as seen in some protostomes - 5p* read count is low in the 2020SmallRNAseq
- **Gene Cluster:**
  - **Protostome-conserved** cluster: Mir-750_Mir-1175
    - **in Pdu:** Mir-750_Mir-1175 (within 2050 ntds)

- # Mir-1993:
  - **Prostostome-specific**
  - **Paralogues:**
    - **All Prototstomes** have **1 gene - Mir-1993_3p**
    - **Efe** has to copies:
      - **P1_3p**
      - **P2_5p/3p**
  - **Seeds:**
    - **AUUAUGC** is the only **3p_seed**
    - **Efe** P2_5p_seed is CGGAAAU
  - **Gene Clusters:**
    - No known gene clusters

- **In Pdu:**
  - **Pdu-Mir-1993_3p** - U**auuaugc**UGUUAUUCACGAGA
    - **5p*** - ucgggaauaucaguguucuaugcc
    - 3p-mature shows **complete identity** with Molluscs and almost
    - complete with Lan and Cte

- **Gene Clusters:**
    - No known gene clusters

- ## Mir-277:
    - **Protostome-specific**
    - **Paralogues:**
        - **1 copy** for most of the species
        - **Cte** has **2 copies:**
            - **P1_3p**
            - **P2_3p**
                - Named the same has the ones of C.Elegans(?)
        - **Lan** has **3 copies:**
            - **P3_3p**
            - **P4_3p**
            - **P5_3p**
    - **Seeds:**
        - **3p_seed AAAUGCA** is the **only one** used by all species apparently
    - **Gene Clusters:**
        - **Protostome-conserved** cluster Mir-34_Mir-277_Mir-317
        - **In Cte:** Mir-34_~400ntds_Mi-277-P2_~100ntds_Mi-277-P1_~150ntds_Mir-317

- **In Pdu:**
  **1 copy** present in Pdu:

- **Pdu-Mir-277_3p** - U**aaaugca**UUAUCUGGUAUGUA
    - **5p* -** cauaucagaaaugcacuuugca

- **3p_mature** shows **complete identity** to Cte and Lan
- **5p*_star** is not conserved

- **Gene Clusters:**
    - **Protostome-conserved cluster** cluster Mir-34_Mir-277_Mir-317
    - Mir-317_~3500ntds_Mir-277_~26.000ntds_Mir-34

- ## Mir-278:
    - **Protostome-specific**
    - **Paralogues:**
        - **1 copy** in most of the species
        - **Efe** has **3 copies:**
            - **P1-v1_3p (+)**
            - **P1-v2_3p (-)**
            - **P2_3p**
    - **Seeds:**
        - **3p_seed CGGUGGG** is the most common seed with few exceptions
    - **Gene Clusters:**
        - No known conserved clusters

- **In Pdu:**
  **1 copy** is present

- **Pdu-Mir-278_3p -** U**cggugggg**ACUUUCGUUCGUUC
  - **5p* -** acgaaugaaauucucgacaggguc

- **3p_mature** shows **complete identity** with other species
- **5p*_star** is not conserved

- **Gene Clusters:**
  - No Known gene clusters

- ## Mir-279:
  - **Prostostome-specific**
  - **Paralogues:**
    - Indipendent duplications/paralogues appeared in several clades.
    - **Cte** has 3
      - **o11_3p**
      - **o12_3p**
      - **o13_3p**
    - **Efe** has 7
      - **o14_3p**
      - **o15_3p**
      - **o16_3p**
      - **o17_3p**
      - **o18_3p**
      - **o19_3p**
      - **o20_3p**
    - **Lan** has 2
      - **o40_3p**
      - **o41_3p**
    - **Cgi** and **Lgi** have 1 copy
  - **Seeds:**
    - Most common seed is the 3p_mature_seed **GACUAGA**
      - Very few exceptions
  - **Gene Clusters:**
  **Anellid-specific: Mir-36_Mir-279**
- **Cte:** Mir-36_Mir-279-o11 within ~2000 ntds

- **Pdu:**
  Might have **2 to 4 copies**
- **2 copies Mir-279-1** with different Star sequences:
  - **Pdu-Mir-279-1a_3p -** U**gacuaga**UCCACACUCAUCC
    - **5p*-(a) -** augacuguaggucuaguccaug - scaffold_9:2657710..2657763:-

- This one has great allignment with 5p* of few other organisms

**> Lan-Mir-279-o41_5p***
Length=22

Score = 25.1 bits (13), Expect = 0.13

```
 Identities = 19/22 (86%), Gaps = 0/22 (0%)
 Strand=Plus/Plus

Query  1   ATGACTGTAGGTCTAGTCCATG  22
           |||| |||  ||||||||||||
Sbjct  1   ATGATTGTGCGTCTAGTCCATG  22
```

- **Pdu-Mir-279-1b_3p -** U**gacuaga**UCCACACUCAUCC
  - **5p*-(b) -** ggugagugggauuuuggucaug - scaffold_1:184772430..184772489:-

- This one has just one good allignment with a 5p*

```
> Efe-Mir-279-o18_5p*
Length=23

 Score = 21.4 bits (11),  Expect = 1.6
 Identities = 18/21 (86%), Gaps = 1/21 (5%)
 Strand=Plus/Plus

Query  3   TGAGTGGGATTTTGGTC-ATG  22
           ||||||||||| || || |||
Sbjct  3   TGAGTGGGATTCTGTTCCATG  23
```

- **Gene Cluster:**
  - **Anellid-specific Mir-36_Mir-279.** Only found in Cte:
    **In pdu:** Mir-36_Mir-279 - within ~4800ntds

- **1-2 copy/copies of Pdu-Mir-279-2:**
  - 2 copies Mir-279-2 with different Star sequences:

- **Pdu-Mir-279-2a_5p-3p -** A**ugacugu**AGGUCUAGUCCAUG
  - **3p** - u**gacuaga**uacacacucaucc - scaffold_3:17716402..17716455:+

- **Arm-switch** from 3p to 5p-mature, thus completely changing seed.
  Has great allignments with few 3p of other organisms:

```
> Lan-Mir-279-o41_5p*
Length=22

 Score = 25.1 bits (13),  Expect = 0.13
 Identities = 19/22 (86%), Gaps = 0/22 (0%)
 Strand=Plus/Plus

Query  1   ATGACTGTAGGTCTAGTCCATG  22
           |||| |||  ||||||||||||
Sbjct  1   ATGATTGTGCGTCTAGTCCATG  22
```

- **Both arms** might **be active.** In the 2019SmallRNAseq-2020MirDeep2-2019Genome prediciotn **star read** and **mature read** are in the same order of magnitude. In many samples and developmental stages the 3p arm can be higher respect to 5p arm. Sometimes they have equivalent number or reads.

- **Pdu-Mir-279-2b_5p -** A**ugacugu**AGGUCUAGUCCAUG
  - **3p** - ugauuagucaaacacucauagc - **s**caffold_3:90160574..90160628:-

- Does not show any significant alignments with 3p of other organisms

- **Gene Clusters:**
    - No gene clusters found

- # Mir-12:
    - **Prostome-specific**
    - **Paralogues:**
        - **1 copy** is present in almost all clades, with few exceptions
        - **Efe** has 2
            - **P1_5p**
            - **P2_5p**
        - **Lan** has 2
            - **P3_5p**
            - **P4_5p**

**Seeds:**

- **5p_GAGUAU** is the only seed

- **Pdu:**
    - **Pdu-Mir-12_3p-5p** - C**gugccuu**UUGUGAUUCUCUUG
        - **5p(*)** - u**gaguauu**acaucagguacuga

- **Arm-switch** from 5p to **3p-mature** or **both-arms_mature.**
    - In the 2020SmallRNAseq, in many samples, the ttwo 5p and 3p sequences are in the same order of magnitude and sometimes equal (~ 5000 total sum of all samples)
- **5p-arm** also, has great allignment with several 5p of other organisms.

```
> Lgi-Mir-12_5p
Length=22
Score = 41.7 bits (22),  Expect = 1e-06
 Identities = 22/22 (100%), Gaps = 0/22 (0%)
 Strand=Plus/Plus
Query  1   TGAGTATTACATCAGGTACTGA  22
           ||||||||||||||||||||||
Sbjct  1   TGAGTATTACATCAGGTACTGA  22
```

- **Gene Cluster:**
    - **Protostome-conserved** Gene Cluster **Mir-12_Mir-216:**
        - **Mir-12_Mir-216-P1_Mir-216-P2**. (within ~2.300 ntds)

- # Mir-2:
    - **Protostome-specific**
    - **Paralogues**:

- It is not clear either from phylogenetic or syntenic information how many Mir-2 genes were present in the last common ancestor of protostomes and thus these multiple paralogues in invertebrates other than Drosophila are classified here as orphans pending further data and analysis.
- Cte has 8, named from o36 to o43
- Efe has 7, named from o44 to o50
- Pdu might have betwwen 8-10 genes
    - 8 or 9 have the AUCACAG seed
- It might well be that many of duplications present in Anellids already originated in the Annelid-last common ancestor or in the .
-

- **Seed:**
    - Most common 3p_mature seed is **AUCACAG**
    - Some have **seed-shift** to **5p-direction**: CACAGCC
    - Other variants are also present
- **Gene Clusters:**
    - Two Mir-2 clusters were "probably" already present in the Protostome-LCA with one of them containing also Mir-71.
    - **In Cte:**

**In Cte:**

- **Cluster 1 (Mir-2 + Mir-71):**
    - Contains 4 Mir-2 copies + Mir-71 within <600ntds
        - Mir-71_Mir-2-o37_Mir-2-o42_Mir-2-o38_Mir-2-o36
- **Cluster 2 (Mir-2):**
    - Is composed of 3 Mir-2 family genes:
        - Mir-2-o41_Mir-2-o43_Mir-2-o40 (within < 400ntds)
-
- **Pdu:**
    - Two **Mir-2** genes clusters containing different mir-2 copies are present in pdu:
    - **Nomenclature:** Capital Letters A,B,C,... indicate the position on the cluster.
        - Small letters in second and third position identify the unique mature sequence and star sequence respectively
        - The number 1 and 2 indentify the cluster.

- **Cluster 1:**
    - **Pdu-Mir-2-oAa1_3p -** U**aucacag**CGUGCUUUGAUGAGCU - scaffold_3:67088839..67088898:+
        - 5p* - cucgcgaagucgcugugccagg

- **Pdu-Mir-2-oBb1_3p-5p** - U**aucacag**CAGCGCUUUGAUGAGUC - scaffold_3:67088402..67088463:+
    - 5p* - ggcaucaaggguucugugaaaug

- **Both arms** might be **active** depending on **Developmental stage**: In the 2020SmallRNAseq-MirDeep2-2019Genome 5p reads are higher or equivalent to 3p in early stages (24-36h). 3p is around one order of magnitude higher in later stages (72hpf-6dpf).

- **Pdu-Mir-2-oCc-a1_3p** - U**aucacag**CCAGCUUUGAUGAGC - scaffold_3:67088097..67088157:+

- 5p* - cccaucuuaguggcuguggugug

- **Pdu-Mir-2-oDc-b1_3p** - U**aucacag**CCAGCUUUGAUGAGC - scaffold_3:67087900..67087958:+
  - 5p* - ccaucuaaguggcugugaugug

- Same mature sequence as  Pdu-Mir-2-oCc-a1_3p, but slightly different *star sequence

- **Pdu-Mir-2-oEd1_3p** - U**aucacag**CCAGCUUUGACGAGC - scaffold_3:67087490..67087548:+
  - 5p* - cgucacaguggcugugaagug
- **Pdu-Mir-2-oFe1_3p** - U**aucacag**CCUUGCUUGGAUCAUA - scaffold_3:67087095..67087160:+
  - 5p* - cgguccaagugguugagauagc
- **Pdu-Mir-2-oGc-c1_3p** - U**aucacag**CCAGCUUUGAUGAGC- scaffold_3:67086276..67086336:+
  - 5p* - caucaauccuggcauguuaua

- 
  - Pdu-Mir-2-oAa1_Pdu-Mir-2-oBb1_**Pdu-Mir-2-oCc-a1**_Pdu-Mir-2-oDc-b1_Pdu-Mir-2-oEd1_Pdu-Mir-2-oFe1-

- **Cluster 2:**
  - **Pdu-Mir-2-oA2_5p-3p** - U**ggccaag**GGGGCUGUGAUGUG - scaffold_3:66418561..66418621:+
    - **3p*** - uaucacagcccaugcuuuggucaaa

- Arm-shift from 3p to **5p_mature**, or both
  - From the 2019SmallRNAseq this is not clear. 3p and 5p reads are in the same order of magnitude. It looks like that in early stages (24-36-48hpf) the 5p read is more abbundant, but in 72hpf and 6dpf the reads are more or less equivalent. However, this needs to be further investigated. it might be that both arms are active
- The **5p_seed** is **not conserved** or found in other organisms
- 5p_mature shows some good allignments with few other 5p*_star of other species

```
> Lan-Mir-2-o76_5p*
Length=22
Score = 34.4 bits (18),  Expect = 2e-04
 Identities = 18/18 (100%), Gaps = 0/18 (0%)
 Strand=Plus/Plus
Query  5   CAAGGGGGCTGTGATGTG  22
           ||||||||||||||||||
Sbjct  5   CAAGGGGGCTGTGATGTG  22

> Cte-Mir-2-o36_3p
Length=23
Score = 21.4 bits (11),  Expect = 1.6
 Identities = 13/14 (93%), Gaps = 0/14 (0%)
 Strand=Plus/Minus
```

```
Query  6   AAGGGGGCTGTGAT  19
           ||| ||||||||||
Sbjct  15  AAGCGGGCTGTGAT  2
```

- 3p*_star shows good allignments with many others 5p_mature of other species

```
> Cte-Mir-2-o39_3p
Length=23
Score = 28.8 bits (15),  Expect = 0.014
 Identities = 20/22 (91%), Gaps = 2/22 (9%)
 Strand=Plus/Plus
Query  2   ATCACAGCCCATGCTTTGGTCA  23
           |||||||||  |||||||||||
Sbjct  2   ATCACAGCC--TGCTTTGGTCA  21

> Cte-Mir-2-o40_3p
Length=24
Score = 27.0 bits (14),  Expect = 0.050
 Identities = 18/20 (90%), Gaps = 0/20 (0%)
 Strand=Plus/Plus
Query  1   TATCACAGCCCATGCTTTGG  20
           |||||||| | |||||||||
Sbjct  1   TATCACAGTCAATGCTTTGG  20

> Tca-Mir-2-o52-v1_3p
Length=25
Score = 23.3 bits (12),  Expect = 0.65
 Identities = 17/19 (89%), Gaps = 2/19 (11%)
 Strand=Plus/Plus
Query  1   TATCACAGCCCATGCTTTG  19
           |||||||| ||| ||||||
Sbjct  1   TATCACAG-CCA-GCTTTG  17
```

- **Pdu-Mir-2-oB2_3p -** U**aucacag**CCCGCUUUGACAAGU - scaffold_3:66418750..66418810:+
  - 5p* - guguugaagugguugugaugug
- **Pdu-Mir-2-oC2_3p** - U**aucacag**CCAAUUUGACAGCC - scaffold_3:66419071..66419128:+
  - 5p* - cuguuggauuuguugugacuaug
- **Pdu-Mir-2-oD2_3p** - U**aucacag**CCCGCUUUGGUCAUA - scaffold_3:66419294..66419351:+
  - 5p* - cggucaaaguggcugagauaug
- **Pdu-Mir-2-oE2_3p** - U**aucacag**CCAGAUUUGGUAUAUC - scaffold_3:66419578..66419640:+
  - 5p* - agccaaaucggcugugaaaug

- **Gene Clusters:**
  - The 2 clusters sit on the same scaffold 10^6ntds apart
  - **Cluster 1/2 + Mir-71:**
    - **Is composed of 7 Mir-2 family genes together with Mir-71:**
      - **Pdu-Mir-2-oAa1_Pdu-Mir-2-oBb1_Pdu-Mir-2-oCc-a1_Pdu-Mir-2-oDc-b1_Pdu-Mir-2-oEd1_Pdu-Mir-2-oFe1_Pdu-Mir-2-oGc-c1_~1000ntds_Mir-71**

Mir-2 genes all within ~2500ntds.

- This Cluster containing Mir-71 is conserved in Protostomes.

- **Cluster 2/2:**
  - **Is composed of 5 Mir-2 family genes.**

        - **Pdu-Mir-2-oA2_Pdu-Mir-2-oB2_Pdu-Mir-2-oC2_Pdu-Mir-2-oD2_Pdu-Mir-2-oE2**
            - **All within 1000ntds**

- Also Cte presents 2 Clusters of mir-2 genes

- # Mir-317 Family:
    - **Protostome-specific**
    - **Orthologues and Paralogues:**
        - Most Protostomes appear to have only one copy
        - **Cgi** has 2 copies**:**
            - P1_3p
            - P2_3p
        - **Lan** has 2 copies:
            - P10_3p
            - P11_3p
        - **Efe** has 3 copies:
            - P5_3p
            - P6_3p
            - P7_3p
    - **Seeds:**
        - **3p_seed** GAACACA is the only seed present

- **Gene clusters:**
    - **Protostome conserved** cluster Mir-34_Mir-277-Mir-317:
        - **In Cte:** Mir-34_~400ntds_Mi-277-P2_~100ntds_Mi-277-P1_~150ntds_Mir-317
- **In Pdu:**
    - **Pdu-Mir-317_3p -** U**gaacaca**GCUGGUGGUAUCUCUUU
        - 5p* - agaguacuccagcguggucgca
    -
    - **Gene Clusters:**
        - **Protostome-specific gene cluster** Mir-34_Mir-277-Mir-317
        - Mir-317_~3500ntds_Mir-277_~26.000ntds_Mir-34

- # Mir-36 Family:
    - **Protostome-specific**
    - **Orthologues/Paralogues:**
        - Not present in all Protostomes, probably due to sencondary loss. In Lophotrochozoans prensent in Annelids Cte and Efe and in the Mollusc Cgi.
            - Cte has **1 copy**
            - Efe has **2 copies**
                - **P9_3p**
                - **P10_3p**
    - **Seed:**
        - **3p_seed** CACCGGG is the main one with few exceptions
        - **3p_mature** is the only on e used with few exceptions

- **Gene Clusters:**
  **Anellid-specific: Mir-36_Mir-279**

- **Cte:** Mir-36_Mir-279-o11 within ~2000 ntds

- **In Pdu:**
    - **Pdu-Mir-36_5p-3p 5p -** G**gugagug**GAUACUCAGGUGGUG **3p -** U**caccggg**UAUACAUUCAUCCG

- Only the **3p_arm** shows complete alignments with many Mir-36_3p of other organisms
- The **5p_arm** does not show any significant allignments
- In the 2019SmallRNAseq the 5p and 3p_arm read counts are in the same order of magnitude. The 3p_arm come be slightly more abbundant or equivalentto 5p in early stages (24-48hpf). However, in later stages (72hpf-6dpf) the 5p_arm is more abundant (around 5-fold higher or more)

- > Asu-Mir-36-o27_3p

```
Length=22

 Score = 38.1 bits (20),  Expect = 2e-05
 Identities = 20/20 (100%), Gaps = 0/20 (0%)
 Strand=Plus/Plus

Query  1   TCACCGGGTATACATTCATC  20
           ||||||||||||||||||||
Sbjct  1   TCACCGGGTATACATTCATC  20
```

```
> Cgi-Mir-36_3p
Length=23
Score = 36.2 bits (19),  Expect = 6e-05
 Identities = 21/22 (95%), Gaps = 0/22 (0%)
 Strand=Plus/Plus
Query  1  TCACCGGGTATACATTCATCCG  22
          |||||||||| |||||||||||
Sbjct  1  TCACCGGGTAAACATTCATCCG  22
```

```
> Cte-Mir-36_3p
Length=22
Score = 30.7 bits (16),  Expect = 0.003
 Identities = 20/22 (91%), Gaps = 0/22 (0%)
 Strand=Plus/Plus
Query  1  TCACCGGGTATACATTCATCCG  22
          |||||||||  |||||||||||
Sbjct  1  TCACCGGGTTAACATTCATCCG  22
```

- **Gene Cluster:**
    - **Anellid-specific Mir-36_Mir-279.** Only found in Cte:
      **In pdu:** Mir-36_Mir-279 - within ~4800ntds

- # Mir-67:
    - **Protostome-specific**
    - **Orthologues and Paralogues:**
        - Most annotated Protostomes show only one copy of the gene
        - **Efe** shows 3 copies:
            - P1_3p
            - P2_3p/5p
            - P3_3p
        - **Cgi** has 2 copies:
            - P5_3p
            - P6_3p

- **Seeds:**
    - **3p_seed** CACAACC is the most common
        - Some organisms show seed shifts of this seed
    - Some organisms arm_shifted to **5p_mature**

- **Gene Clusters:**
    - No Known gene clusters
- **In Pdu:**
    - **Pdu-Mir-67_3p_5p - 3p_**UcacaaccUGCAUGAAUGAGGUA
        - 5p_accuuauucagugguguuguggu

- in the 2019SmallRNAseq the 3p and 5p_arms read are in the same order of magnitude. The proportion of the two varies according to the developmental stage, even if the reads are always in the same order of magnitude. Mainly at 24hpf the 3p can be 2-folds lower than 5p. In other stages, 3p it is equivalent or 2-3 folds higher respect to 5p.

- **Gene Clusters:**
    - No gene clusters found

- # Mir-750:
    - **Protostome-specific**
    - **Orthologues and Paralogues:**
        - Most of the organisms show only one copy of the gene
        - **Efe** has **3 copies:**
            - P1_3p
            - P2_3p
            - P3_3p

- **Seeds:**
    - **3p_seed** CAGAUCU is the only seed present

- **Gene Clusters:**
    - Protostomes conserved cluster Mir-750_Mir-1175
        - In Cte: Mir-750_Mir-1175 (within 1200 ntds)
        -

- **In Pdu:**
    - **Pdu-Mir-750_3p -** C**cagaucu**AACUCUUCCAGCUCA
        - 5p* - cgcuggaagcuuaggucuccgca

- **3p_mature** sequence shows **complete allignment** with other organisms

- **Gene Clusters:**
    - Protostomes conserved cluster Mir-750_Mir-1175
    in Pdu: Mir-750_Mir-1175 (within 2050 ntds)

- # Mir-87:
    - **Protostome-specific**

- **Orthologues and Paralogues:**
  - Most of the Protostomes have different copies of the gene
    - However, it is not clear either from phylogenetic or syntenic information how many Mir-87 genes were present in the last common ancestor of protostomes and thus these multiple paralogues are classified here as orphans pending additional data.
  - **Dme**, **Dan** and **Dmo** have **2 copies**
    - **v1_3p**
    - **v2_3p**
  - **Lgi** has **1 copy**
  - **Cgi** has **2 copies:**
    - **o7_3p**
    - **o8_3p**
  - **Lan** has **2 copies:**
    - **o25_3p**
    - **o26_3p**
  - **Cte** has **2 copies:**
    - **o9_3p**
    - **o10_3p**
  - **Efe** has **7 copies:**
    - **o11_5p/3p**
    - **o12_3p**
    - **o13_3p**
    - **o14_3p**
    - **o15_3p**
    - **o16_3p**
    - **o17_3p**
- **Seeds:**
  - **3p_seed UGAGCAA** is almost ubiquitous.
  - Few show **arm-shift** to **5p_mature**

- **Gene Clusters:**
  - No known conserved gene clusters. However, in many organisms the gene copies are in cluster:
    - In Cte, Cgi, Isc, Dpu, Drosophilas, Tca, Lan.

- **In Pdu:**
  - Pdu-Mir-87_3p - G**ugagcaa**AGUUUCAGGUGUGU
    - 5p* - gugagcaaaguuucaggugugu

- **Complete allignment** of both 3p - 5p with many other species

- **Gene Clusters:**

  No gene cluster found

### Lophotrochozoa

- # Mir-1986 Family:
  - **Lophotrochozoa-specific** family
  - **Paralogues:**
    - Only **1 copy** in lophotrochozoans Lgi, Cgi, Lan, Cte
    - **Efe** appears to have **lost** the gene

- **Seeds:**
  - **3p_seed** is **GGAUUUC**
  - **In Cte** the gene is **only predicted**, so it is not known which mature sequence is used.
- **Gene Clusters:**
  - **Lophotrocozoan-specific** miRNA Mir-1990 paralogues and Mir-1986:
    - **In Cte:** Mir-1990-P2_<150ntds_Mir-2692_~6000ntds_Mir-1990-P1_~1000ntds_Mir-1986

- **In Pdu:**
  - **1 copy** of the gene is present.
  - **Arm-shift** respect to others from **3p to 5p_mature** or **both-arms active.**
  - In the 2019SmallRNAseq-MirDeep2-2019Genome prediction the reads counts have a difference of less than a order of magnitude order of magnitude. This trend is seen almost in all the developmental stages.

- **Pdu-Mir-1986-5p -** C**aaggauc**CUGGGGAAGCCCCC
  - 3p* - uggauuucccaugauccguaac

- **5p_ mature** shows partial alignment only with Lan-5p*

```
> Lan-Mir-1986_5p*
Length=22

 Score = 27.0 bits (14),  Expect = 0.035
 Identities = 16/17 (94%), Gaps = 0/17 (0%)
 Strand=Plus/Plus

Query  3   AGGATCCTGGGGAAGCC  19
           ||| |||||||||||||
Sbjct  3   AGGGTCCTGGGGAAGCC  19
```

- **3p*_star** shows complete alignment with 3p_mature of other Lophotrochozoans

```
> Cte-Mir-1986_3p
Length=22

 Score = 41.7 bits (22),  Expect = 1e-06
 Identities = 22/22 (100%), Gaps = 0/22 (0%)
 Strand=Plus/Plus

Query  1   TGGATTTCCCATGATCCGTAAC  22
           ||||||||||||||||||||||
Sbjct  1   TGGATTTCCCATGATCCGTAAC  22
```

- **Gene Clusters:**
  - Conserved Lophotrocozoan-specific **gene cluster:**
    - Mir-1986_~1500ntds_Mir-1990-(+/-)_~1500ntds_Mir-2692

- 
- # Mir-1989:
    - **Platytrochozoa-specific:**

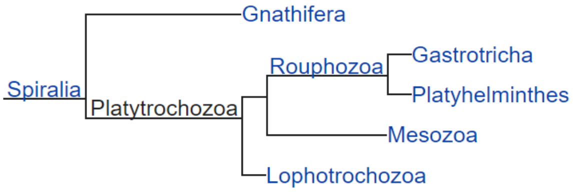


- **Paralogues and Orthologues:**
    - **Many Lophotrocozoa** appear to have **1 copy: Mir-1989_5p**
        - **Cgi**, **Lgi**, **Cte**
    - **Lan** has **3 copies:**
        - **P1_5p**
        - **P2_5p**
        - **P3_5p**
    - **Efe** probably lost it.
- **Seed:**
    - 5p_CAGCUGU is the **only seed** known
- **Gene Clusters:**
    - Not known conserved gene clusters
- **Pdu:**

Pdu might have in total 5 copies of the Mir-1989:

- All share the same 5p_seed-CAGCUGU.
- 5p_mature are slightly different.
- 3p* sequences are all different between the 5 genes.
- They are organized in 1 Cluster.

- The genes are named this way:
    - 1, 2, 3, 4 ...: stands for the order of the gene in the cluster
    - a, b, c, d: indentify the 3 different 5p_mature sequences

- **Pdu-Mir-1989-1-a_5p - Ucagcugu**AAAGAUGCCUUCUU
    - **3p*** - ggagccauuuuaacugcugaca

- 5p_mature shows **almost complete identity** with all other Lophotrochozoans

- **Pdu-Mir-1989-2-a_5p - Ucagcugu**AAAGAUGCCUUCUU
    - **3p*** - ggagucauuuuaacugcugaca

- **Pdu-Mir-1989-3-a_5p - Ucagcugu**AAAGAUGCCUUCUU - Version1_2022Genome
    - **3p*** - gagacauuuuaacugcugacau

- **Pdu-Mir-1989-4-b_5p - Ucagcugu**AACGAUGCCUUCUU
    - **3p*** - ggagucaucuuaacugcugaca

- 5p_mature shows **almost complete identity** with all other Lophotrochozoans

- **Pdu-Mir-1989-5-c_5p -** U**cagcugu**CACGAUGCCUUCUU
    - **3p*** - gggggcauuuuaacugcugaca

- 5p_mature shows **complete identity** and **almost complete identity** with all other Lophotrochozoans

- **Pdu-Mir-1989-6-d_5p** - U**cagcugu**CGCGAUGCCUUCUU
    - 3p* - gaagguguuaccacugcugaca

- 5p_mature shows **almost complete identity** with other Lophotrochozoans

- **Gene Clusters:**
    - Platynereis-specific Mir-1989 genes cluster:
      Mir-1989-1-a_~200ntds_2-a_~300ntds_3-a_~300ntds_4-b_~400ntds_5-c_~450ntds_6-d

- ## Mir-1990:
    - **Lophotrocozoa-specific**
    - **Paralogues:**
        - **Mir-1990_3p - 1 Gene** in **Lophotrocozoa** (some **molluscs** and **brachiopods**).
            - **3p arm** is the **mature** sequence (seed GGGACUA)
            - 5p*-seed would be "aGUAAGUUga"
        - **Mir-1990_5p - 1 Gene** in Lgi.
            - **5p arm** is the **mature** sequence (seed UAAGUUG) G**uaaguug**AUGGGGUCCCAGG
            - In this ca Lgi is using the 5p arm
            - 3p*-seed would be "GGGACUA"

          C**gggacua**CGUCAACUACUAG

- **P1_5p and P2_5p** paralogues are **Annelid-specific**
  Both present in Cte and Efe.
- **P1_5p:**
    - **5p arm is the mature sequence (seed GUAAGUU).**
      In P1 there was, probably, a shift from the ?"ancestral-Lophotrocozoan"? 3p arm to the 5p arm (the 3p* arm, in fact, still shows the *seed GGGACUA in P1).
- **P2_5p:**
    - **5p arm is the mature sequence (seed AAGUUGA).**
    - **3p-*seed would be "aCUUGUGGuua"**
      Also in P2 there was, probably, a shift from the ?"ancestral-Lophotrocozoan"? 3p arm (however, it shows a different 3p*_seed CUUGUGG) to the 5p arm.
- Respect to P1_5p (and 1990_5p*) it shows a seed-shift in 5'-direction. This would also explain the change in the 3p*_seed.
- **P3_3p:** is **Efe-specific** paralogue.
    - **3p arm** is the **mature** (seed "CGGGACU")

    - P3, probably, kept the ?"ancestral-Lophotrocozoan"? 3p arm with a seed-shift in 3'-direction.
    - 5p*-seed is GUAAGUU. Which is the same 5p seed of the other paralogues.
- **Seeds:**
    - **Cgi/Lan-Mir-1990_3p** - GGGACUA
    - **P1_5p -** GUAAGUU
    - **P2_5p** - AAGUUGA
    - **Efe-P3_3p** - CGGGACU

- **Gene clusters:**
    - **Mir-1990-P2_Mir-2692_Mir-1990-P1_Mir-1986** Cluster in Cte

- **Pdu:**
    - Appears to have 2/3 copies of the gene.
        - **Copy 1**
          Both strands of the gene appear to be active, giving rise to:
- **Pdu-Mir-1990-1orP1_3p (+)** (U**gggacua**UGUCAACUUACAAC)
    - 5p* - uggaaguuaacguagucccggg
    - This one has complete (+/-)-identity with Efe-1990-P1_5p

```
> Efe-Mir-1990-P1_5p
Length=22
Score = 38.1 bits (20),  Expect = 2e-05
 Identities = 20/20 (100%), Gaps = 0/20 (0%)
 Strand=Plus/Minus
Query  1   TGGGACTATGTCAACTTACA  20
           ||||||||||||||||||||
Sbjct  20  TGGGACTATGTCAACTTACA  1
```

- **3p-Seed:** is equal to the one found in Molluscs' paralogues
- **Pdu-Mir-1990-1orP1_5p-3p _-_** U**guaaguu**GACAUAGUCCCAGG
    - **3p* - cgggacuacguuaacuuccagc**

- **Both-arms** might **be active.**
- In the 2019SmallRNAseq-MirDeep2-2019Genome prediction the reads counts have a difference of less than a order of magnitude. This trend is seen almost in all the developmental stages and samples.

- This one has complete identity with Efe-Mir-1990-P1_5p

```
> Efe-Mir-1990-P1_5p
Length=22Score = 41.7 bits (22),  Expect = 1e-06
 Identities = 22/22 (100%), Gaps = 0/22 (0%)
 Strand=Plus/Plus
Query 1   TGTAAGTTGACATAGTCCCAGG  22
          ||||||||||||||||||||||
Sbjct 1   TGTAAGTTGACATAGTCCCAGG  22
```

**Seed:** Equal to the P1 paralogue of Annelids

- The identity with Cte-P2_5p is still high, but with a seed shift.

```
> Cte-Mir-1990-P2_5p
Length=22

 Score = 32.5 bits (17),  Expect = 8e-04
 Identities = 19/20 (95%), Gaps = 0/20 (0%)
 Strand=Plus/Plus

Query  3   TAAGTTGACATAGTCCCAGG  22
           ||||||||| ||||||||||
```

Sbjct 1 TAAGTTGACGTAGTCCCAGG 20

- **Gene Cluster:** in the conserved Annelid gene cluster of **Mir-1990_Mir-2692_Mir-1986.** That in Pdu is organized in this way:
    - **Mir-1986_Mir-1990-(+/-)_Mir-2692**
- **Copy 2:**
    - **Pdu-Mir-1990-2orP2_5p** (U**guaaguu**AACGUAGUCCAAGGUU)

        This one has almost complete identity with Cte-Mir-1990-P1_5p:

        **> Efe-Mir-1990-P1_5p**
        Length=22
        Score = 25.1 bits (13), Expect = 0.16
        Identities = 19/22 (86%), Gaps = 0/22 (0%)
        Strand=Plus/Plus
        Query 1 TGTAAGTTAACGTAGTCCAAGG 22
        |||||||| || |||||| |||
        Sbjct 1 TGTAAGTTGACATAGTCCCAGG 22

        The identity with Cte-P2_5p is also quite high. Almost equivalent to P1, but with a seed shift.

        **> Cte-Mir-1990-P2_5p**
        Length=22

        Score = 27.0 bits (14), Expect = 0.045
        Identities = 18/20 (90%), Gaps = 0/20 (0%)
        Strand=Plus/Plus

        Query 3 TAAGTTAACGTAGTCCAAGG 22
        |||||| ||||||||| |||
        Sbjct 1 TAAGTTGACGTAGTCCCAGG 20

        **Seed:** Equal to the P1 paralogue of Annelids

- It is not clear wether this duplication in Pdu is related to the one of the other two annelids or is indipendent. The two genes are not located in the same gene cluster, like in *Cte*. However, since the genes order in the cluster is also different, it might be that genomic changes in the locus (translocations) might have changed the position of one of the copies.

○ **Gene Cluster**
    - Conserved Annelid gene cluster of **Mir-1990_Mir-2692_Mir-1986**
        - That in Pdu is organized in this way:
        **Mir-1986_Mir-1990-1orP1(+/-)_Mir-2692**
- In Cte is:
        **Mir-1990-P2_Mir-2692_Mir-1990-P1_Mir-1986**

- # Mir-1992:
    - **Platytrochozoa-specific [Spiralian sister group to Gnathifera])**

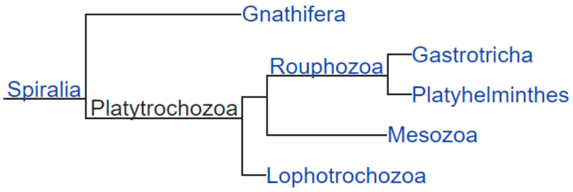

- **Paralogues:**
    - **Cte, Lan** and **Cgi** have **one copy**.
    - **Lgi** have **2 copies**:
        - **P1_3p**
        - **P2_3p**
    - **Efe** has **2 copies:**
        - **P3_3p**
        - **P4_3p**
- **Seeds:**
    - **3p_seed CAGCAGU** is the only one used
- **Gene Clusters:**
    - No known conserved gene clusters

- **In Pdu:**
    - **2** copies in a cluster might be present:
    - **Arm-shift** from **3p to 5p_mature** or **both arms** are **used as mature** sequences. Since in the 2019SmallRNAseq-MirDeep2 the two sequences have reads in the same order of magnitude in all developmental stages.
    - MirDeep2 finds two overlapping hairpins for this gene:
        - The second having the 5p_sequence starting at the 3p_sequence of the first one.However, for the first one 5p_star sequence is only predicted. So it might be an artifact.

**I Hairpin -------------------------------------------------------------------------------**
**cucagcacuaauccaucugug**cgucuuccugacucuggcacuggcagca**cauuaguggauaacggcuggua**guaacauuacgu
cugccua**ucagcaguuguaccacugaugug**
II Haripin---------------------------------------
-----------------------------------------

- **Pdu-Mir-1992-v1_5p-3p -** C**auuagug**GAUAACGGCUGGUA
    - **3p -** U**cagcagu**UGUACCACUGAUGUG

- **3p(*)_sequence** shows **complete alignment** with Cte_3p and almost complete alignments with all other lophotrochozoans.
- **5p(m)_sequence** does not show any significant alignment.

> Cte-Mir-1992_3p
Length=23
Score = 43.6 bits (23), Expect = 4e-07
Identities = 23/23 (100%), Gaps = 0/23 (0%)
Strand=Plus/Plus
Query 1 TCAGCAGTTGTACCACTGATGTG 23
|||||||||||||||||||||||
Sbjct 1 TCAGCAGTTGTACCACTGATGTG 23

- **Pdu-Mir-1992-v2_3p -** C**auuagug**GAUAACGGCUGGUA
    - **5p*(predicted)-** cucagcacuaauccaucugug

- **Gene Clusters:**
    - If both hairpins are transcribed, this might be considered a cluster

- ## Mir-1994:
    - **Lophotrocozoa-specific**
    - **Paralogues:**
        - **Annelids (Cte and Efe)** have **1 gene -** Mir-1994_3p
        - **Molluscs** (Lgi, Cgi) have **2 copies:**
            - **P1_3p**
            - **P2_3p**
        - **Lan** has **4 copies:**
            - **P3a**
            - **P3b**
            - **P4a**
            - **P4b**
    - **Seeds:**
        - GAGACAG is th only 3p_seed
    - **Gene Clusters:**
        - No known conserved gene clusters

- **In Pdu:**
    - **Pdu-Mir-1994_3p -** U**gagacag**UGUGUCCUCCCUCG
        - **5p*** -  caggaggacaugcuaucuucauc
        - 3p_mature shows **complete identity** with other lophotrocozoans
    - **Gene Clusters:**
        - No known gene clusters

### Annelida

- ## Mir-1987:
    - **Annelid-specific**
    - **Orthologues and Paralogues**:
        - **1 copy** present in both **Efe** and **Cte - Mir-1987_3p**
    - **Seed:**
        - **3p_mature -** seed CUGCCAG
            - Is the only seed
    - **Gene Clusters:**
        - **In Cte:** Mir-2705_Mir-1987 (within ~1000 ntds)
    -
    - **Pdu:**
        - **Pdu-Mir-1987_3p-5p -** A**cugccag**AUGUCAUGUUGUGCA
            - **5p -** cauaacuggcaccugggcauguug

- Probably **both arms** are **active.** However, in the2019SmallRNAseq-MirDeep2-2019Genome it looks like there is a change in the ratio from early to late stages. In early stages (24-36 hpf) the abundancy of the 5p is to order higher respct to 3p. However, later stages 72hpf-6dpf, 3p is higher but the difference is less pronounced

        - 3p_mature shows **almost complete identity** with **Cte. Good identity** with **Efe**
        - 5p* does not show significant alignments with related miRNAs
- **Gene Clusters**
    - **Annelid-conserved gene cluster:** Mir-2705_Mir-1987 (within ~5000)

- # Mir-1995:
    - **Annelid-specific:**
    - **Orthologues and Paralogues**:
        - **Cte** has **1 gene -** Cte-Mir-1995_5p
        - **Efe** has **2 copies:**
            - **P1_5p**
            - **P2_5p**
    - **Seeds:**
        - **UACAUCU -** is the only **5p_seed**
    - **Gene Clusters:**
        - No known conserved gene clusters

- **In Pdu:**
    - **Pdu-Mir-1995_5p-3p -** G**uacaucu**CACAUUGUGACCAU
        - **3p*** - ggucucugugugaaaugucgau

- 5p_mature shows almost **complete identity** to Cte

- **Both-arms** might **be active. But 5p** is higher than an order of magnitude in late stage (6dpf)
  In the 2019SmallRNAseq-MirDeep2-2019Genome prediction the reads counts have a difference of less than a order of magnitude order of magnitude. This trend is seen mostly in early stages: at 24hpf there is almost no difference between 3p and 5p reads. However, in later stages (6dpf) the difference between 5p and 3p is higher than an order of magnitude in all the developmental stages and samples.

- **Gene Clusters:**
    - No known gene clusters

- # Mir-1996:
    - **Annelid-specific:**
    - **Orthologues and Paralogues**:
        - **Cte** has **1 gene -** Cte-Mir-1996_5p
        - **Efe** has **3 copies:**
            - **P1_5p**
            - **P2_5p**
            - **P3_5p**
    - **Seeds:**
        - **UCAAGUG** is the only **5p_seed**
    - **Gene Clusters:**
        - No known conserved gene clusters

- **In Pdu:**

- Pdu has probably an **indiendent duplication** (from *Efe)* of the Mir-1996 locus, for which **two paralogues** (with the same seed, but different at the 3'-end) in the same cluster are present.
-
- **Pdu-Mir-1996-1_5p** - C**ucaagug**AGGUCAGUGCUACU
  - **3p*** - gggcacugcccucaacugaagc
  - 5p_mature shows **almost complete identity** with Cte and Efe-P1

- **Pdu-Mir-1996-2_5p -** A**ucaagug**AGGUCAGAUCUUGG
  - **3p*** - agggucauggucucaauugaccc
  - 5p-mature shows quite good identity with Efe paralogues and Cte (ntds 2-15). Low identity at the 3'-end of the sequence (ntds 15-22).

- **Gene Clusters:**
  - **Pdu-specific** Mir-1996 paralogues cluster:
    **Pdu-Mir-1996-1-5p_Pdu-Mir-1996-2_5p** within 800ntds

- # Mir-1997:
  - **Annelid-specific:**
  - **Orthologues and Paralogues**:
    - **Cte** has **1 gene** - Cte-Mir-1997_3p
    - **Efe** has **5 copies:**
      - **P1_3p**
      - **P2_3p**
      - **P3_3p**
      - **P4_5p**
      - **P5_3p**
  - **Seeds:**
    - **3p_seed** CUGCAGG
    - **Efe-P4_5p**_seed GGCUGAU
  - **Gene Clusters:**
    - No known conserved gene clusters
  - **In Pdu:**
    - **Pdu-Mir-1997_3p -** U**cugcagg**UUCACAUCAGCCCCA
      - **3p* -** gggcgaugguucucgcagggu
      - 3p_mature has **complete identity** to Cte and almost complete identity to all 3p_mature paralogues of Efe.

- # Mir-1998:
  - **Annelid-specific**
  - **Paralogues:**
    - **Cte** has 1 gene :**Mir-1998_3p**
    - **P1_3p** and **P2_3p** in Efe (seed UGAACGC)
  - **Seeds:**
    - **Cte-Mir-1998_3p** - UUGAACG
    - **Efe P1/P2_3p -** UGAACGC
  - **Cluster:**
    - **njjjAnellid conserved** cluster **Mir-2000_~200ntds_Mir-1998.** Conserved in Cte

- **Pdu:**
  - **Pdu-Mir-1998_3p** - U**ugaacgc**AGAGAUGUACAUCA
    - **5p*** - auguauaucccuacguccacacu
    - **Seed: UGAACGC** is equal to the one of P1/2 paralogues of Efe. Which have a Seed-shift, respect to Cte, of 1 ntd in 5p-direction.
    - No significant allignments of the 5p*

- **Gene Clusters:**
  - **Anellid conserved** Cluster **Mir-2000_~6000ntds_Mir-1998**. Conserved in Cte.

- # Mir-2000:
  - **Annelid-specific**
  - **Paralogues:**
    - **Cte** has **1 copy**
      - **Cte-Mir-2000_3p**
    - **Efe** has **2 copies**
      - **P1_3p**
      - **P2_3p**
  - **Seeds:**
    - **3p-mature** AAGUCUU
  - **Gene Clusters:**
    - **Anellid conserved** cluster **Mir-2000_~200ntds_Mir-1998.** Conserved in Cte

- **Pdu:**
  - **Pdu-Mir-2000_5p** - U**gacuaaa**AGUAUUGAAGGCUU
    - **3p*** - **gucuu**cacuacuuuuaguuugg
    - **Arm-shift** from 3p to **5p-mature**
    - 3p* arm also shows great allignments with 3p-mature of the other annelids

**> Efe-Mir-2000-P2_3p**
Length=22

```
 Score = 30.7 bits (16),  Expect = 0.003
 Identities = 18/19 (95%), Gaps = 0/19 (0%)
 Strand=Plus/Plus

Query  1   GTCTTCACTACTTTTAGTT  19
           |||||| ||||||||||||
Sbjct  4   GTCTTCTCTACTTTTAGTT  22
```

**> Cte-Mir-2000_3p**
Length=22

```
 Score = 30.7 bits (16),  Expect = 0.003
 Identities = 18/19 (95%), Gaps = 0/19 (0%)
 Strand=Plus/Plus

Query  1   GTCTTCACTACTTTTAGTT  19
           |||||||||||||| ||||
Sbjct  4   GTCTTCACTACTTTCAGTT  22
```

- **Gene Clusters:**

- **Anellid conserved** Cluster **Mir-2000_~6000ntds_Mir-1998**. Conserved in Cte.

- # Mir-2685 family:
  - **Annelid-specific** family
  - **Paralogues:**
    - **2 paralogues** in **Cte** and **Efe:**
      - **P1_3p**
      - **P2_3p**
    - **Efe** has a duplication of **P2:**
      - **P2a_3p**
      - **P2b_3p**
  - **Seeds:**
    - **3p_seed AACUCAG** is th only one
  - **Gene Clusters:**
    - Annelid-specific (probably **Sedentaria-specific)** Mir-2685-Paralogues cluster
    - **In Cte: Mir-2685-P1_Mir-2685-P2**

- **In Pdu:**
  - One copy of the gene detected in Pdu:
    - **Pdu-Mir-2685_3p** - U**aacucag**UCAGAUCACAGUC
      **5p*** - cagugguauggcucgaguugua
- **Gene Clusters:**
  - No gene clusters found

- # Mir-2689 family:
  - **Annelid-specific** family
  - **Paralogues:**
    - **Cte** has **1 copy**
    - **Efe** has **2 copies**
      - **P1_3p**
      - **P2_3p**
  - **Seeds:**
    - **3p_seed AUCCUG** is the only seed.
  - **Gene Clusters:**
    - No known conserve gene clusters

- **In Pdu:**

- **Pdu-Mir-2689_3p -** U**auccugg**CCUGCAAGUGCACA

- **5p* -** ugcauucgcaugucuggaugca

- **Gene clusters:**
  - No gene clusters found

- # Mir-2691 Family:

- **Annelid-specific** family
- **Orthologues and Paralogues**:
    - **Cte** has 1 copy - **Mir-2691_3p**
    - **Efe** has 4 copies
        - **P1_3p**
        - **P2_3p**
        - **P3_3p**
        - **P4_3p**
- **Seeds:**
    - **3p-mature -** UUUGCAA
- **Gene Clusters:**
    - Unknown

- **Pdu**
    - Pdu appears to have maybe 5 copies, temporary called:
        - **Pdu-Mir-2691-1_3p** - A**uuugcaa**UGUAUCACUGCCCG
            - **5p*** - ggcgaugucacauugcagucg

- 5p* has no significant allignments
- However, in the 2020SmallRNASeq is not found. but only predicted

```
> Efe-Mir-2691-P4_3p
Length=22

 Score = 28.8 bits (15),  Expect = 0.010
 Identities = 15/15 (100%), Gaps = 0/15 (0%)
 Strand=Plus/Plus

Query  2   TTTGCAATGTATCAC  16
           |||||||||||||||
Sbjct  2   TTTGCAATGTATCAC  16
```

- **Pdu-Mir-2691-2_3p** (A**uuugcaa**UGAAUCACUGCCCA)
    - **5p* -** gguggcgauucuuugcagucg

- 5p* has no significant allignments
- However, in the 2020SmallRNASeq is not found. but only predicted

```
> Efe-Mir-2691-P4_3p
Length=22

 Score = 23.3 bits (12),  Expect = 0.46
 Identities = 14/15 (93%), Gaps = 0/15 (0%)
 Strand=Plus/Plus

Query  2   TTTGCAATGAATCAC  16
           ||||||||| |||||
Sbjct  2   TTTGCAATGTATCAC  16

> Efe-Mir-2691-P3_3p
Length=21

 Score = 23.3 bits (12),  Expect = 0.46
 Identities = 16/18 (89%), Gaps = 0/18 (0%)
 Strand=Plus/Plus
```

```
Query  2   TTTGCAATGAATCACTGC  19
           ||||||| | ||||||||
Sbjct  2   TTTGCAAAGTATCACTGC  19
```

- **Pdu-Mir-2691-3_3p** (A**uuuugca**UCGUAUCACUGCCGG)
    - **5p* -** g**guagcga**uucccugacaaaaucu

- This paralogue presents a **seed-shift** in **3p-direction**. Or maybe caused by the insertion of a U.
- 5p* has no significant allignments

```
> Efe-Mir-2691-P3_3p
Length=21

 Score = 25.1 bits (13),  Expect = 0.14
 Identities = 17/19 (89%), Gaps = 0/19 (0%)
 Strand=Plus/Plus

Query  2   TTTTGCATCGTATCACTGC  20
           |||||||  ||||||||||
Sbjct  1   TTTTGCAAAGTATCACTGC  19
```

- **Pdu-Mir-2691-4_3p** (G**uuuugca**UGGUAUCACUGCUC)
    - **5p* -** guagugauacuuugcacaaaucg

- This paralogue shows good alignment only with Efe-Mir-2691-P3_3p

```
> Efe-Mir-2691-P3_3p
Length=21

 Score = 27.0 bits (14),  Expect = 0.035
 Identities = 18/20 (90%), Gaps = 0/20 (0%)
 Strand=Plus/Plus

Query  2   TTTTGCATGGTATCACTGCT  21
           |||||||  |||||||||||
Sbjct  1   TTTTGCAAAGTATCACTGCT  20
```

5p* shows good Plus/Minus Alignment with Efe-Mir-2691-P3_3p

```
> Efe-Mir-2691-P3_3p
Length=21

 Score = 27.0 bits (14),  Expect = 0.040
 Identities = 17/18 (94%), Gaps = 1/18 (6%)
 Strand=Plus/Minus

Query  3   AGTGATACTTTGCACAAA  20
           |||||||||||||| |||
Sbjct  17  AGTGATACTTTGCA-AAA  1
```

- Note: 2 copies of this gene, with identical Heirpin sequences are found with the 2019SmallRNA-2022MirDeep2-2022Genome:
    - scaffold_44:156337..156395:+
    - scaffold_297:185454..185512:-

- However, scaffold 297 is quite small and has a lot of repetitive elements, so this second copy might be an artifact.

- **Pdu-Mir-2691-5_5p** (G**acgguga**AACUUGUGCAAAACC)
    - **3p*** - uuuugcaaaguaucacagcuua
      uuuugcaaaguaucacagcuua
- This paralogue shows good alignment only with Cte-Mir-2691_5p*.
- **Arm-shift** from 3p to **5p-mature** arm.
    - This is the only case known in this family.

```
> Cte-Mir-2691_5p*
Length=23

 Score = 27.0 bits (14),  Expect = 0.040
 Identities = 18/20 (90%), Gaps = 0/20 (0%)
 Strand=Plus/Plus

Query  1   GACGGTGAAACTTGTGCAAA  20
           |||||||| |||| ||||||
Sbjct  1   GACGGTGAGACTTCTGCAAA  20
```

- The 3p* shows great alignment with 3p of the other annelids

```
> Cte-Mir-2691_3p
Length=23

 Score = 36.2 bits (19),  Expect = 6e-05
 Identities = 19/19 (100%), Gaps = 0/19 (0%)
 Strand=Plus/Plus

Query  1   TTTTGCAAAGTATCACAGC  19
           |||||||||||||||||||
Sbjct  1   TTTTGCAAAGTATCACAGC  19
```

- Paralogy with Efe paralogues is not clear. Probably are independent duplications.

- **Gene Clusters:**
    - No gene clusters found

- # Mir-2692 Family:
    - **Annelid-specific** family
    - **Paralogues:**
        - **Only Cte** has 1 copy
        - **Efe** does not have it. Maybe lost it.
    - **Seeds:**
        - **3p_seed CAGUCAA** is the only one
    - **Gene Clusters:**
        - In Cluster with lophotrocozoan-specific miRNA Mir-1990 paralogues and Mir-1986
            - **In Cte:** Mir-1990-P2_<150ntds_Mir-2692_~6000ntds_Mir-1990-P1_~1000ntds_Mir-1986

- **In Pdu:**
    - **1 copy** is present
    - **Pdu-Mir-2692_3p** - C**cagucaa**UGUUGACACCACCGCU
        - **5p* -** ggugaugucccauugacuuuggug

- **3p_mature** has complete allignment only with Cte-mir-2692
- 
- **Gene Clusters:**
    - Conserved Lophotrocozoan-specific **gene cluster:**
        - Mir-1986_~1500ntds_Mir-1990-(+/-)_~1500ntds_Mir-2692

- # Mir-2705 Family:
    - **Annelid-specific** family
    - **Paralogues:**
        - **1 copy** present in **Cte** and **Efe**
    - **Seeds:**
        - **3p_mature GCAAGGU** is the only seed used

- **Gene Clusters:**
    - In Cte in Cluster with another **annelid-specific** Mir-1987
        - Mir-2705_~1000ntds_Mir-1987

- **In Pdu::**
  **1 or maybe 2 copy** of the gene:

- **Pdu-Mir-2705-1_3p** - U**cugcaag**GUAAGUGCUGUCU
    - **5p* -** gcagugcuuaccuuugcagcug

- **Almost Complete identity** of the 3p_mature with Cte
- **5p*_star** sequence is **not conserved**

- **Pdu-Mir-2705-2_3p** - U**cugcaac**GUAAGUGCUGUCA
    - 5p* - acagcgcuaugccuuuauugcaacc

- **Almost Complete identity** of the **3p_mature** with Cte and Efe
- In the 2019SmallRNAseq-2022MirDeep2-2022Genome the 3p_mature read count are very low (<300) from all the samples together. The 5p_star read is only predicted

```
> Cte-Mir-2705_3p
Length=23

 Score = 32.5 bits (17),  Expect = 0.001
 Identities = 20/21 (95%), Gaps = 1/21 (5%)
 Strand=Plus/Plus

Query  1   TCTGCAAGGT-AAGTGCTGTC  20
           |||||||||| ||||||||||
Sbjct  1   TCTGCAAGGTAAAGTGCTGTC  21

> Efe-Mir-2705_3p
Length=23

 Score = 27.0 bits (14),  Expect = 0.046
 Identities = 19/21 (90%), Gaps = 1/21 (5%)
 Strand=Plus/Plus

Query  1   TCTGCAAGGTAA-GTGCTGTC  20
           ||||||||| || ||||||||
Sbjct  1   TCTGCAAGGCAAAGTGCTGTC  21
```

- **Gene Clusters:**
    - **Annelid-specific** cluster with Mir-1987. Conserved in Cte
        - Mir-1987_~4500ntds_Mir-2705-1

- ## Mir-2707 Family:
    - **Annelid-specific** family
    - **Paralogues:**
        - **Cte** has just 1 copy - **Cte-Mir-2707_3p**
        - **Efe** has 1 copy, but both strands are active:
            - **v1_5p**
            - **v2_3p**
    - **Seeds**
        - **3p-mature** UACUUAU
        - **5p-mature** GUCAAGC
    - **Gene Cluster:**
        - No conserved gene clusters found
    -
    - **Pdu:**
        - **Pdu-Mir-2707_3p-5p -** A**auacuua**UUUUGCUUCUGACAGA
            - **5p** - u**gucaagc**agauuaaguauugu

      5p-arm is also conserved and shows great allignments with the other annelids

```
> Cte-Mir-2707_3p
Length=24

 Score = 28.8 bits (15),  Expect = 0.011
 Identities = 20/22 (91%), Gaps = 1/22 (5%)
 Strand=Plus/Plus

Query  2   ATACTTATTT-GCTTCTGACAG  22
           |||||||||  |||||||||||
Sbjct  1   ATACTTATTCAGCTTCTGACAG  22
```

- **Gene Clusters:**
    - No known gene clusters found

### Gene Clusters:

- ## Bilaterians:
    - ☑ **Mir-10-P2_Let-7_Mir-10-P3**
        - **In Cte:** Mir-10-P2_Let-7_Mir-10-P3

      **In Pdu:** Mir-10-P2_Let-7_Mir-10-P3 - within ~3000ntds

- ☑ **Mir-10** (Not a real cluster but on the same scaffold):
    - All the other Mir-10 genes sit on the same scaffold, within 1,6x10^6 ntds.

      Pdu-Mir-10-P6_Pdu-Mir-10-P5_Pdu-Mir-10-P7__Pdu-Mir-10-P1_Pdu-Mir-10-P4-2_Pdu-Mir-10-P4-1

- ☑ **Mir-1_Mir-133**

    - **In Cte:** Mir-133_Mir-1 (within 6000ntds)
    - **In Pdu:** Mir-1-3p(Plus and Minus strand)_Mir-133 (within 17.000 ntds)

- ☑ **Mir-22-P1_Mir-22-P2:**
    - The Cluster is present in Lophotrochozoans (Cgi, Lan, Lgi) but looks likes all the other lineages, lost one of the paralogues.
    - **in Cte:** Mir-22-P1_Mir-22-P2 within 500 ntds
    - **in Pdu:** Mir-22-P1_Mir-22-P2 within 1000ntds

- ☑ **Mir-252-Paralogues_Mir-2001**
    - **Bilaterian-conserved** cluster: Mir-252-Paralogues_Mir-2001
    - Note: in the new Genome assembly v2.1 (2022), Mir-2001 is not present anymore.
    - **In Cte:** Mir-2001_~2000ntds_Mir-252-P2_~1000ntds_Mir-252-P1
    - **In Pdu:** Mir-2001_~500ntds_Mir-252-P2_~1000ntds_Mir-252-P1

- ☑ **Mir-29-P1_Mir-29-P2**
    - **In Cte:** Mir-29-P1_~800ntds_Mir-29-P2
    - **In Pdu:** Mir-29-P1_~11.500ntds_Mir-29-P2

- **Protostomes:**
    - ☑ **Mir-750_Mir-1175**
        - **In Cte:** Mir-750_Mir-1175 (within 1200 ntds)
        - **in Pdu:** Mir-750_Mir-1175 (within 2050 ntds)

- ☑ **Mir-12_Mir-216:**
    - **In Cte:** is organized this way **Mir-12_Mir-216-P1d_Mir-216-P2_Mir-216-P1c**
    - **In Pdu: Mir-12_Mir-216-P1_Mir-216-P2** within ~ 3200 ntds

- ☑ **Mir-34_Mir-277-Mir-317:**
    - **In Cte:** Mir-34_~400ntds_Mi-277-P2_~100ntds_Mi-277-P1_~150ntds_Mir-317
    - **In Pdu:** Mir-317_~3500ntds_Mir-277_~26.000ntds_Mir-34
        - Note: In Pdu there are more then 26.000ntds between Mir-34 and the other 2 genes.

- ☑ **Mir-2_Mir-71 - Mir-2:**
  **In Cte:**
- **Cluster 1 (Mir-2 + Mir-71):**
    - Contains 4 Mir-2 copies + Mir-71 within <600ntds
        - Mir-71_Mir-2-o37_Mir-2-o42_Mir-2-o38_Mir-2-o36
- **Cluster 2 (Mir-2):**
    - Is composed of 3 Mir-2 family genes:
        - Mir-2-o41_Mir-2-o43_Mir-2-o40 (within < 400ntds)
- **In Pdu:**
    - **Note:** the 2 clusters sit on the same scaffold, 10^6 ntds apart

        - **Cluster 1 (Mir-2 + Mir-71):**
            - Is composed of 5 or 6 Mir-2 family genes:
                - Pdu-Mir-2-oA1_Pdu-Mir-2-oB1_Pdu-Mir-2-oC1_Pdu-Mir-2-oD1_Pdu-Mir-2-oE1_?Pdu-Mir-2-oF1?_Pdu-Mir-71
                    - All within 2000ntds

    - **Cluster 2 (Mir-2):**
        - Is composed of 4 or 5 Mir-2 family genes.
            - Pdu-Mir-2-oA2_Pdu-Mir-2-oB2_Pdu-Mir-2-oC2_Pdu-Mir-2-oD2_Pdu-Mir-2-oE2
                - All within 1000ntds

- **(!)Mir-92 Cluster :**

  Is not clear wether this is a a conserved cluster, because of lack of Phylogenetic information (see Mir-92 annotation). However, Pdu shows a cluster of 3 Mir-92 copies as Cte.
- **In Cte:** Mir-92-o37_Mir-92-o35_Mir-92-o36 (within 1000ntds)
- **In Pdu:** Pdu-Mir-92-1_Pdu-Mir-92-2_Pdu-Mir-92-3 (all within <2100ntds)

- ## Lophotrochozoans:
    - **Mir-1990_(2692)_1986**

      In Annelids includes also Mir-2692
- **In Cte:** Mir-1990-P2_<150ntds_Mir-2692_~6000ntds_Mir-1990-P1_~1000ntds_Mir-1986
- **In Pdu:** Mir-1986_~1500ntds_Mir-1990-(+/-)_~1500ntds_Mir-2692
- **Mir-36_Mir-279**:
    - **Gene Cluster:**
        - **Cte:** Mir-36_Mir-279-o11 within ~2000 ntds
        - **In pdu:** Mir-36_Mir-279 - within ~4800ntds
- **Mir-96_96_315:**

  **Cte:** Mir-96-P2_Mir-315
  Lgi: Mir-96-P2_Mir-96-P3
  Cgi: Mir-96-P3_Mir-96-P2_315
  Lan: Mir-96-P2_Mir-96-P3

  **Pdu:**
  A pdu specific Mir-96-P1 cluster that might not be related to Lophotrochozoans, containing:
- Pdu-Mir-96-P1(bx or dx)_Pdu-Mir-96-P1(by or dy) - (within 1800ntds)

- ## Annelids:
    - **Mir-2705_Mir-1987**

        - **In Cte:** Mir-2705_Mir-1987 (within ~1000 ntds)
        - **In Pdu:** Mir-1987_Mir-2705 (within ~4500 ntds)

- ☑ **Mir-210 Cluster**
    - **In Cte:** Mir-210-P5_Mir-210-P6 (within ~5000 ntds)
    - **In Pdu:** 3 copies present in the same scaffold. 2 of them form a cluster within ~5800 ntds. the other one is >100.000ntds far fro the cluster
        - Mir-210-2_~5.800ntds_Mir-210-3
        - Mir-210-1_>100.000ntds_Mir-210-2_~5.800ntds_Mir-210-3

- ☑ **Mir-2000_Mir_1998:**
    - **In Cte:** Mir-2000_~200ntds_Mir-1998.
      **In Pdu:** Mir-2000_~6000ntds_Mir-1998.

- ## Pdu:
    - ☑ **Bantam-PX_Bantam-PY**
        - Only in Pdu

- ☑ **Mir-1989-genes Cluster**:
    - Only in Pdu:
      Mir-1989-1-a_~200ntds_2-a_~300ntds_3-a_~300ntds_4-b_~400ntds_5-c_~450ntds_6-d

- ☑ **Mir-1996-1 and 2**:
    - Only in Pdu: Pdu-Mir-1996-1_Pdu-Mir-1996-2 (within ~800ntds)

- ☑ **Mir-242-Paralogues Cluster:**
    - Pdu-Mir-242-(1a)_~300ntds_Pdu-Mir-242-(2b)_~300ntds_Pdu-Mir-242-(3c)_~9500_Pdu-Mir-242-(4a)

- ☑ **Mir-1992-v1/v2 - Overlapping Hairpins**

  **I Hairpin ------------------------------------------------------------------------------------**
  **cucagcacuaauccaucugug**cgucuuccugacucuggcacuggcagca**cauuaguggauaacggcuggua**guaacauuacgucugccua**ucagcaguuguaccacugaugug**

  II Haripin---------------------------------------------------------------------------------
