## Supplementary figures and images for "A genome resource for the marine annelid *Platynereis dumerilii*"

### Supplementary Figure 2

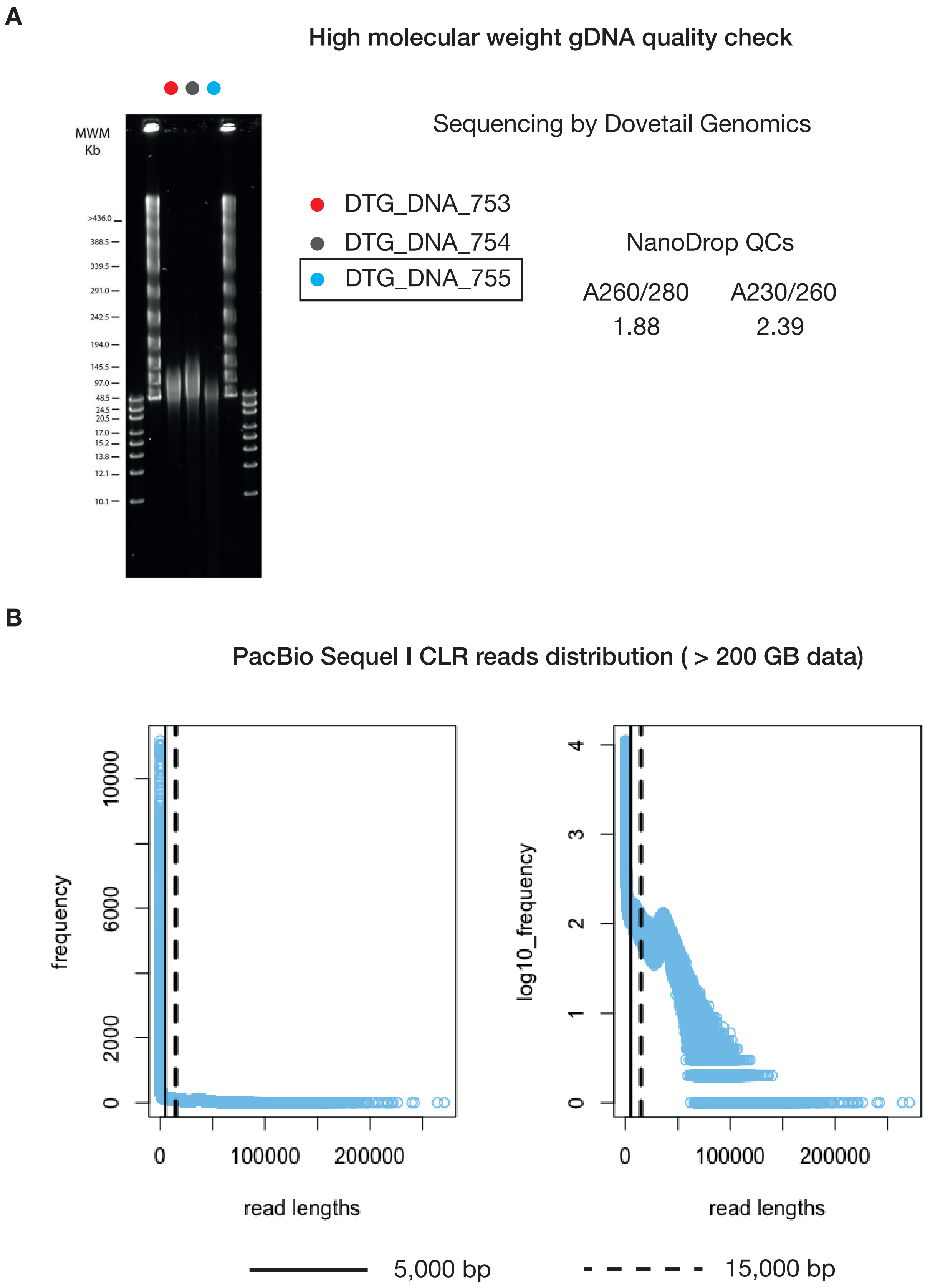

### Supplementary Figure 3

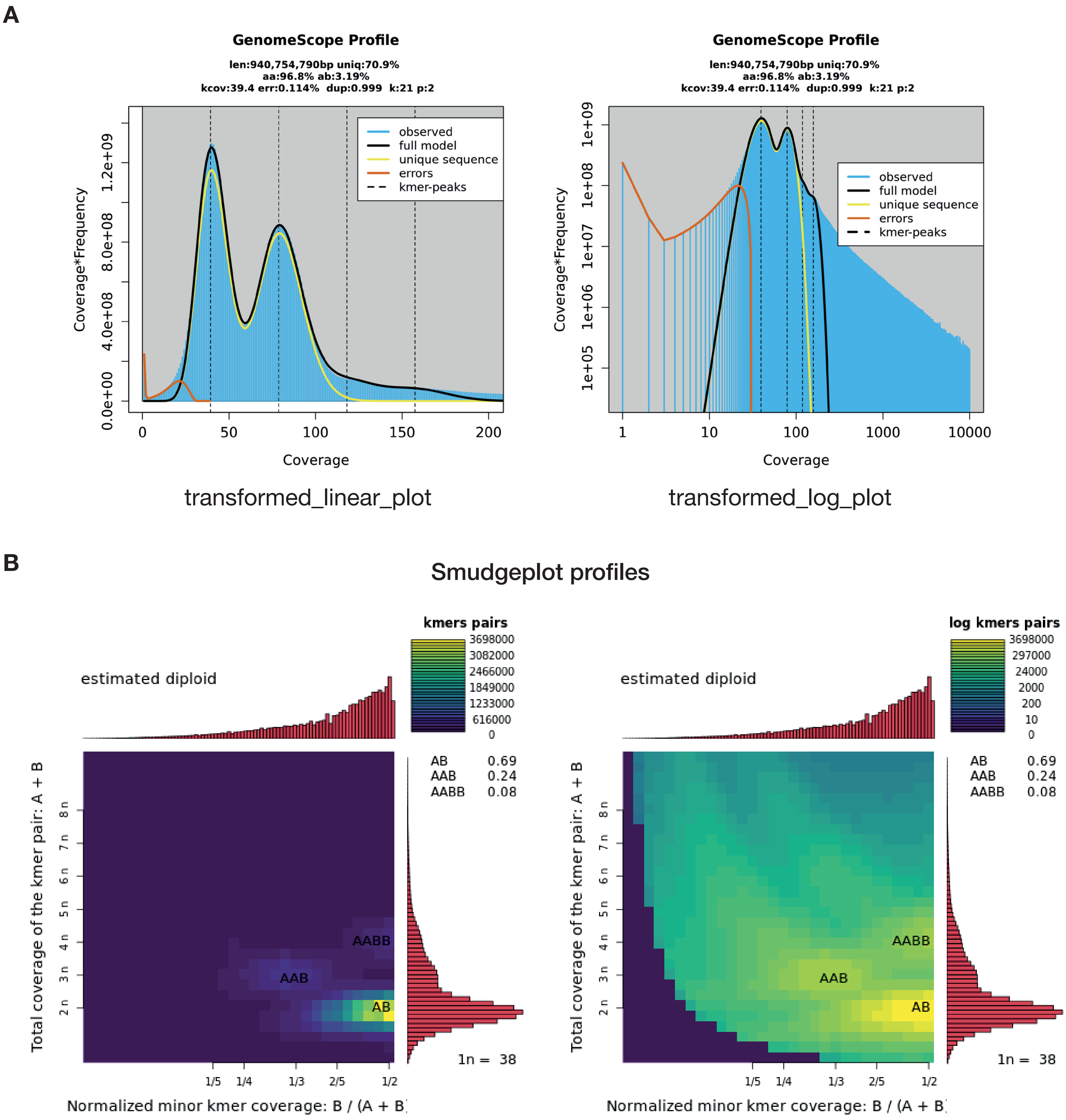

### Supplementary Figure 5

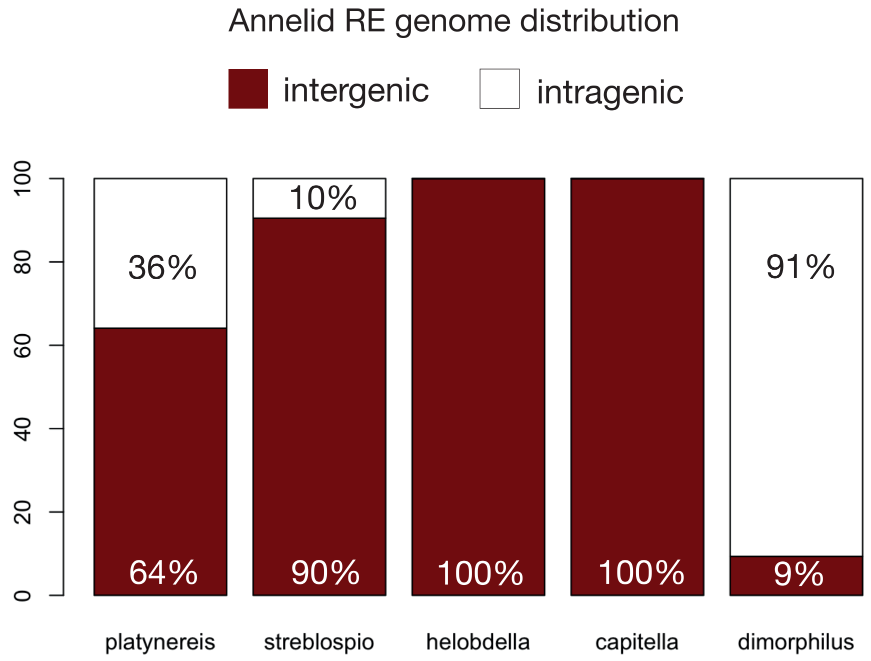

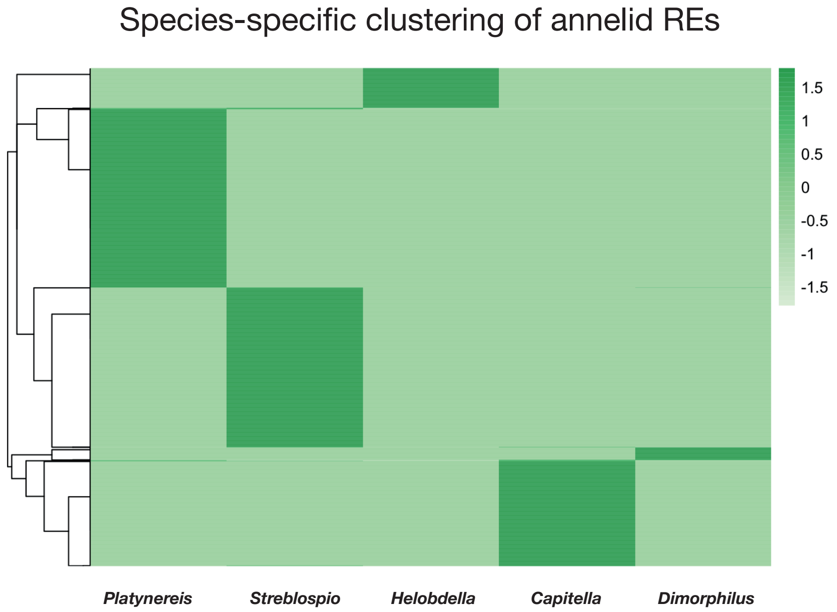

### Supplementary Figure 6

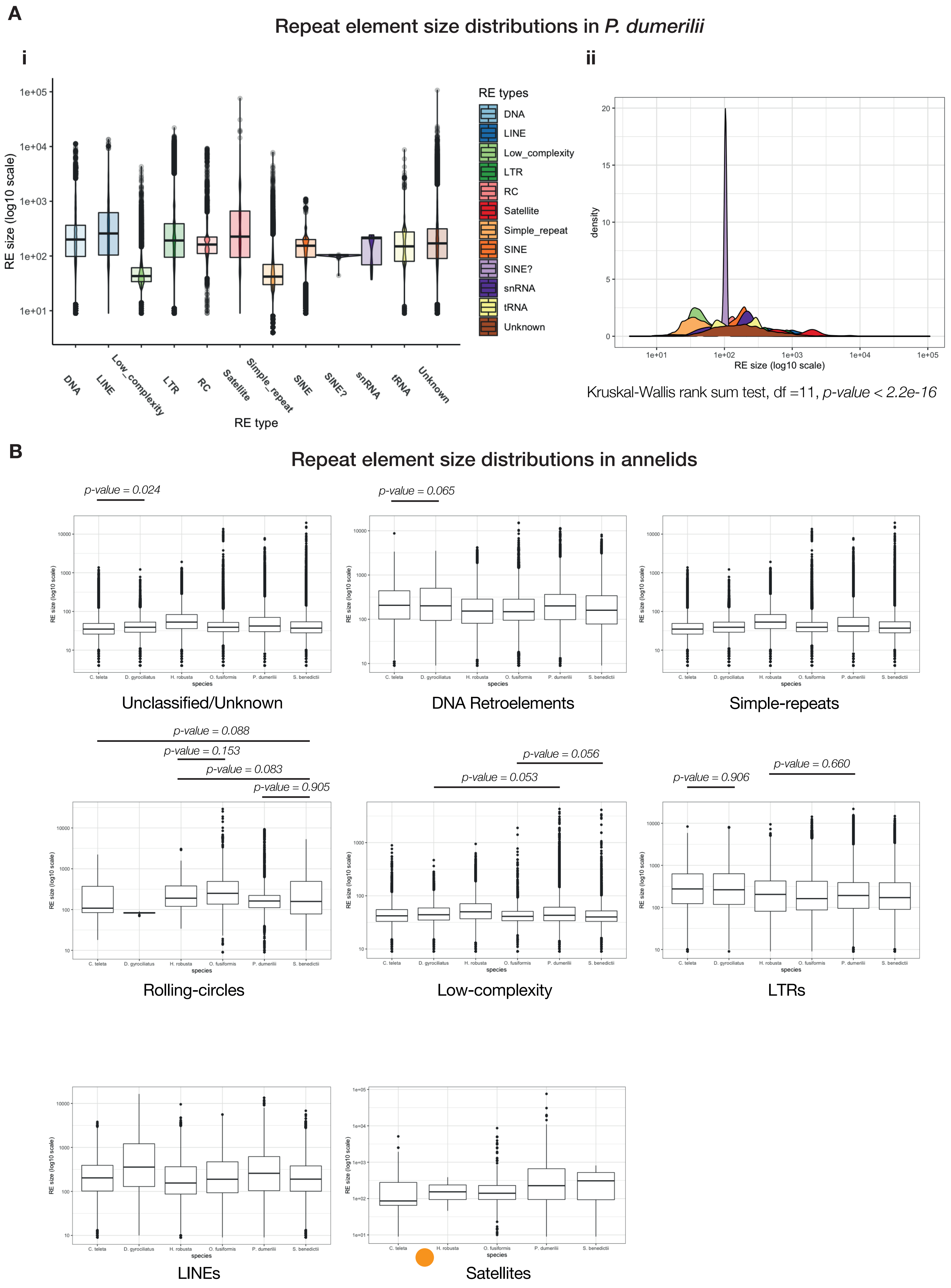

### Supplementary Figure 7

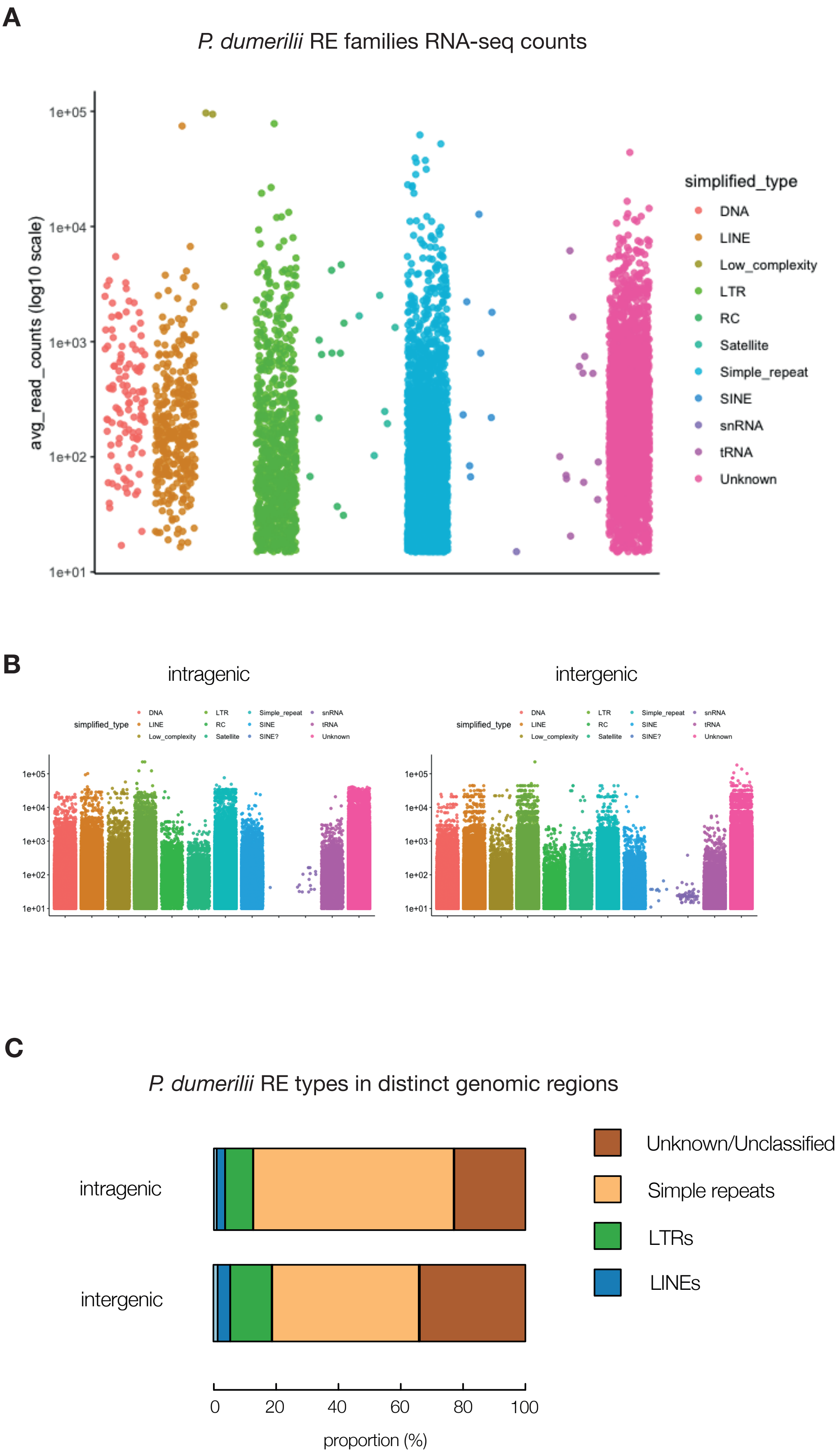


***distinct RE families only, redundant family-type pairs filtered out*

### Supplementary Figure 8

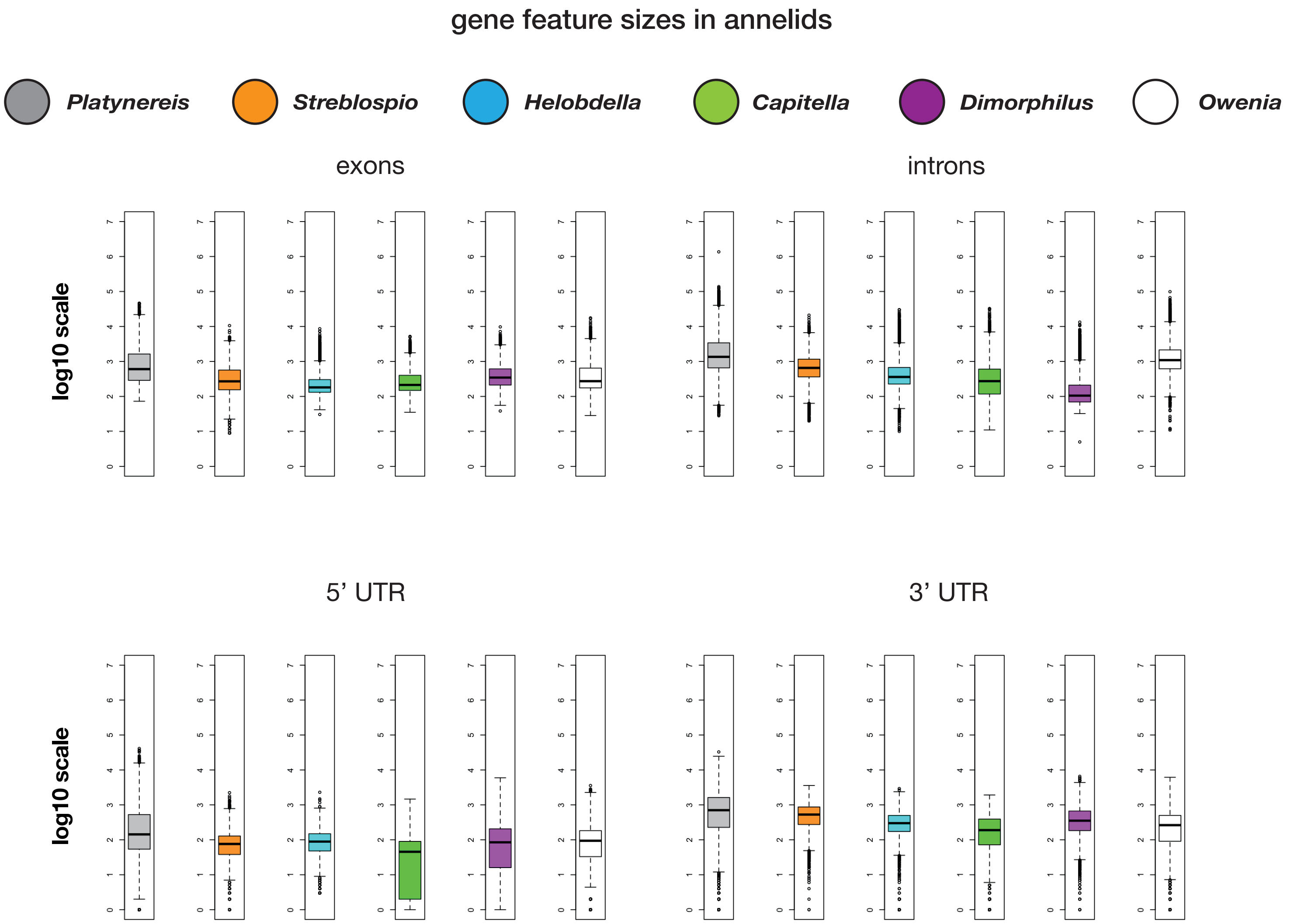

### Supplementary Figure 9

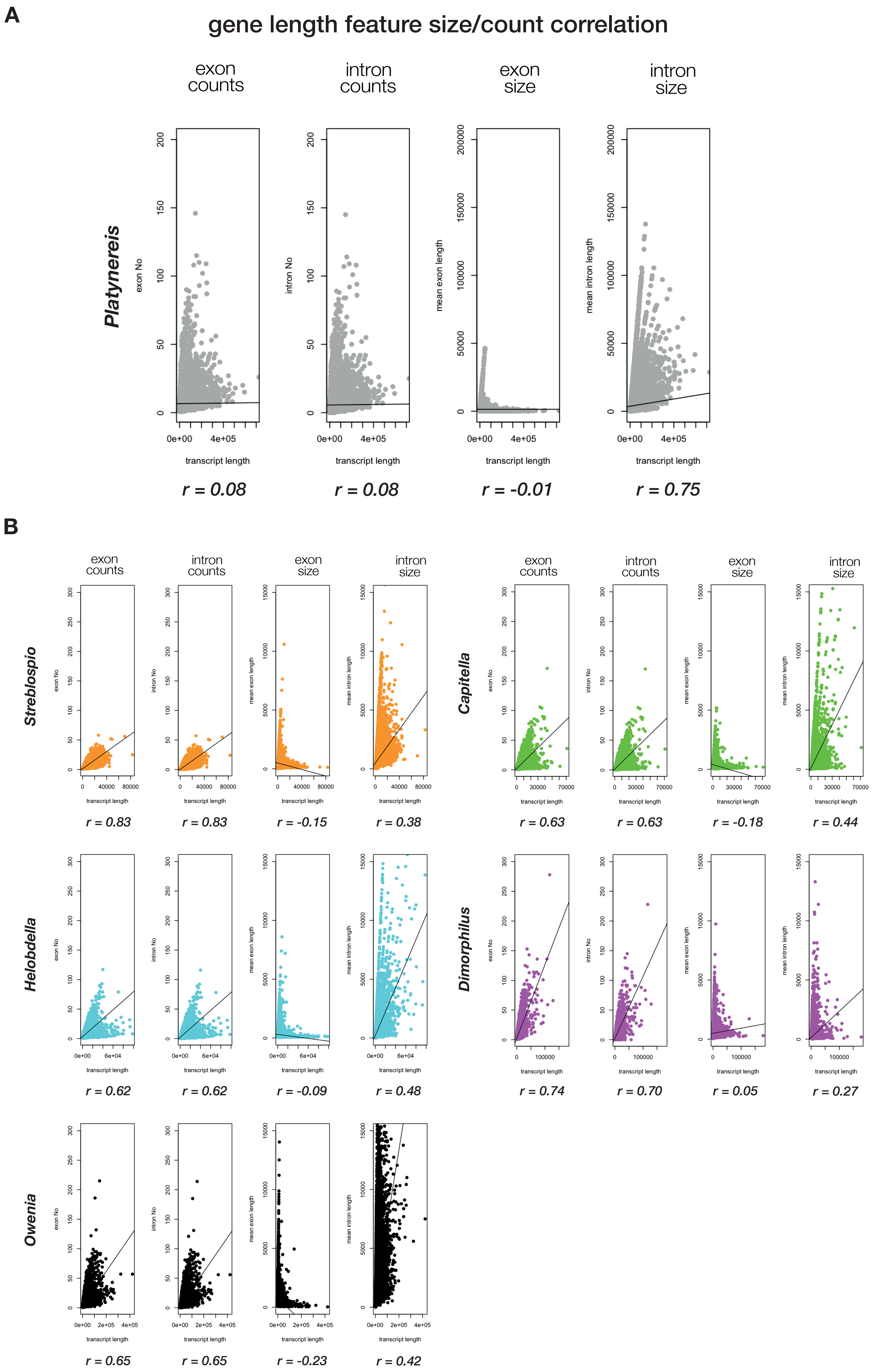

### Supplementary Figure 10

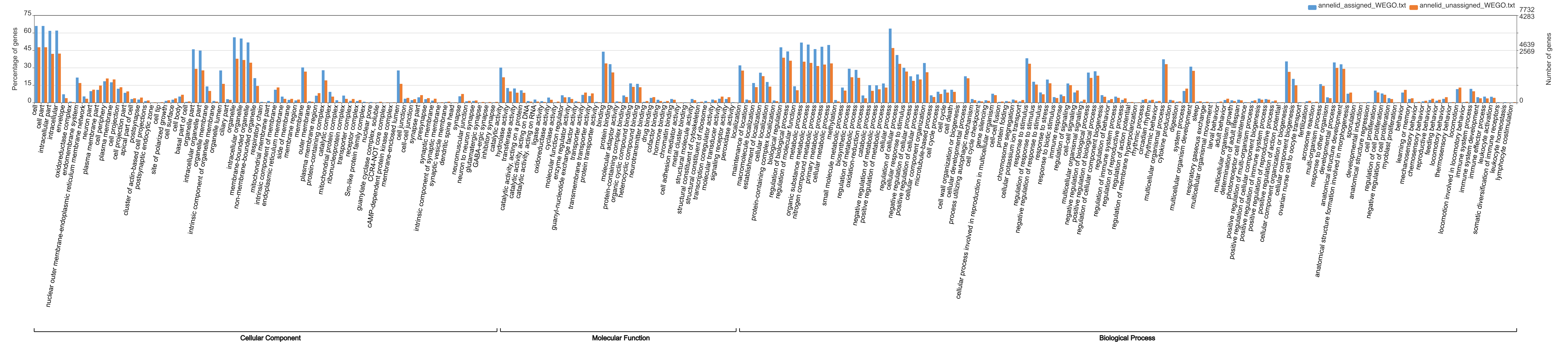

### Supplementary Figure 11

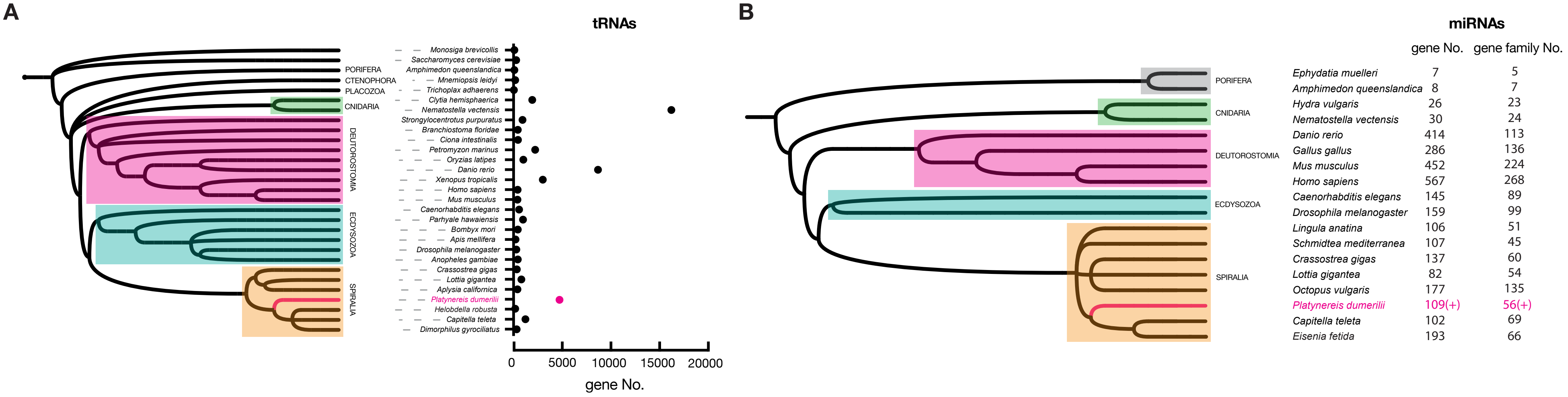

### Supplementary Figure 12

#### SNPs + In/Del(s) count per gene correlation

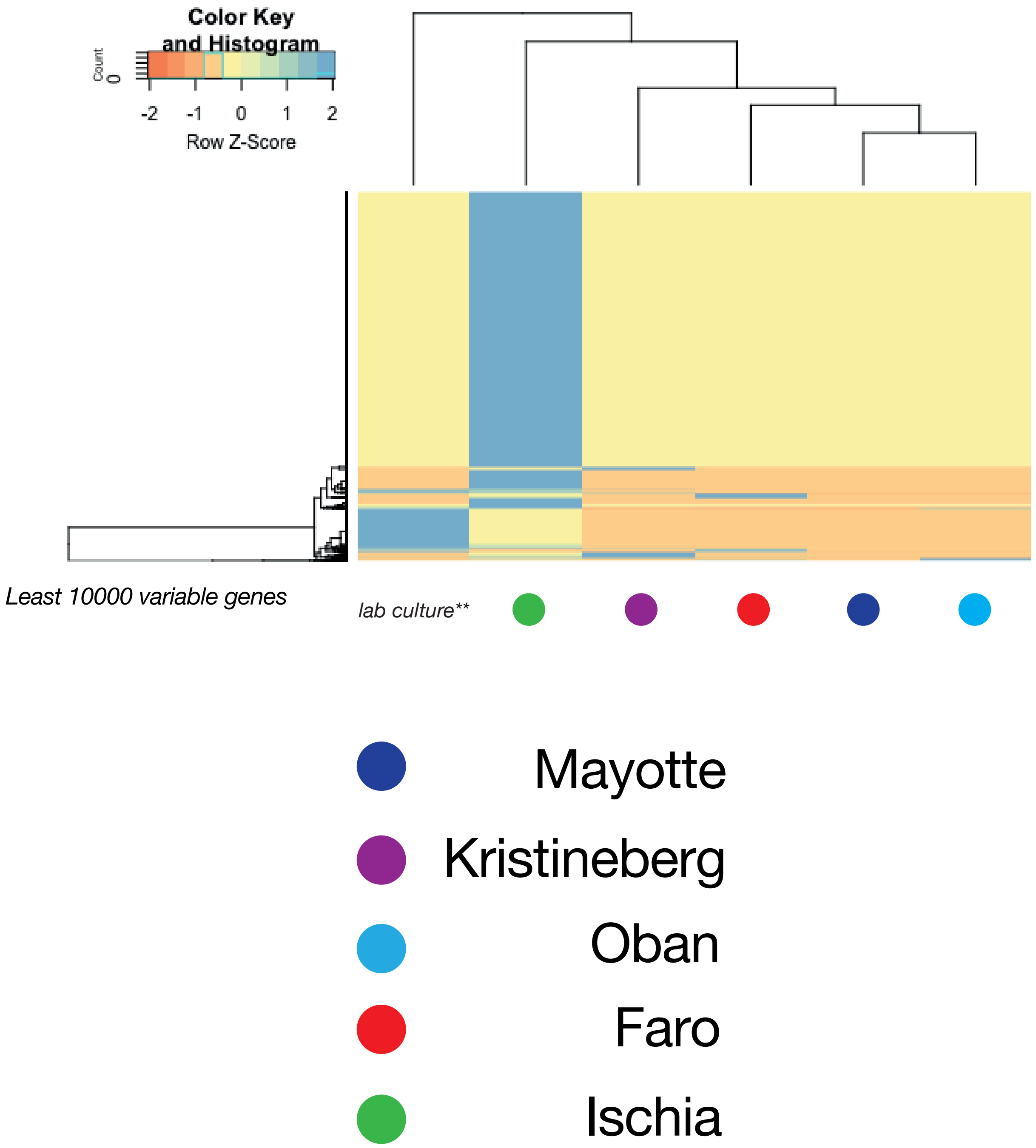

### Supplementary Figure 13

#### SNP/InDel gene feature abundance

Ischia

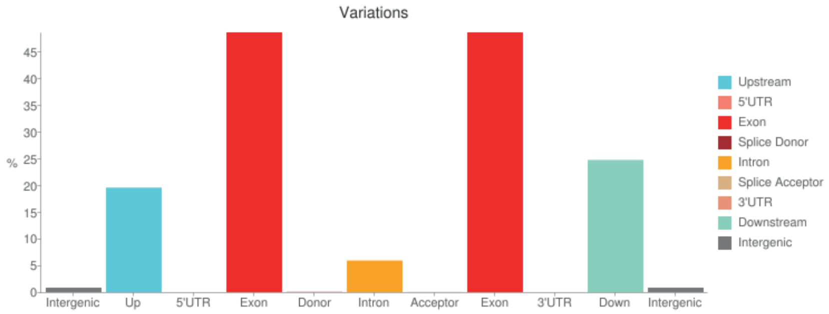


Faro

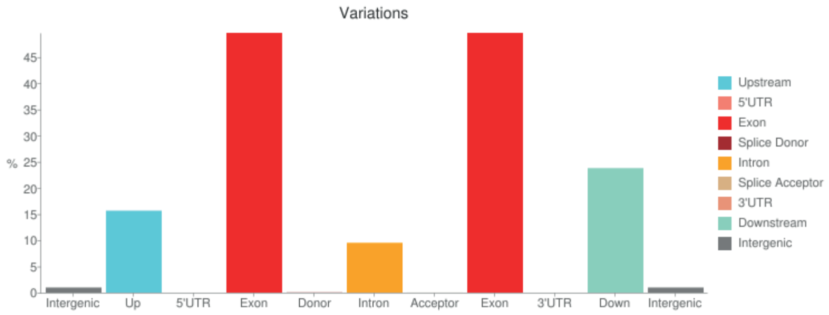


Oban

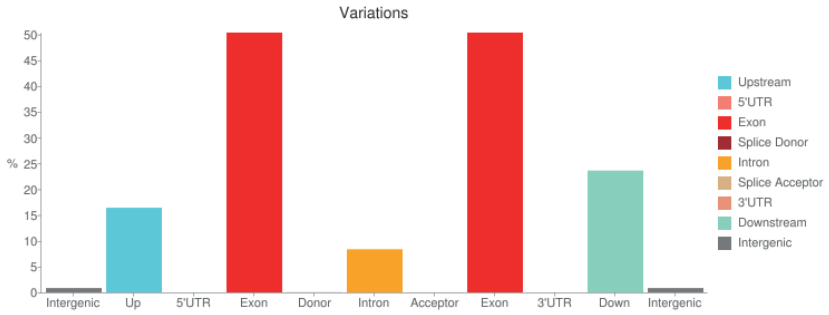


Mayotte

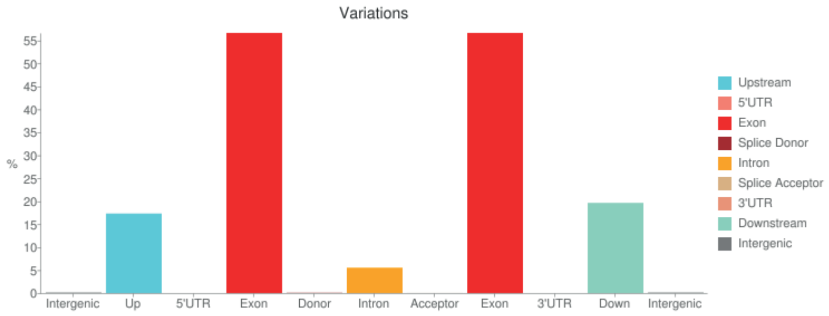

### Supplementary Figure 14

#### *P. dumerilii* XLOC_000415

scaffold_1

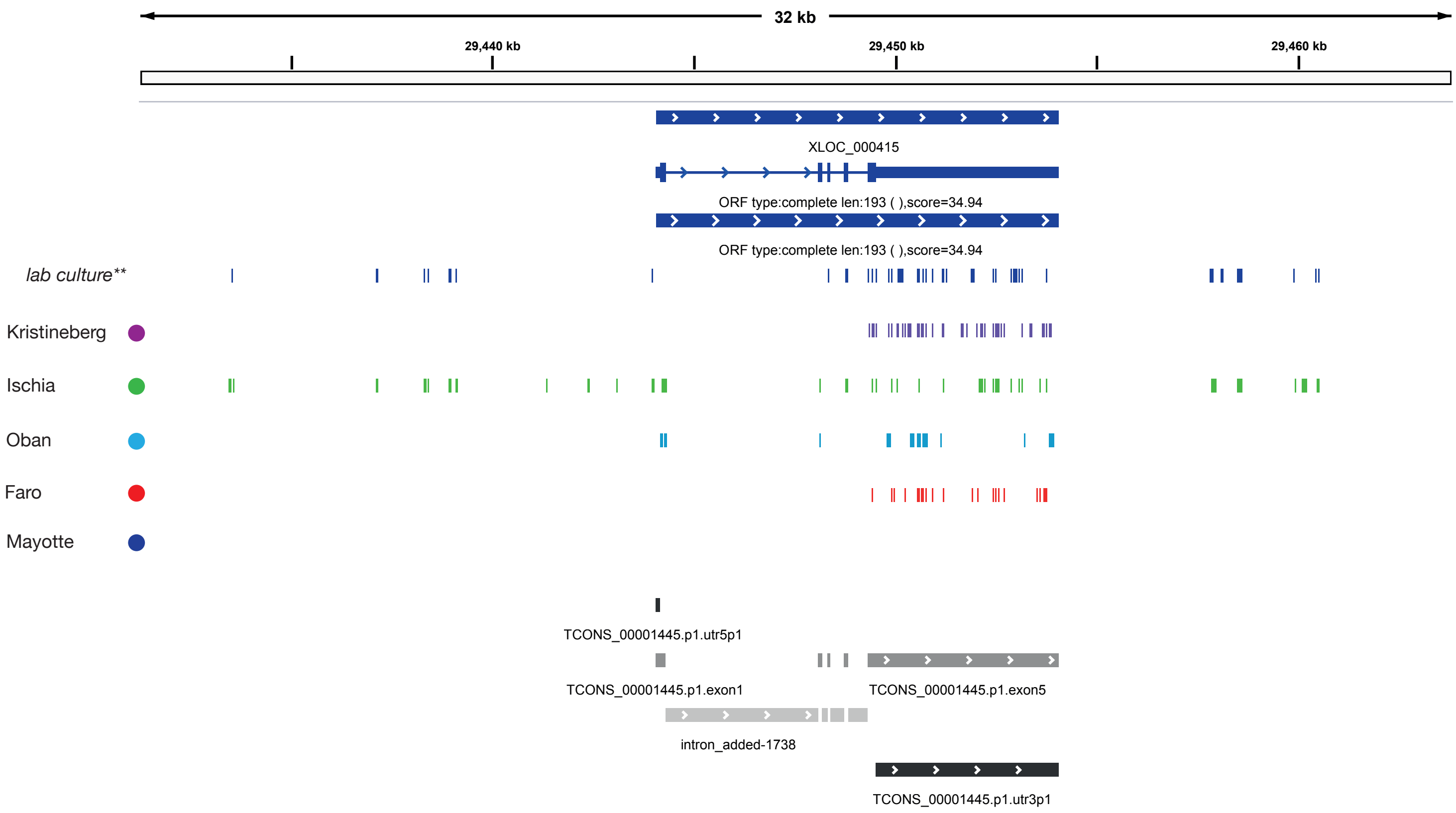

### Supplementary Figure 15

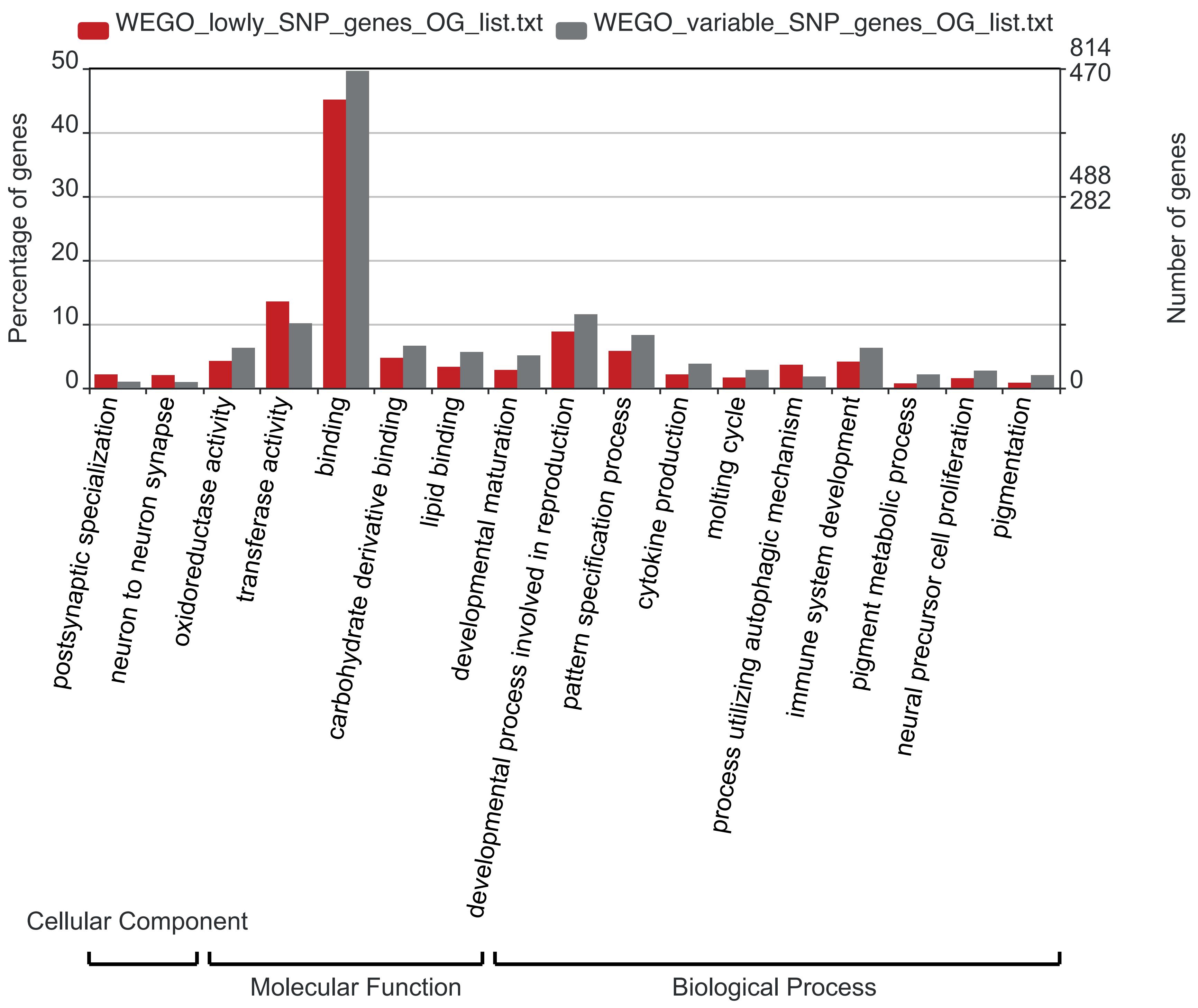
