## Supplementary Figure 4 for "A genome resource for the marine annelid *Platynereis dumerilii*"

*P. dumerilii* self-self alignment map

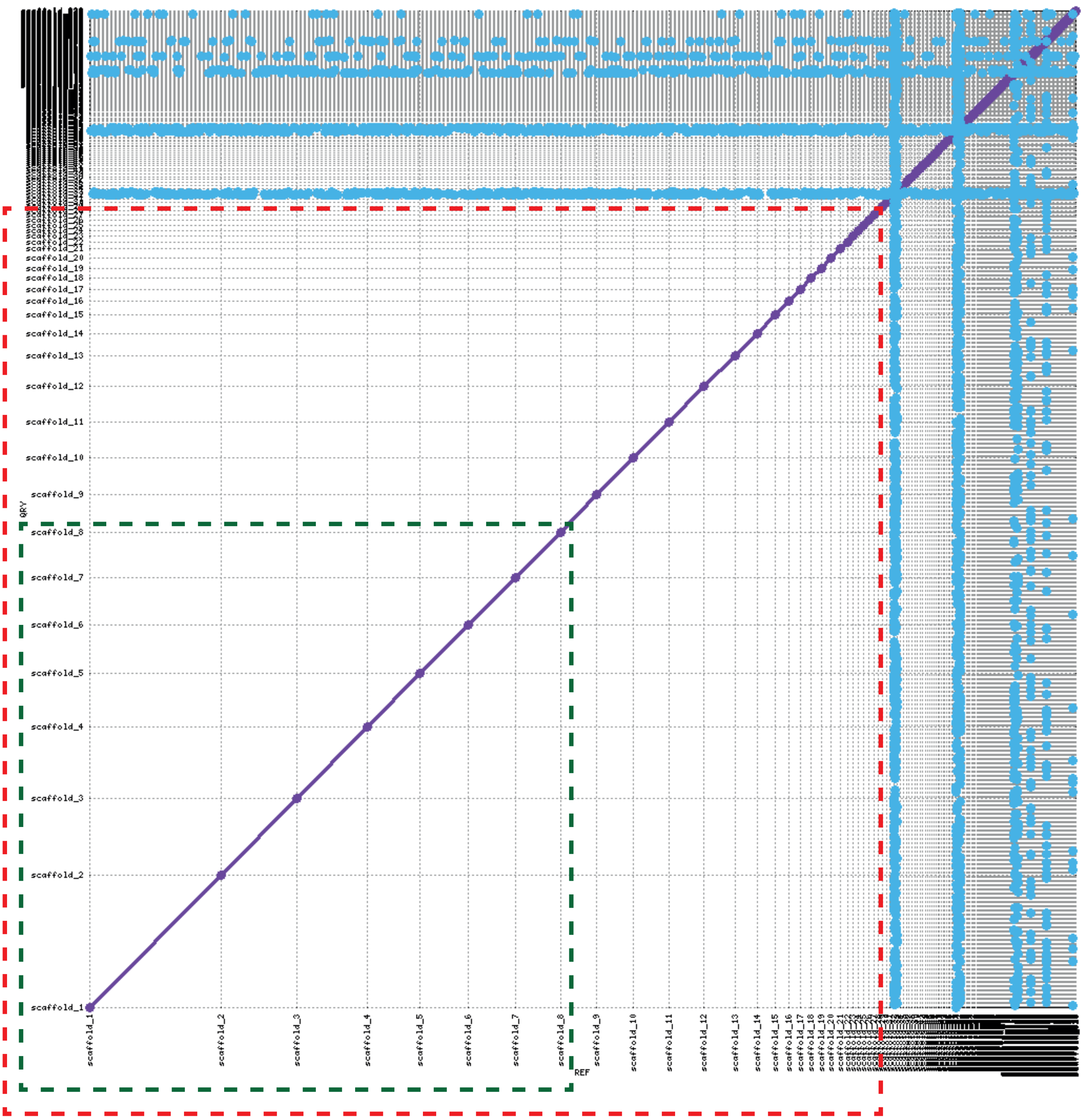


50% of assembly in 8 scaffolds

80% of assembly in 28 scaffolds

Probable repeat elements/centromeric and/or telomeric genomic sequences
