## Supplementary Figure 1 for "A genome resource for the marine annelid *Platynereis dumerilii*"

### General sequencing, assembly and annotation pipeline

**Sequencing**

PacBio Sequel II CLR
Long-read-seq

> 200 Gb raw-data

**Assembly**

CANU assembly [#]

**Removal of duplicates/haplotypes**

(i) purge_dups**4_iter

purge_haplotigs**2_iter

(ii) repeat purge_dups + purge_haplotigs iteratively**4_iter

BUSCO completion assessment at each step (>95% completion)

**Scaffolding**

(i) LINKS

(ii) Hi-C Arima pipeline + SALSA2

**Illumina polishing**

2X 150bp paired-end sequencing

**Genome annotation**

(i) Mapping total RNA-seq data

(ii) Repetitive element annotation/masking

(iii) tRNA + miRNA annotation

(iv) StringTie genome annotation

(v) Variant calling

[#] *Three different assemblers were tried and tested prior to settling on the CANU algorithm*
